# Host Endoplasmic Reticulum Stress and Interferon Responses Contribute to AAV-Induced Ocular Toxicity

**DOI:** 10.64898/2026.01.13.698457

**Authors:** Apolonia Gardner, Christin Hong, Sophia Rong Zhao, Adam Daniels, Constance L. Cepko

## Abstract

Adeno-associated viruses (AAVs) are popular gene therapy vectors, but AAVs can cause toxicity. This is particularly evident following expression of some transgenes, e.g. GFP, in the retinal pigment epithelium (RPE), which leads to loss of RPE cells and photoreceptors. Here, we sought to unravel the toxicity mechanism(s). Several transgenes, self and non-self, were tested for toxicity. Some transgenes were toxic while others were not, with no clear correlation for toxicity with regards to self vs non-self transgenes. RPE RNA-sequencing revealed upregulation of translational processes, cell stress, cytokine release, antiviral responses, and leukocyte infiltration pathways. Toxicity-inducing pathways were explored for causality by injecting toxic AAVs into mice deficient for intrinsic, innate, or adaptive immune pathways. The CHOP KO partially alleviated toxicity for RPE but not photoreceptors, whereas the type I interferon receptor KO partially alleviated toxicity for photoreceptors but not RPE. In situ hybridization of interferon pathway transcripts (IFNB1, IFNAR1) revealed that the RPE and retina can produce and potentially respond to interferon. These data suggest that transgene-induced cell stress responses in the RPE lead to RPE cell death, while interferon signaling contributes to the death of photoreceptors.

## Introduction

The field of adeno-associated virus (AAV) gene therapy has enjoyed several clinical approvals since the first FDA-approved gene therapy in 2017. This first gene therapy, Luxturna, uses AAV to express RPE65 and mitigates retinal dystrophy due to loss-of-function mutations in the gene *RPE65*.^1–3^ AAV has since become the most widely used ocular gene therapy vector. It can be easily engineered, produced at high titer, and importantly, is relatively safe. With a few caveats, wild-type (WT) AAV has historically not been associated with any human disease even though an estimated 70% of the human population is seropositive for at least one serotype of AAV.^4^ One caveat to AAV’s nonpathogenicity surfaced a few years ago, when cases of pediatric hepatitis were linked to high liver titers of AAV2.^5–7^ In the children with hepatitis, co-infections with human adenovirus (HAdV) or human herpesvirus 6B (HHV-6B) likely served as helper viruses for WT AAV2 replication in the liver and may have contributed to the pathology.^5–7^ Recombinant AAVs (rAAVs) for gene therapy, which are designed to have all viral genes removed and replaced with therapeutic transgenes, are theoretically safer. However, rAAVs are commonly administered at much higher doses than would be encountered in the wild, particularly when delivered systemically, and AAV gene therapy has a well-documented history of toxicity at higher doses.^8–14^ The issue of AAV toxicity came to a head in 2020, when the first AAV-associated human deaths were reported in a phase 2 clinical trial run for X-linked myotubular myopathy (XLMTM).^15^ More deaths have been subsequently reported,^16–19^ such as three patient deaths from acute liver failure following systemic AAV gene delivery to treat Duchenne muscular dystrophy (DMD).^17^

Research on AAV-associated toxicity in gene therapy has focused on immune responses to AAV.^20^ These include cytotoxic T cell responses to the AAV capsid,^21–25^ TLR2-based recognition of the AAV capsid,^26^ TLR9-based responses to AAV DNA,^27–29^ MAVS-based recognition of double-stranded RNA (dsRNA),^30^ and immune responses to a given transgene.^31–33^ In addition, Hinderer et al. noted that AAV toxicity can occur independently of activation of the adaptive immune response, perhaps due to cellular stress pathways that prime cells for an exaggerated response to inflammation.^8^ Unsurprisingly, the clearest correlation between AAV and toxicity is the dose of AAV. The most severe side effects from AAV gene therapy have been from systemic delivery with high doses, which can be >1000-fold higher than those used for targeted administration.^34^

The eye is a particularly attractive target for AAV gene therapy. The eye is a relatively immune privileged site, e.g., it can support tissue grafts for an extended period of time without rejection,^35,36^ and it is normally protected from surveillance by T cells or antibodies. During the development of Luxturna, the same AAV was administered to patients in their second eye after positive results were obtained from treating their first eye. The results were positive, indicating that no or insufficient systemic neutralizing antibodies were generated from the first treatment.^37^ Despite encouraging results from Luxturna, AAV can still cause ocular toxicity. Chorioretinal atrophy of the RPE, occurring in the area around the subretinal Luxturna injection bleb, has been reported to occur within the first year post-treatment.^38,39^ Our lab has also observed disruption of the RPE and retina after subretinal injection of various AAVs in mice.^12^

In this study, we evaluated potential mechanisms that may contribute to ocular toxicity. We compared responses to a toxic vs a benign AAV. The toxic AAV expresses GFP only in the RPE using the RPE-specific Bestrophin1, or Best1, promoter (Best1::GFP). The benign AAV expresses GFP only in photoreceptors using the rod-specific Rhodopsin, or Rho, promoter (Rho::GFP). Mice with a KO of the type 1 interferon receptor (IFNAR) or the cGAS cytosolic DNA sensor had partial alleviation of photoreceptor toxicity, but not RPE toxicity. Supporting the involvement of a type I interferon response, >200 interferon-stimulated genes (ISGs) were shown to be induced following subretinal injection of a toxic AAV relative to a benign AAV. In situ hybridization demonstrated expression of type I interferons (IFNB1) and receptors (IFNAR1) from AAV-infected RPE and retina. In addition to interferon responses, inflammatory responses were also explored. Microglia, the resident immune cells of the retina, are recruited to the photoreceptor and RPE layers upon injection of toxic AAVs.^12^ They undergo pyroptosis, as we found expression of the inflammasome adaptor ASC in this condition. Pharmacologic depletion of microglia partially alleviated RPE toxicity and cone toxicity at a high AAV dose, but not ONL toxicity. Genetic deletion of CX3CR1 and CCR2 (chemokine receptors found on microglia), or genetic deletion of gasdermin D (a key mediator of pyroptosis in myeloid cells) also did not alleviate toxicity.

The expression of GFP in the RPE likely triggered some form of translational stress, as suggested by RNA-seq data as well as the CHOP KO mouse. CHOP is a downstream effector of the unfolded protein response (UPR), which is induced by endoplasmic reticulum (ER) stress.^40^ The CHOP KO mouse exhibited less RPE cell pathology, but did not improve photoreceptor health, perhaps due to the continued induction of interferon in this KO strain. Transgenes other than GFP were also tested, giving varying levels of toxicity. Some transgenes (e.g., RPE65, the Luxturna transgene) elicited no detectable toxicity at the doses and timepoints tested. Overall, the data suggest that transgenes can induce ER stress in the RPE, resulting in RPE toxicity. Interferon responses generated concurrently with this RPE insult can contribute to retinal toxicity. It is likely that each transgene and virus dose elicit distinct responses, with the data presented here providing examples of responses that can lead to AAV toxicity.

## Results

### Assessment of toxicity from AAVs expressing GFP exclusively in RPE or photoreceptors

Our initial report of AAV-induced toxicity in the eye showed that AAVs with promoters that are broadly expressed (CAG, UbiC, CMV) or specifically active in the RPE (Best1 promoter) are toxic.^12^ In contrast, AAVs with photoreceptor-specific promoters (4/4 promoters tested) were benign, at the same or higher doses, using the same capsid and preparation methods. However, this original set of AAVs harbored different genomic elements, including different introns, polyA sequences, ITRs, +/-WPRE. This made it difficult to interpret whether the only factor contributing to AAV toxicity was expression in the RPE.

To determine if transgene expression only in the RPE is sufficient for RPE pathology, the RPE-specific Best1^43,52,53^ promoter and the rod-specific Rho promoter were cloned into the same AAV backbone (i.e., the remaining vector sequences were identical). This backbone is the one used for our initial study of toxicity by the CMV::GFP AAV (Figure 1A).^12^ Both promoters were human sequences of the relevant genes. To compare the toxicity of the resulting AAV vectors, 2e9 genome copies (gc) of Rho::GFP or Best1::GFP were injected subretinally into C57BL/6J (B6J) mice at P0.

**Figure 1.**
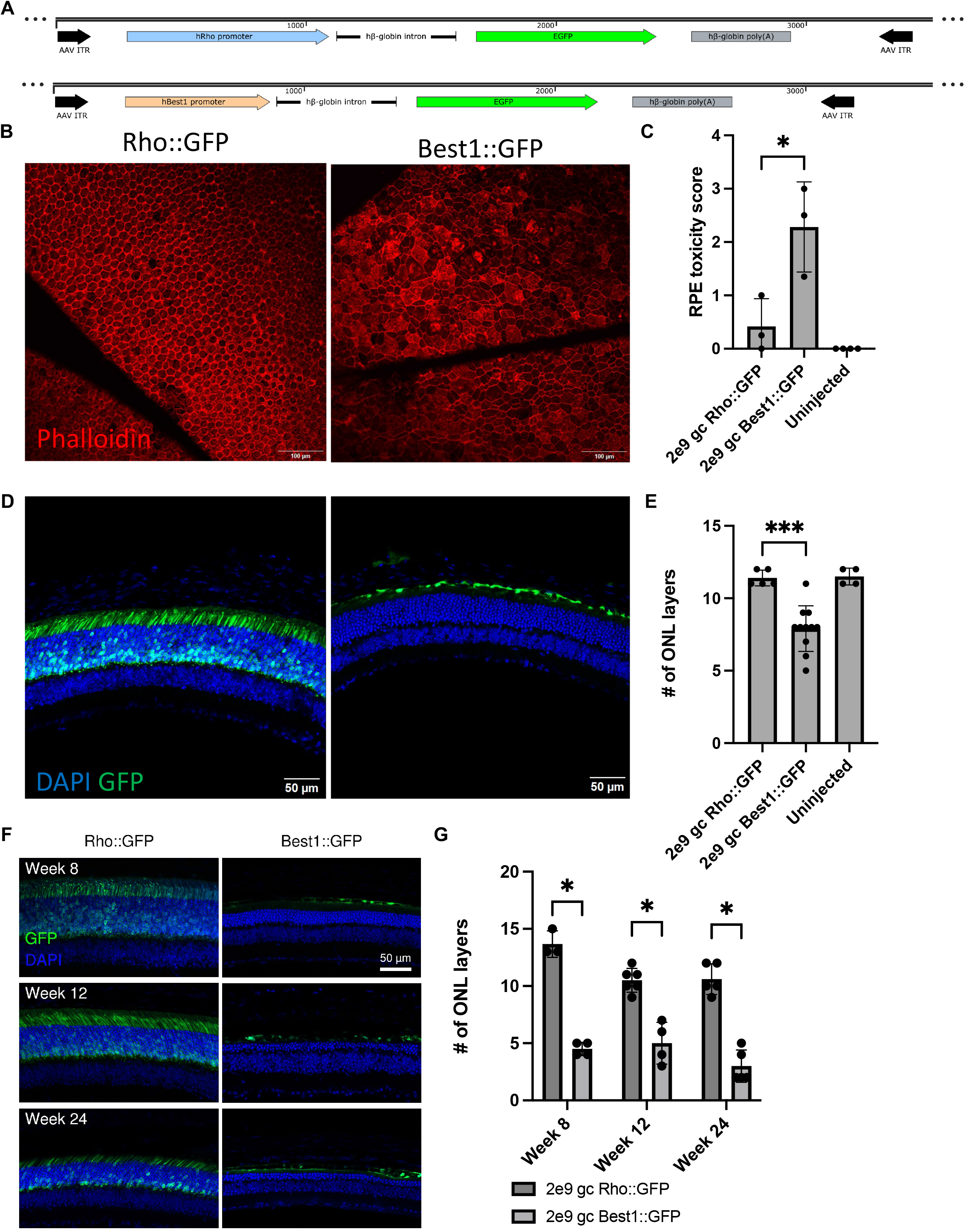
Assessment of toxicity from AAVs expressing GFP exclusively in RPE or photoreceptors. A) AAV vector maps for Rho::GFP and Best1::GFP. B) RPE morphology upon staining RPE flat mounts with phalloidin, which labels the F-actin enriched RPE cell junctions. Flat mounts were harvested 2 weeks post subretinal injection of 2e9 gc of either Rho::GFP or Best1::GFP into P0 pups. RPE score of Best1::GFP is 3. Scale bar: 100 µm. C) Semi-quantitative scoring of RPE cell toxicity from samples described in panel B (n=3-4 eyes per group, *p=0.03, mean ± SD, unpaired t-test). Uninjected P30 C57BL/6J RPE flatmounts were used as a control. D) Retinal morphology upon staining whole eyecup sections with DAPI, which labels nuclei. Eyecups were harvested 4 weeks post subretinal injection of 2e9 gc of Rho::GFP or Best1::GFP into P0 pups. Scale bar: 50 µm. E) Median number of ONL layers remaining for samples shown in panel D (n≥4 eyes per group, ***p=0.0003, mean ± SD, unpaired t-test). Uninjected 4 week C57BL/6J whole eyecup sections were used as a control. F) Representative whole eyecup sections from mice injected with 2e9 gc Rho::GFP or Best1::GFP at various times post infection ranging from 8-24 weeks. Green: GFP. Blue: DAPI-labeled nuclei. Scale bar: 50 µm. G) ONL thickness quantification over time after subretinal injection of Rho::GFP vs. Best1::GFP (n=3-6 eyes per group, *p=0.0001 for week 8, *p=0.0008 for week 12, *p<0.0001 for week 24, mean ± SD, multiple unpaired t-tests with Bonferroni-Dunn multiple comparisons correction).

To evaluate RPE pathology after administration of Rho::GFP or Best1::GFP, RPE flatmounts were collected two weeks post subretinal injection and stained with phalloidin to label the actin-rich RPE cell boundaries. Eyes injected with 2e9 gc/eye of Rho::GFP had a predominantly healthy RPE monolayer with RPE cells arranged in a regular hexagonal array (Figure 1B). In contrast, eyes injected with 2e9 gc/eye of Best1::GFP had dysmorphic RPE cells (Figure 1B). A semiquantitative scoring method for the RPE that was previously developed, based upon overall cellular morphology and regularity of the cellular distribution, revealed a marked increase in toxicity in Best1::GFP vs. Rho::GFP samples (Figure 1C and Figure S1A). Rho::GFP injected RPE flatmounts were as healthy, or nearly so, as uninjected RPE flatmounts (Figure S1A).

Retinal pathology also was evaluated after administration of Rho::GFP or Best1::GFP. Four weeks after subretinal injection, the number of ONL layers was measured from cross-sections of uninjected eyes, or eyes injected with Best1::GFP or Rho::GFP (Figure 1D-E and Figure S1B). At this timepoint, the number of ONL layers was significantly lower following injection of Best1::GFP relative to Rho::GFP and uninjected controls (Figure 1E). The number of ONL layers remaining at 4 weeks for Rho::GFP injected eyes and uninjected eyes was nearly identical (∼12 layers, Figure 1E). The number of ONL layers got progressively lower over time in samples that received Best1::GFP. At 24 weeks post-injection, Rho::GFP-treated eyes had an average of ∼10.6 ONL layers remaining, whereas Best1::GFP-treated eyes averaged ∼3 layers (Figure 1F-G).

To determine if the dose of Best1::GFP correlated with the magnitude of toxicity, B6J mice were injected with Best1::GFP at one of four doses: 2e7 gc, 2e8 gc, 4e8 gc, and 2e9 gc. They were sacrificed 4 weeks post injection and the number of ONL layers was quantified. Quantification of whole eyecup sections revealed a reduction in the number of ONL layers at doses of 2e8 gc Best1::GFP or higher, relative to uninjected samples (Figure S2A-B). The 2e7 gc Best1::GFP dose did not give noticeable toxicity to the ONL when compared to uninjected eyes at the same timepoint. Therefore, we designated this dose as a “tracer” dose of Best1::GFP to track injection sites when using AAV vectors that did not encode a fluorescent protein. Given that 2e8 gc, 4e8 gc, and 2e9 gc gave comparable toxicity profiles over time, we picked 4e8 gc as a dose to measure toxicity and host responses. This AAV dose has been previously used by our lab to achieve full RPE coverage and therapeutic rescue of RPE and photoreceptors in mouse models of oxidative stress.^43,52,54^

### Analysis of differential gene expression in RPE infected with Best1::GFP vs. Rho:GFP

To screen for potential mechanisms of toxicity generated by Best1::GFP compared to Rho::EGFP injections, bulk RNA-sequencing was performed. RPE RNA from mice transduced at P0 with 4e8 gc of Best1::GFP or Rho::GFP in contralateral eyes, or uninjected eye controls, were used for this analysis. Eyes were harvested at 1 or 2 weeks post injection. These timepoints were chosen to identify pathways that might initiate the toxicity. Principal Components Analysis (PCA) performed on normalized counts indicated that the greatest contribution to variability within the dataset was associated with the timepoint (PC1, 36%, Figure S3A). The type of AAV was the second greatest factor, as seen from PC2 and PC3 (14% and 9%, respectively, Figure S3B). Principal components 1-3 accounted for ∼60% of the total variance in the dataset, as shown by a cumulative sum of the variance contributed by each PC (Figure S3C). Differential gene expression analysis by DESeq2^49^ identified 2261 genes with a log2 fold change > 1 and an adjusted p-value < 0.05 for differential expression among all four groups. A heatmap of the relative level of gene expression by group illustrated the importance of timepoint compared to AAV (Figure S4A).

Gene ontology (GO) enrichment analysis indicated that the most significant difference between the Best1::GFP and Rho::GFP samples at week 1 was upregulation of ribosome biogenesis and other ribosome-associated processes (Figure 2A and Table S1). Even more notable was the lack of immune upregulation at Week 1 compared to Week 2. Among the top 25 enriched GO biological process (BP) terms associated with genes upregulated in the Best1::GFP samples at Week 1, the only suggestion of immune activation was "response to dsRNA" within the "response signaling to pathway" category (Figure 2A). Cell stress was also apparent, as “intrinsic apoptotic signaling pathway in response to ER stress” was another GO term enriched at week 1. In contrast, at week 2, there were multiple immune-related categories of enriched GO terms. These included “cytokine-mediated signaling pathway”, “regulation of tumor necrosis factor (TNF) production”, “regulation of innate immune response”, “response to interferon-beta”, and “myeloid leukocyte activation” (Figure 2B and Table S2). As with the week 1 GO analysis, week 2 GO analysis continued to show upregulation of ribosomal and translation-related pathways (Figure 2B). For GO terms associated with downregulated genes, the downregulation of retinoid and retinol metabolism was observed at week 1 (Figure S5), and persisted at week 2 (Figure S6A). As retinol metabolism is critical for maintenance of the visual cycle, which the RPE plays a critical role in, this pathway downregulation may further sensitize the RPE and retina to toxicity.

**Figure 2:**
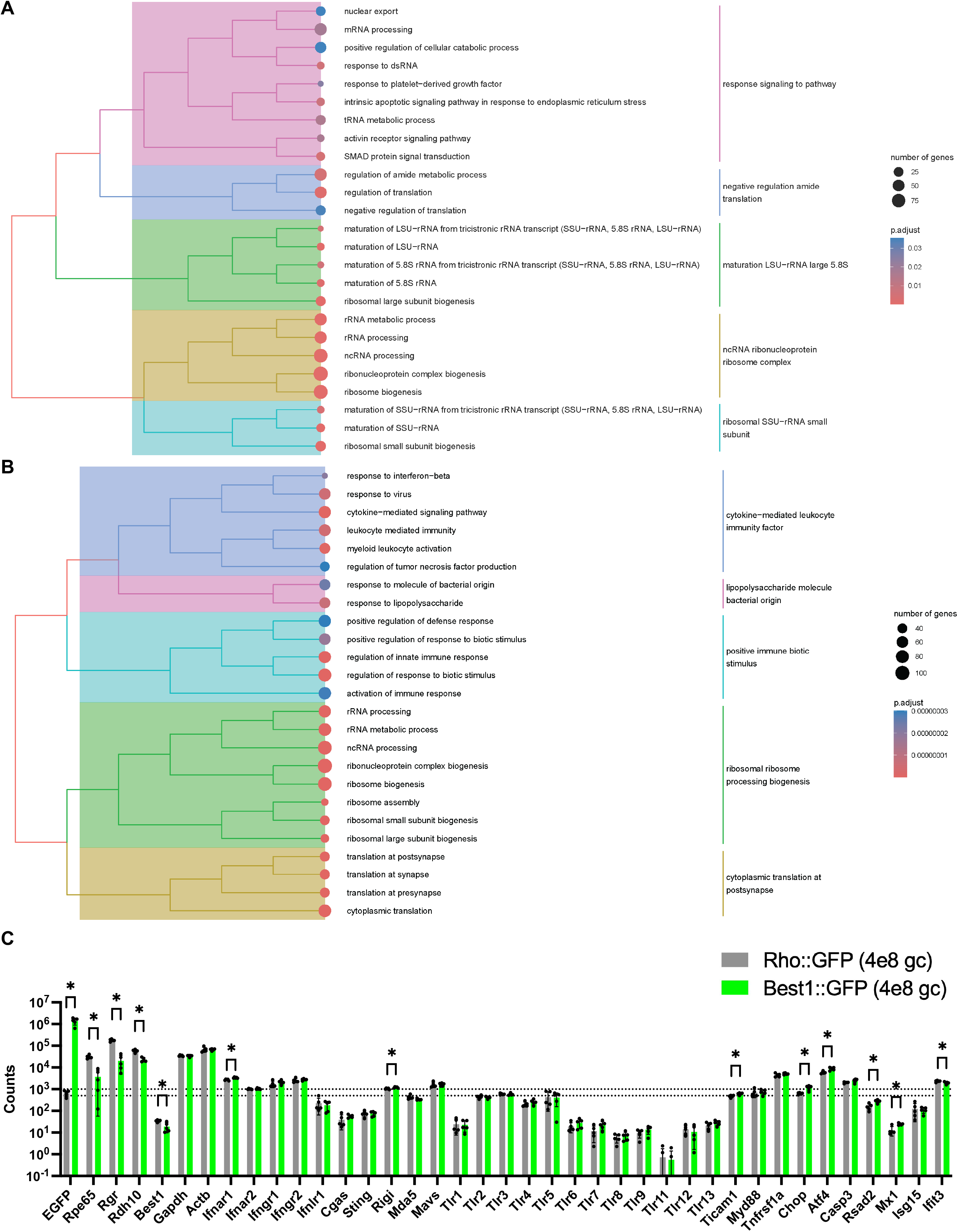
Bulk RNA sequencing of RPE cells to reveal candidate mechanisms of ocular toxicity from Best1::GFP expression. P0 pups were subretinally injected with 4e8 gc of Rho::GFP or Best1::GFP in contralateral eyes. Enriched RPE cells were collected at 1 week or 2 weeks post-transduction for bulk RNA-sequencing (n=4-5 eyes per group). (A) Tree plots of the top 25 Gene Ontology Biological Process terms associated with upregulated DE genes at week 1. Pathways are upregulated in the Best1::GFP relative to the Rho::GFP infection context. (B) Tree plots of the top 25 Gene Ontology Biological Process terms associated with upregulated DE genes at week 2. Pathways are upregulated in the Best1::GFP relative to the Rho::GFP infection context. (C) Read counts (normalized using DESeq2’s median of ratios method^50^) for selected stress and immune-related genes at 1 week post injection of 4e8 gc Best1::GFP or 4e8 gc Rho::GFP. Read counts for the EGFP transgene, the Best1 endogenous gene, some housekeeping genes (GAPDH, ACTB), and some visual cycle genes (RPE65, RGR, RDH10) are plotted for reference. The dotted lines represent read counts of 500 and 1000 and are included to help visualize expression levels on the plot (n=5 eyes per group, p<0.0001 for Rgr, Rpe65, and Rdh10, *p<0.001 for EGFP, *p<0.01 for Ifnar1, Chop, Mx1, Atf4, Rsad2, Ticam1, Rigi, and Best1, *p<0.05 for Ifit3, mean ± SD, multiple unpaired t-tests).

To focus on the genes underlying some of the immune pathway signatures identified by the RNA analysis, counts for individual immune or cell stress response-related genes were plotted at 1 or 2 weeks of age (Figure 2C and Figure S6). EGFP and additional genes normally highly expressed in the RPE (e.g., RPE65, RGR, RDH10) were plotted for reference. EGFP read counts were orders of magnitude higher in Best1::GFP vs. Rho::GFP injected RPEs at 1 week of age (Figure 2C). As noted earlier, retinol metabolism genes (e.g., RPE65, RGR, RDH10) were downregulated in Best1::GFP vs. Rho::GFP RPEs at week 1 (Figure 2C). Many immune-related receptors/sensors did not significantly differ in their expression level regardless of whether Best1::GFP or Rho::GFP was injected. However, stress-response related genes (e.g., CHOP, ATF4) and some ISGs) genes (e.g., RSAD2 and MX1) did increase following Best1::GFP relative to Rho::GFP injection at week 1 (Figure 2C). Compared to the visual cycle gene counts, expressed in the 10^4-10^5 read count range, most immune and/or stress-related genes were 10-1000x fold less expressed, with expression levels in the 10^2-10^3 range at week 1 (Figure 2C). When assessing counts per million reads (CPM), most immune- or stress-related genes surveyed were expressed at a CPM<100 following Best1::GFP injection at week 2 (Figure S6A). For comparison, unstimulated microglial CPM data for these same genes, obtained from prior literature,^55^ showed CPM ∼10x higher for many genes (Figure S6B). This suggests that stress and immune-related genes are moderately expressed in mice as young as 1-2 weeks of age, which may enable them to respond to viral infections.

### Assessment of the contributions of intrinsic, innate, and adaptive immune pathways on AAV-associated toxicity

RNA-seq data suggested that early cell stress in the RPE, and/or cytokine-mediated cellular toxicity, might cause toxicity by Best1::GFP. To investigate these potential mechanism(s) further, knockout (KO) mice were used. Toxicity was first evaluated in ten KO strains chosen on the basis of previous linkage to AAV toxicity in preclinical or clinical studies,^12,26,29,30,56–62^ as well as the RNA-sequencing data in Figure 2. Two interferon pathway knockout strains (IFNAR1 KO and IFNGR1 KO), one inflammatory cytokine receptor KO strain (TNFR1 KO), a complement pathway KO strain (C3 KO), a toll-like receptor signaling KO strain (MyD88/Ticam1 dKO, also referred to as TLR dKO), a cytosolic DNA sensing KO strain (cGAS KO), and a double-stranded RNA sensing KO strain (MAVS KO) comprised the innate immune KO mice tested. Strains deficient for B cells and T cells (RAG1 KO) or B cells, T cells, and natural killer cells (IL2RG KO) comprised the adaptive immune KO mice tested. Lastly, an unfolded protein response pathway KO strain (CHOP KO) was the only KO strain tested that represents an intrinsic response pathway KO, rather than an innate or adaptive immune response pathway KO. P0-P1 mice were uninjected, or were subretinally injected with Rho::EGFP or Best1::EGFP, and eyes were harvested at 4-5 weeks of age for RPE and/or retinal toxicity analyses (Figure 3A).

**Figure 3:**
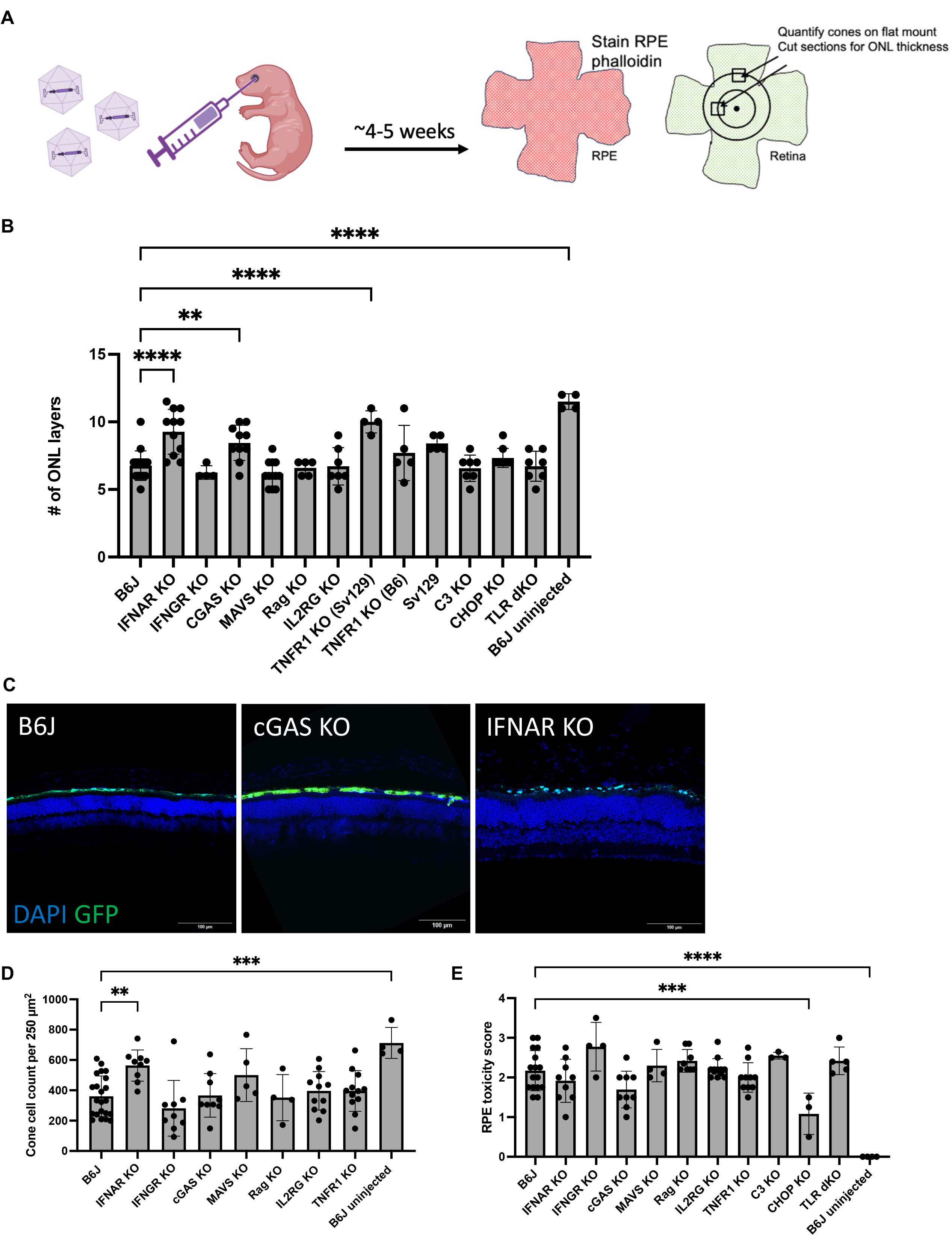
Knockout (KO) mouse injection screen to determine the contributions of intrinsic, innate, and adaptive immune pathways on AAV-associated toxicity. For panel B, two strain backgrounds of the TNFR1 KO mouse (Sv129 or C57BL/6, e.g., B6) were used, as indicated by the labels. For panels D and E, only the C57BL/6 TNFR1 KO strain was used. The TLR dKO mouse strain refers to Myd88/Ticam1 double KO mice. A) Schematic of the immune KO mouse screening experiment to alleviate toxicity. Neonatal mouse pups of the indicated genotype were injected with 4e8 gc Best1::GFP and eyes were harvested for RPE flatmounts, retinal flatmounts, or whole eyecup sections 4-5 weeks post injection. Figure created with BioRender. B) Median number of ONL layers in midperipheral regions of whole eyecup sections from Best1::GFP injected eyes of the indicated genotype (n=4-17 eyes per group, ****p<0.0001, **p=0.005, mean ± SD, one-way ANOVA with Dunnett’s multiple comparison’s correction). Uninjected control samples are the same as shown in Figure 1. C) Representative whole eyecup sections for C57BL/6J mice or selected immune KO strains from the plot shown in panel B. Blue: DAPI. Green: Best1::GFP expression. Scale bar: 100 µm. D) Cone quantification from retinal flatmounts (n≥4 eyes per group, **p=0.003, ***p=0.0001, mean ± SD, one-way ANOVA with Dunnett’s multiple comparison’s correction). All mice were 4-5 weeks of age at harvest. E) Semi-quantitative scoring of RPE cell toxicity (n≥3 eyes per group, mean ± SD, ****p<0.0001, ***p=0.001, one-way ANOVA with Dunnett’s multiple comparison’s correction). Uninjected control samples are the same as shown in Figure 1.

The mean number of layers in the ONL in Best1::GFP-injected eyes of most KO strains, which reflects the number of rod photoreceptors as they are 95% of cells in this layer,^63^ was comparable to the average number of layers seen in injected B6J eyes (Figure 3B-C). However, IFNAR KO mice showed higher ONL layer counts from this screen, with a mean of ∼9.3 layers, relative to ∼6.8 layers for B6J mice (Figure 3B-C). Another strain which showed protective effects, cGAS KO, had a higher number of ONL layers, e.g., ∼8.5 vs ∼6.8 for B6J injected mice (Figure 3B-C). Interestingly, TNFR1 KO mice showed a higher number of retinal layers, suggesting an alleviation of toxicity. However, as this KO allele was on a Sv129 background, and the other KO strains were on a B6J background, there was a possibility that the phenotype was due to the Sv129 background. Indeed, when TNFR1 KO was tested on a C57BL/6 background, there was no alleviation of toxicity (Figure 3B-C). An injected Sv129 background control strain likewise revealed a slightly higher number of layers compared to equivalently injected B6J mice (Figure 3B-C). This result demonstrates that the genetic background of the mouse strains can affect the response to AAV-induced toxicity.

To further examine the retinas from the KO mouse strains, cone counts from Best1::GFP-transduced retinal flatmounts from B6J or KO mice were compared. IFNAR KO, but not cGAS KO nor any other KO strain, had significantly more cones relative to wild type controls (Figure 3D). These results suggest that cones might be slightly more sensitive to toxicity than rods, and/or use different pathways that were not tested here, to trigger toxicity. However, both photoreceptor types roughly followed the same rescue trends across all immune KO strains tested.

To assess toxicity in the RPE, RPE flatmounts were collected and a semi-quantitative analysis of RPE toxicity was performed. Surprisingly, RPE cells were not noticeably healthier in IFNAR KO mice, in contrast to the rods and cones (Figure 3E and Figure S7). The only strain that gave noticeably healthier RPE cells following Best1::GFP infection was the CHOP KO strain (Figure 3E). Photoreceptor toxicity was not alleviated in the CHOP KO.

Given the protective effects of the IFNAR KO strain on both rods and cones following Best1::GFP injection, it was of interest to determine if the protective effects would extend to a virus with a higher expression level of GFP. An AAV expressing GFP under the stronger, ubiquitous CMV promoter, CMV::GFP, was injected subretinally at the same dose (4e8 gc) into B6J or IFNAR KO pups. Four weeks post injection, RPE and retinal flatmounts were harvested from these animals. Cone counts were again preserved following CMV::GFP injection into IFNAR KO animals relative to B6J controls, but RPE cells were not preserved (Figure S8).

To check whether the KO strains have comparable RPE morphologies, cone counts, and ONL layer counts at baseline, eyes from uninjected mice, or mice injected with 4e8 gc Rho::GFP, were collected at 1-2 months of age and assessed by each cell-type specific readout. RPE cells were comparably healthy among all strains tested. Cone counts were comparable among all strains tested except for a slight decrease in cone counts in IFNGR KO mice relative to B6J controls (Figure S9A). The number of ONL layers was comparable among most strains, except for the TNFR1 KO (Sv129 background). Rho::GFP injected samples from this strain had slightly higher ONL layer counts at baseline, matching earlier observations about Sv129 background-related rescue effects (Figure S9B). These results strengthen the interpretation that the KO strains (IFNAR KO and cGAS KO) that provided ONL layer retention did so due to the KO allele, rather than due to the specific strain background.

### Assessment of type I interferon responses in RPE and retina following AAV injection

To further investigate the protective effects of the IFNAR KO strain, differentially expressed genes were analyzed. Genes whose expression was enriched in Best1::GFP vs Rho::GFP infected RPE samples at weeks 1 and 2 post infection (p-adj value<0.05) were quantified. The expression levels of 1688 genes were enriched at week 1, and 1461 genes were enriched at week 2 (Tables S3-S4). Each list of gene names was submitted to the online Interferome database.^64^ This database comprises genes involved in the interferon response, including the ISGs. The number of ISGs found among the week 1 and week 2 differentially expressed genes was 245 and 281, respectively (Tables S5-S6). Of these, the number of differentially expressed genes induced by a factor of 2 or more in the Best1::GFP vs Rho::GFP context was 47 (week 1) and 59 (week 2) genes, respectively (Tables S5-S6). Selected differentially expressed ISGs from the week 2 dataset are plotted in Figure 4A. Table S7 contains a normalized counts matrix of all genes. These observations support the existence of an active type I interferon response in RPE cells.

**Figure 4:**
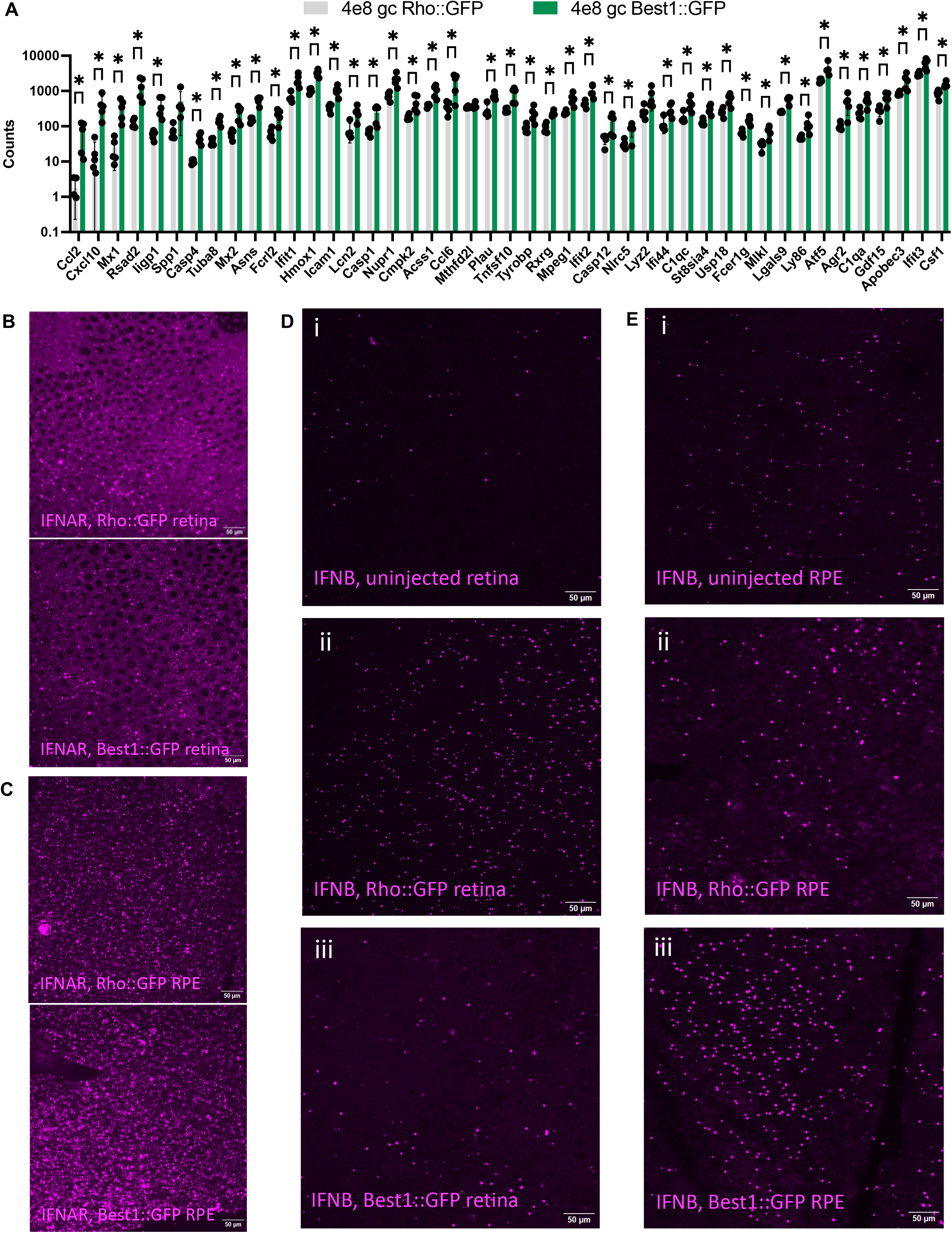
Analysis of expression of interferon beta (IFNB), the type I interferon receptor (IFNAR), and interferon stimulated genes (ISGs) following AAV infection. A) Selected differentially expressed ISGs (45 total). Week 2 read counts (normalized using DESeq2’s median of ratios method^50^) from Rho::GFP (gray) and Best1::GFP (green) injected RPE RNA-sequencing data are plotted (n=5 eyes per group, *p<0.001 for Asns, Hmox1, Rxrg, Tuba8, Lgals9, and Casp4, *p<0.01 for Plau, Fcer1g, Csf1, Icam1, St8sia4, Fcrl2, Mlkl, Ifit1, Nupr1, Acss1, Usp18, Mx2, Mpeg1, Gdf15, Ifit2, Mx1, Lcn2, and Rsad2, *p<0.05 for Nlrc5, Tnfsf10, Ifit3, C1qa, Iigp1, Ifi44, Casp1, Agr2, Ccl6, Atf5, Cxcl10, Cmpk2, C1qc, Ccl2, Tyrobp, Apobec3, Casp12, and Ly86, mean ± SD, multiple unpaired t-tests). B) HCR-FISH for IFNAR1 transcripts was performed on retinal flatmounts from B6J mice injected at birth with 4e8 gc Rho::GFP (top) or 4e8 gc Best1::GFP (bottom) and harvested at P14. Scale bar: 50 µm. C) HCR-FISH for IFNAR1 transcripts was performed on RPE flatmounts from B6J mice injected at birth with 4e8 gc Rho::GFP (top) or 4e8 gc Best1::GFP (bottom) and harvested at P14. Scale bar: 50 µm. D) HCR-FISH for IFNB1 transcripts was performed on retinal flatmounts from B6J mice that were uninjected at birth (i) or injected with 4e8 gc Rho::GFP (ii) or 4e8 gc Best1::GFP (iii) and harvested at P14. Scale bar: 50 µm. E) HCR-FISH for IFNB1 transcripts was performed on RPE flatmounts from B6J mice that were uninjected at birth (i) or injected with 4e8 gc Rho::GFP (ii) or 4e8 gc Best1::GFP (iii) and harvested at P14. Scale bar: 50 µm.

To test whether retinal cells sense and respond to interferons post AAV injection, fluorescent in situ hybridization (FISH) was performed using the hybridization chain reaction (HCR) method. Probes were made for interferon-associated transcripts and applied to RPE and retinal flatmounts from animals injected with Best1::GFP or Rho::GFP (Figure 4B-E). IFNAR1 FISH signal was observed in both RPE and retinal tissues injected with either virus (Figure 4B-C). In addition, FISH signals for IFNB1 were observed in uninjected RPE and retinal tissue (Figure 4D-E). However, the signal was greater following injection of Rho::GFP and Best1::GFP in both the RPE and retina. Interestingly, it appeared that the retina expressed more IFNB1 in response to Rho::GFP than to Best1::GFP injections. Conversely, there was higher IFNB1 signal in Best1::GFP infected RPE than in Rho::GFP infected RPE (Figure 4D-E). This likely is due to GFP expression leading to IFNB1 expression in the respective tissue types. These FISH results support the existence of a type I interferon response in Rho::GFP and Best1::GFP infected RPE and retina. Given the ISG analysis in Figure 4A, the observed toxicity in the context of Best1::GFP infection may result from elevated ISG induction, elevated GFP-induced damage to the RPE, or a combination of both.

### Assessment of the contributions of microglia, pyroptosis, and chemokine receptors on AAV-associated toxicity

The Rag1 KO and IL2RG KO mouse strains eliminate almost all adaptive immune responses, i.e., B cells, T cells, and NK cells. As there was not a reduction in AAV-induced toxicity following neonatal AAV injections into these strains, it was unlikely that adaptive immune responses contribute to the toxicity. The screen did not, however, exclude a contribution by the myeloid cells: retinal microglia or peripheral infiltrating macrophages. Microglia are the main tissue-resident immune cells in the retina and can be a source of retinal inflammation through their ability to secrete pro-inflammatory cytokines.^65^ Prior work from our lab showed that microglia were recruited to the ONL in response to toxic AAV subretinal injections, and two pro-inflammatory cytokines, TNFa and IL-1B, were significantly induced.^12^ To determine whether microglia contribute to retinal toxicity, they were pharmacologically depleted as early in development as possible.

Microglial depletion was attempted using a CSF1R inhibitor (PLX5622) incorporated into rodent chow. PLX5622 or regular chow was administered to the lactating dam and her pups at P7 and continuously provided until the pups reached P28, when the pups were sacrificed for histology (Figure S10A). The B6J pups in this experiment were injected with 4e8 gc Best1::GFP at birth, and tissue was harvested 4 weeks later. A statistically significant rescue of RPE cells was seen (Figure S10B). However, the number of cones quantified on retinal flatmounts did not show an appreciable rescue in animals fed the PLX5622 diet vs. the control diet (Figure S10C). These retinal flatmounts were subsequently sectioned and stained with DAPI to count the number of ONL layers, but no rescue was seen in animals treated with PLX5622 vs. controls by this metric (Figure S10D). An analogous experiment performed using an injection dose of 4e9 gc Best1::GFP also demonstrated partial rescue of RPE cells (Figure S10E). In this case, cone counts were slightly higher, but the number of ONL layers was not different between PLX5622 treated and control mice (Figures S10F-G). To determine the efficiency of microglial depletion in these two experiments, immunohistochemistry for a marker of microglia and macrophages, IbaI, was used on RPE flatmounts (Figure S11A-B and Figure S12A-B). Manual counting of IbaI staining in these images revealed a 91% percent depletion for the eyes injected with 4e8 gc Best1::GFP and 98% for those injected with 4e9 gc Best1::GFP (Figures S11C and Figure S12C). These data indicate that the PLX5622 feeding regimens used in this study depleted microglia in PLX5622 chow-fed animals by P28.

As pups rely on their mother’s milk for nourishment through postnatal day 21 (∼P21) before starting to eat solid food directly, it was possible that not enough of the PLX5622 drug passed through the mother’s milk to deplete microglia in the pups prior to this age. To determine the preweaning age depletion efficiency, uninjected Cx3cr1-GFP heterozygous reporter pups, which have GFP+ myeloid cells, and their nursing dams were administered PLX5622 or regular chow from P8-P19. At P19, the mice were harvested for quantification of Cx3cr1+ cells (Figure S13A-B and S13E). As positive controls, postweaning P31 Cx3cr1-GFP heterozygous reporter mice were fed PLX5622 or regular chow for 1 week, then harvested to test for depletion of Cx3cr1+ cells (Figure S13C-E). Quantification of the adult depletion experiment revealed ∼100% Cx3cr1+ cell depletion after 1 week of CSF1R inhibition (Figure S13E). Quantification of the pup depletion experiment revealed a slight but not statistically significant depletion at P19 (Figure S13E). These results suggest that most of the depletion occurs when the pups start to eat solid food around the third week of life. The lack of substantial rescue of cones and ONL layers in these experiments may thus have been limited by the actual depletion window of the chow, ∼1 week before the experiment ended.

Given the inability to fully deplete microglia during the entire developmental period, an alternative approach was taken to investigate whether there were pathways that microglia might use to induce ocular toxicity. Retinal flatmounts injected with 4e8 gc or 4e9 gc Best1::GFP were harvested 4-5 weeks post infection and immunofluorescence was used to detect the inflammasome marker ASC. This marker is induced when cells undergo pyroptosis. ASC specks were present in infected retinal flatmounts in much higher numbers than in uninfected retinal flatmounts. The specks appeared to be localized to myeloid cell types, rather than to retinal cell types, based on the morphology of the ASC+ cells (Figure S14A). In order to use the Cx3cr1-GFP reporter mouse to determine if the ASC+ cells in the infected retina were myeloid cells, an AAV which does not encode a fluorescent protein was used. The CMV::null virus, shown to be toxic from prior work in our lab, was thus injected into the reporter strain (Figure S14B). Cx3cr1+ cells in these infected retinal flatmounts colocalized with the ASC marker, confirming the myeloid lineage of the ASC+ cells (Figure S14B). The Best1::6xSTOP-mutGFP AAV, previously constructed and characterized by our lab, was used here as a benign control. This AAV carries a mutated (non-expressing) GFP transgene and lacks any open reading frames >73 amino acids in its genome sequence.^43^ The Best1::6xSTOP-mutGFP infected retinal flatmounts showed many fewer ASC+ cells (Figure S14A). Together, these data suggest that retinal myeloid cells may undergo pyroptosis predominantly in response to toxic AAV injections.

As ASC is a marker of cells dying via pyroptosis, we investigated if the pyroptosis pathway was active in tissues exposed to toxic AAV. This possibility was investigated by using gasdermin D (GSDMD) KO mice, as GSDMD is required for pyroptosis. GSDMD KO mice were injected with 4e8 gc Best1::GFP and sacrificed at 4 weeks post infection. The RPE morphology, number of cones, and number of ONL layers were analyzed. No difference was seen in the cone or ONL readouts relative to those of B6J injected controls, although the RPE was seen to be more negatively impacted by the viral injection in the GSDMD KO than in the B6J injected animals (Figure S14C). The GSDMD KO strain has a B6J/N background, which harbors an rd8 allele, which causes retinal degeneration.^66^ To determine if this strain of mice has a phenotype without AAV injection, which might affect the interpretation of the AAV injections, uninjected GSDMD KO mice were directly compared to B6J uninjected controls. The RPE, cone, and ONL readouts were not significantly different between the B6J and GSDMD KO strains (Figure S14D). These observations indicate that, at least at the 4 week timepoint, these strain backgrounds are equivalent.

To examine whether loss of chemokine receptors could reduce AAV-induced toxicity, Cx3cr1-GFP/Ccr2-RFP mice were tested. Cx3cr1-GFP is expressed in monocytes, microglia, dendritic cells, and NK cells, whereas Ccr2-RFP is expressed in peripheral monocytes, T cells, and NK cells. The GFP and RFP alleles create loss of function of the receptors.^67,68^ Homozygous Cx3cr1-GFP/Ccr2-RFP mice were injected with 4e8 gc Best1::GFP at birth and sacrificed 4 weeks later. RPE morphology, the number of cones, and the number of ONL layers were equivalent between the Cx3cr1/Ccr2 double KO mice and the double heterozygous or WT counterparts (Figure S15).

### Assessment of Transgene Dependence on AAV-Associated Ocular Toxicity

An important question relevant to AAV gene therapy is whether ocular toxicity is dependent upon the specific transgene. One factor that might be relevant, especially for injections in adults where adaptive immunity is active, is whether toxicity depends on whether the transgene is a “self” gene or a foreign gene. To investigate this possibility, and the generality of AAV-induced ocular toxicity in mice, a panel of AAVs encoding different transgenes was tested. The transgenes included the toxic and nontoxic AAVs used so far in this study, as well as AAVs encoding various self and foreign transgenes. These AAVs were individually injected into B6J mouse eyes at a fixed dose. The AAV-encoded self-transgenes included: (1) beta-2-microglobulin (Best1::B2M), a subunit of major histocompatibility complexes that aids in antigen presentation, (2) cathepsin D (Best1::Ctsd), a protease that digests shed outer segments of the photoreceptors, and (3) RPE65 (Best1::RPE65), a protein that produces 11-cis retinol for the visual cycle. In addition to being self genes expressed by the RPE, these transgenes were selected as their endogenous expression levels span ∼3 orders of magnitude (Figure 5 legend). One foreign transgene, CasMini (Best1::CasMini), an engineered compact Cas protein derived from Cas12f, which is used for genome editing applications,^69^ was also included. The level of toxicity elicited by each AAV was assayed 4-5 weeks post injection using the number of ONL layers as a readout.

**Figure 5:**
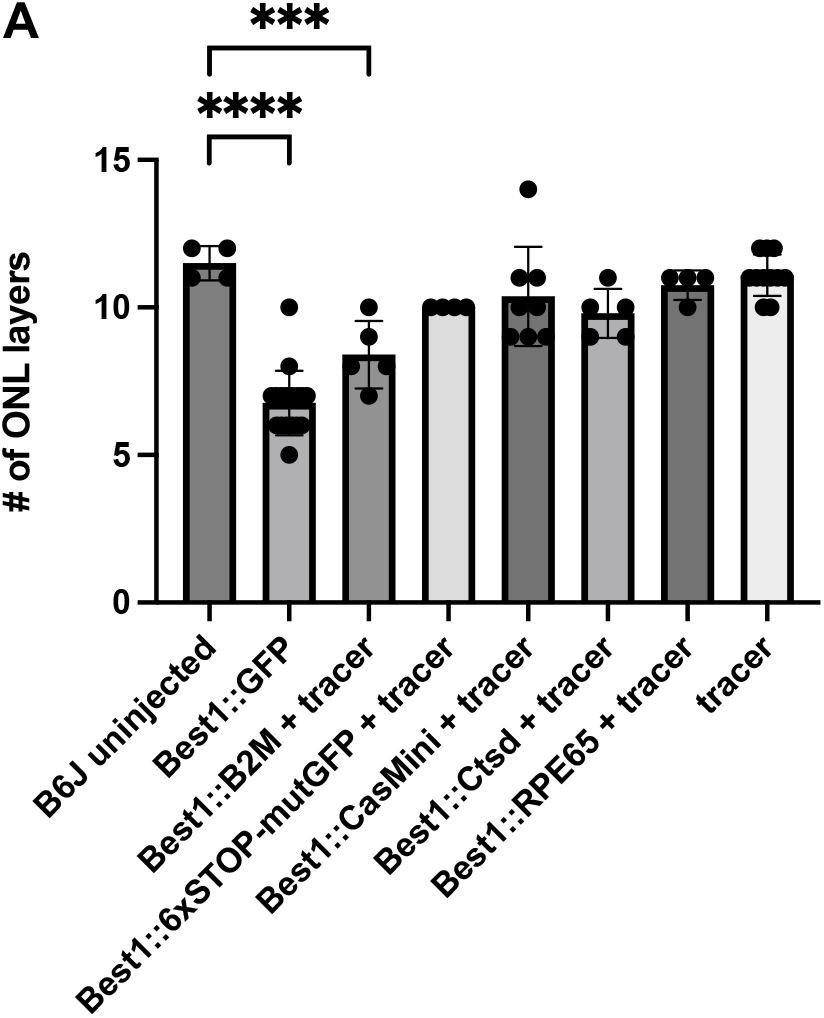
Effect of transgenes on AAV-associated ocular toxicity. All mice in this figure were harvested at 4-5 weeks post injection. B6J controls (either injected with 4e8 gc Best1::GFP or uninjected) were previously plotted in Figure 3. Data for the “tracer” dose of Best1::GFP (2e7 gc) is replotted here from Figure S2. For reference, the read counts (normalized using DESeq2’s median of ratios method^50^) for the EGFP transgene is ∼10^6, whereas the counts for the endogenous transcripts CTSD, B2M, and RPE65 are ∼10^5, ∼10^4, and ∼10^3, respectively. A) Quantification of the number of ONL layers remaining at 4 weeks for a panel of Best1::transgene expressing AAVs, subretinally injected at birth and harvested 4-5 weeks post injection. Each Best1::transgene expressing AAV was administered at 4e8 gc. The Best1::GFP tracer was co-injected at 2e7 gc to track nonfluorescent transgenes. Uninjected, age-matched C57BL/6J eyes were plotted for comparison (n=4-17 eyes per group, mean ± SD, ****p<0.0001, ***p=0.0003, one-way ANOVA with Bonferroni’s multiple comparisons correction).

At this timepoint, Best1::GFP toxicity elicited a ∼41% reduction in the number of ONL layers (∼6.8 layers) relative to age-matched uninjected B6J controls (∼11.5 layers, Figure 5A). Of the self transgenes, Best1::B2M significantly reduced the number of ONL layers by ∼27% compared to uninjected controls (∼8.4 vs. ∼11.5 layers, respectively). All other transgenes were not noticeably toxic compared to uninjected controls (Figure 5A). Nontoxic transgenes included two other “self” genes, Ctsd and RPE65, one “foreign” gene, CasMini, and a benign control AAV (Best1::6xSTOP-mutGFP). As noted earlier (Figure S2, replotted here for comparison), a tracer dose (2e7 gc) of Best1::GFP wasn’t toxic relative to uninjected control eyes, so this tracer dose was used to track injection success for all non-fluorescent AAV transgenes in this experiment.

### Assessment of AAV-Induced Toxicity in Mice Subretinally Injected as Adults

Neonatal subretinal injections were used to achieve full transduction of RPE and retina for toxicity evaluation. Full transduction is difficult to achieve from subretinal injections of adults due to the barrier imposed by mature RPE apical processes and photoreceptor outer segments. This barrier stops the flow of inoculum throughout the subretinal space. However, it was of interest to test whether adult mice are also sensitive to toxic effects from AAV injections, as human gene therapy is usually conducted in adults. We thus injected Best1::GFP at a dose of 2e10 gc in ∼1-2 uL per adult mouse eye (roughly equivalent to a ∼4e9 gc Best1::GFP dose in pups, delivered in ∼0.2 uL) for evaluation of toxicity at 4 weeks post injection. Adult-injected RPE flatmounts displayed RPE toxicity in response to 2e10 gc Best1::GFP at 4 weeks post injection (Figure S16A-B). Interestingly, RPE toxicity was lower than that observed from 4e9 gc Best1::GFP injected into neonates, while it was comparable to that of neonatal 4e8 gc Best1::GFP injections (Figure S16B). Cone counts were also evaluated from the same eyes used to collect RPE flatmounts and were not significantly reduced relative to uninjected eyes, although the exact area of retinal transduction was not readily apparent in each retinal sample for quantification. It’s possible that cone counting from untransduced areas of adult-injected eyes made this readout show a less significant loss of cones compared to neonatal retinas, which were only quantified if the corresponding RPEs were fully transduced (Figure S16C).

To overcome the issue of identifying transduced areas of retinas injected with Best1::GFP and also expand our toxicity characterizations further, we injected CMV::GFP, which expresses GFP in both RPE and retinal cell types, at a dose of 2e9 gc (in ∼1-2 uL/eye) in adult mice. This dose is roughly equivalent to the 4e8 gc (in ∼0.2 uL/eye) dose used in neonates. This AAV induced RPE loss following both neonatal and adult injections, but neonatal injected RPEs showed more toxicity than adult injected RPEs (Figure S17A). CMV::GFP also led to loss of cones, to roughly equal extents from injection of neonates and adults (Figure S17B). IFNAR KO adults injected with this vector did not look healthier compared to B6J injected adults by any metric. This suggests that host responses to AAV injections may be more complicated in adults than in developing mice, and/or they may use different mechanisms to respond to viral infections.

## Discussion

In this study, we sought to understand the mechanism(s) of AAV toxicity arising in what is generally accepted to be an immune-privileged organ, the eye. We previously reported that an AAV expressing GFP in the RPE via an RPE-specific promoter (Best1) or ubiquitous promoters (e.g., CMV, CAG, UbiC), caused RPE cells to become dysmorphic and die. These AAVs also induced loss of visual function, retinal structure, and led to death of photoreceptors.^12^ By contrast, photoreceptor specific promoters (e.g., CAR, RK, RedO, Rho) did not induce toxicity, even at high doses, using the same capsid, injection site, and preparation method. We chose GFP as an easily traceable transgene that also yielded the most toxicity at ∼1 month post infection compared to other transgenes tested (Figure 5A). This toxicity level allowed us to maximize the dynamic range of several assays and reliably distinguish rescue effects from genetic or pharmacological interventions.

Bulk RNA-sequencing of RPE cells infected with toxic (Best1::GFP) or benign (Rho::GFP) AAVs produced pathway hits potentially mediating AAV toxicity. Gene ontology analysis performed on genes differentially upregulated in Best1::GFP vs. Rho::GFP injected samples didn’t pick up immune-response related genes at week 1, apart from one category: “response to dsRNA.” Most genes upregulated at week 1 were broadly related to translation, rRNA, or mRNA processing. One such category, “intrinsic apoptotic signaling in response to ER stress,” included genes like CHOP and ATF4. These genes were induced following Best1::GFP relative to Rho::GFP injection and were still upregulated by week 2. Activation of these genes suggests that AAV-mediated overexpression of GFP in the RPE may initiate toxicity through a GFP-induced ER stress pathway. As discussed further below, the fact that the CHOP KO partially alleviated toxicity in the RPE suggests that ER stress may be part of the pathology from GFP expression. Potentially relevant to this observation is the level of GFP expression in the RPE relative to that of the photoreceptors. We previously performed FISH for AAV genomes in both the RPE and ONL from subretinal injections at a dose of 2.5e8 gc.^70^ We found approximately 10x more genomes in the RPE than in the photoreceptors. The level of GFP induced by Rho::GFP in photoreceptors vs Best1::GFP in the RPE would thus be expected to be considerably lower. However, injection of nearly 10x more Rho::GFP (2e9 gc) still did not induce toxicity in photoreceptors. These observations suggest that the RPE is more sensitive to GFP, and perhaps other overexpressed proteins, suffering from an ER stress response.

By week 2, translation-related pathways were still prominent in gene ontology analysis of differentially expressed genes. Additionally, however, upregulation of genes involved in cytokine release, antiviral responses, and leukocyte activation or infiltration was observed. This implied that a cell stress trigger initiates toxicity by week 1, which may drive cytokine/chemokine release resulting in immune cell recruitment to the RPE and ONL layers. Infiltrating immune cells may further contribute to cytokine release, antiviral signaling, and potentially exacerbate the RPE and retinal toxicity observed by ∼1 month.

At weeks 1 and 2, retinol metabolism genes, e.g., RPE65, RGR, and RDH10, were downregulated in RPE infected by Best1::GFP vs Rho::GFP AAVs (Figure 2C and Figure S6A). As noted earlier, downregulation of multiple genes in the visual cycle may sensitize the RPE and retina to toxicity and contribute to vision loss.^12^ Interestingly, a tamoxifen-inducible RPE65 KO mouse line retained healthy RPE morphology and ONL thickness.^71^ An RGR KO mouse line and a doxycycline-inducible RDH10 KO mouse line also retained healthy ONL thickness, but RPE morphology was not assessed.^72,73^ No mouse line has been reported that has a reduction of expression of several visual cycle genes, so it is unclear whether RPE cells in such a mouse would present with toxicity similar to that induced by Best1::GFP. Future work may address this potential source of pathology by reducing or eliminating a combination of visual cycle genes.

To identify intrinsic, innate, or adaptive immune responses that could cause toxicity, we selected several mouse KO strains for pathways that had been previously linked to AAV-associated toxicity in the literature^12,26,29,30,56–62^ and our RNA-sequencing analyses. The pathways covered by the KO strains included innate immune pathways, such as type I and II interferon responses, cytosolic DNA sensing, dsRNA sensing, TLR-dependent sensing, complement responses, and TNF receptor mediated responses. KO strains for adaptive immune pathways (B, T, and NK cell-mediated responses) and cell intrinsic responses (ER stress) also were tested. To evaluate the effects of these KO strains on toxicity, three cell-type specific toxicity assays of RPE, rod, and cone health were used. The IFNAR KO strain reduced the toxicity observed for rods and cones, but not for RPE, while the cGAS KO strain reduced toxicity only for rods. The reduction in toxicity was in all cases partial, perhaps due to multiple pathways leading to toxicity. As photoreceptors are dependent on the RPE for survival, and since RPE toxicity persisted in these strains even when photoreceptor toxicity was alleviated at the 4 week timepoint, it may be the case that RPE toxicity will eventually cause photoreceptor degeneration. The negative results for the other KO strains may be equally informative. The fact that both the Rag1 KO and IL2RG KO mice didn’t rescue RPE, rods, or cones implies that the adaptive immune system may not be sufficiently developed in mice at this age, as prior work suggests.^74^ The type II interferon receptor KO (IFNGR KO) also didn’t reduce toxicity. This supports the Rag1 KO and IL2RG KO results, since primary producers of type II interferons are T cells and NK cells, which are absent in the Rag1 and IL2RG strains.^75^ Adaptive responses might, however, become relevant for adult subretinal injections. Prior literature has suggested the presence of cellular infiltrates comprised of CD4+ and CD8+ T cells, NK cells, and peripheral macrophages in adult eyes in response to subretinal injection of a CAG-GFP AAV.^76^ Causal relationships using loss-of-function approaches for adaptive immunity were not carried out, but are warranted.

The CHOP KO strain was used to study the contribution of stress pathways, which include ER stress, dsRNA responses, amino acid deprivation, and heme deficiency. CHOP is a pro-apoptotic transcription factor activated in response to these different forms of cellular stress.^40^ CHOP was differentially expressed between Best1::GFP and Rho::GFP infected RPE flatmounts in the RNA-seq data at weeks 1 and 2 post infection. From the RNA-seq data, gene ontology pathway analysis suggested that ER stress could be the initiating factor in the RPE that triggers downstream immune response and toxicity, killing photoreceptor cells. The differential responses of the RPE and photoreceptor cells to the CHOP and innate immune pathways are consistent with this model. The CHOP KO strain benefitted RPE cells, but photoreceptor toxicity was not reduced, while the IFNAR KO and cGAS KO strains benefitted photoreceptors but not the RPE. In addition to the suggestion of different mechanisms of pathology in the RPE and photoreceptors, the responses to the different KO strains might indicate that RPE cells can initiate signaling events that impact the health of photoreceptors, even when their own response to toxicity is blunted, i.e., in the CHOP KO.

The ONL layer rescue from the cGAS KO strain points to a role for cytosolic double-stranded DNA (dsDNA) sensing in response to AAV subretinal injections. AAV contains a predominantly single-stranded DNA genome, but some double-stranded regions are formed near where the ITRs form their characteristic T-shaped structures, and in the process of double strand synthesis of the viral genome. However, since both toxic (Best1::GFP) and benign (Rho::GFP) AAVs have the same genome structure and replication pathway, it was initially unclear how cGAS signaling could contribute to toxicity following Best1::GFP but not Rho::GFP injection. One possibility is that cGAS activation may trigger a type I interferon response in the eye, resulting in ISG induction. Induction of a subset of ISGs, or higher levels of ISGs beyond those induced by Rho::GFP, may occur due to the pathological response in the RPE to GFP. These may include Mx1, Mx2, and Rsad2 (Figure 4A). A heightened ISG response could contribute to the higher retinal toxicity generated by Best1::GFP vs. Rho::GFP.

To investigate a role for myeloid cells in ocular toxicity, several approaches were taken. Mice depleted for microglia, a GSDMD KO strain, and double chemokine receptor (CX3CR1/CCR2) KO strains were investigated. They did not alleviate toxicity. These data do not rule out the possibility that myeloid cells contribute to toxicity, perhaps through interferon signaling. It is likely that microglia respond to and generate their own interferon signaling, as prior literature has shown transcriptomic activation of interferon pathway components following intravitreal injection of AAVs in mice.^77^ Further investigation of this possibility is required to determine if they contribute to toxicity from subretinal injection of AAV.

The TNFR1 KO mouse was tested because TNFA, a proinflammatory cytokine, was highly induced by toxic AAV injection relative to nontoxic AAVs in our prior work.^12^ Unfortunately, we noticed a background-dependent rescue effect with the Sv129-derived TNFR1 KO strain that was eliminated when a C57BL/6-derived TNFR1 KO strain was tested. To completely rule out other pro-inflammatory cytokines or pyroptotic-related mechanisms from causing toxicity, the KO screen could be expanded to include mouse strains deficient in other pro-inflammatory or pyroptotic-related genes (e.g., IL-1B or IL-1R KO, IL-6 or IL-6R KO, caspase 1 KO, etc.) KO of multiple pathways in a single strain may be required to fully explore this possibility.

The MyD88/Ticam1 double KO mouse, which is broadly deficient for TLR signaling, did not rescue RPE or photoreceptors. This is in accord with the lack of high expression of TLR sensors in 1-2 week old animals per our RPE RNA-seq data (Figure 2C and Figure S6A). Specific TLRs have been previously shown to respond to components of AAV like the capsid (TLR2)^26^ or unmethylated CpG genome motifs (TLR9).^27–29^ Regarding TLR9, inhibitory sequences incorporated into AAV genomes were shown to antagonize TLR9 activation. This reduced innate, and subsequently, adaptive, immune responses across AAV doses, transgenes, routes of administration, and animal models.^78^ However, neonatal subretinal injections with the Tlr9 blocking sequence in mice did not show a reduction in toxicity, though it did following subretinal injection in juvenile pigs.^78^ Nonetheless, as the inhibitory sequences may not have completely blocked TLR9 signaling, we tested MyD88/Ticam1 double KO mice, which would eliminate all TLR activity. No reduction in toxicity was seen.

The C3 KO mouse was tested because the complement system has been shown in previous studies to mediate thrombotic microangiopathy (TMA), a condition where antibody-dependent blood coagulation in response to AAV injection can result in organ failure and death.^79^ This phenomenon has only been observed, however, for systemic AAV applications (e.g., liver-, muscle-, or brain-directed gene transfer). In these contexts, massive viral loads in the bloodstream can trigger antibody binding to AAV capsids, formation of a membrane attack complex (MAC) that destroys AAV particles, and generation of systemic inflammatory responses.^80^ Knocking out a key component of the complement pathway, C3, did not produce a benefit to RPE or photoreceptors, which may reflect the inaccessibility of complement components or antibodies to the subretinal space.

The MAVS KO mouse was tested based on prior literature that double-stranded RNA sensing pathways, e.g., RIG-I or MDA5-dependent, which both signal through the MAVS adaptor, can play a role in AAV-mediated immune responses in the brain and liver.^30,59^ Additionally, our RPE RNA-seq data suggested the possibility of a dsRNA response in the week 1 upregulated GO pathway hits. However, MAVS KO did not produce appreciable rescue of RPE or photoreceptors in the KO screen. In a separate experiment, we sought to test the hypothesis that dsRNA could be produced from 5’ and 3’ AAV inverted terminal repeat (ITR)-directed sense/antisense transcripts, which could hybridize to induce dsRNA-dependent toxicity.^30,81^ An AAV constructed to test this possibility did not produce appreciable toxicity. Conversely, subretinal administration/electroporation of poly(I:C) produced significant toxicity in pups injected and harvested at the same age. This argues against a purely ITR-driven dsRNA-based toxicity mechanism but shows that neonatal mice are capable of mounting a response to dsRNA.

Expression of IFNAR1 and IFNB1 in both RPE and retina was observed using FISH assays. Greater IFNB1 expression was seen in cell types expressing GFP, as more IFNb signal was detected in RPEs of Best1::GFP-infected animals or retinas of Rho::GFP-infected animals than in uninfected animals. This suggests that AAV-induced interferon expression is stimulated most strongly in cells overexpressing a transgene, which has also been suggested by previous studies.^59^ However, as discussed above, for toxicity and cell death to be initiated, primarily in the RPE and secondarily in photoreceptors, stress from transgene overexpression in the RPE likely needs to be present, which is not the case for Rho::GFP. The model proposed for the generation of AAV-induced ocular toxicity by the Best1::GFP vector, and possibly other toxic, transgene-expressing AAVs, is shown in Figure 6.

**Figure 6:**
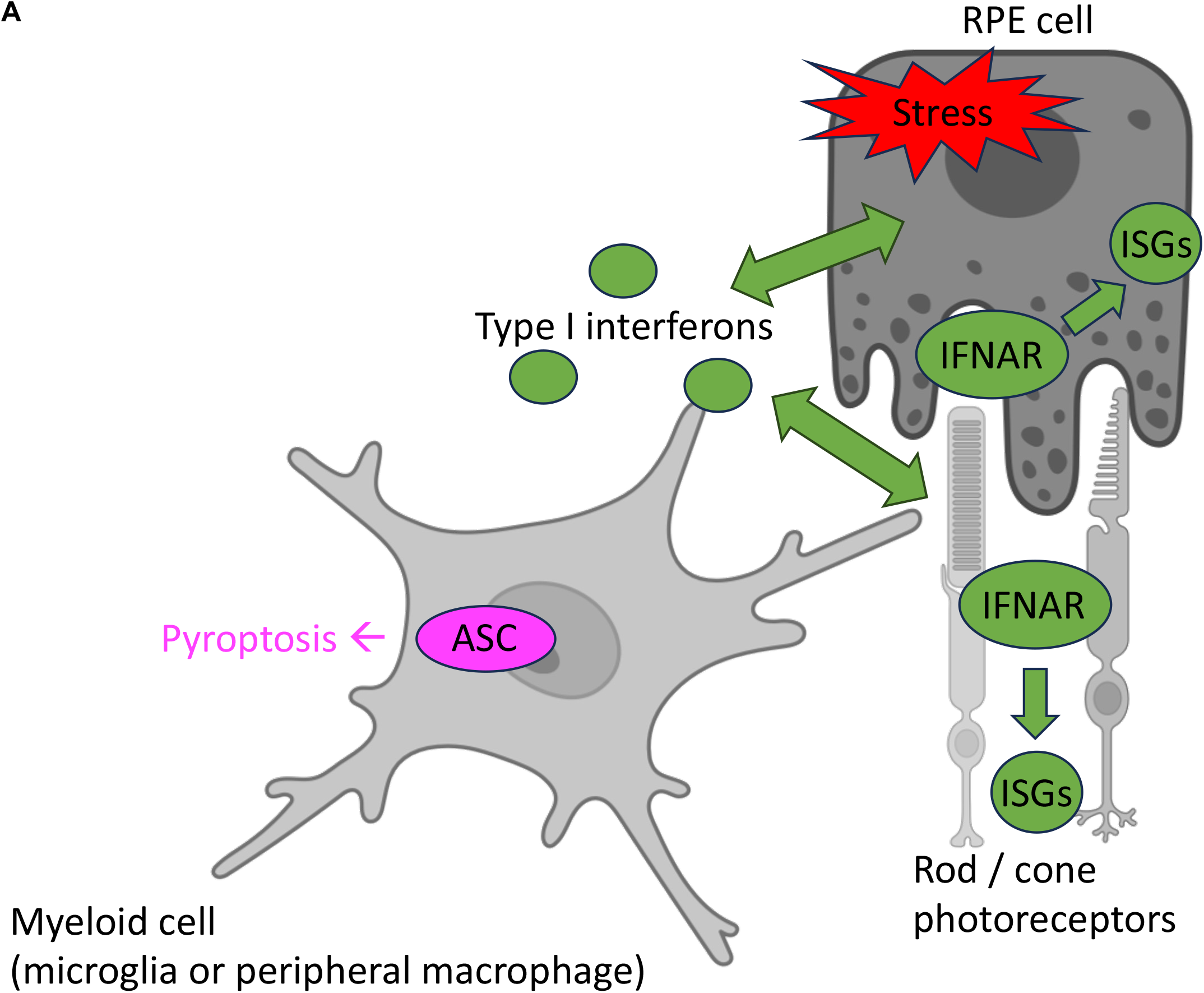
Model for how induced ocular toxicity is generated. A) Following subretinal injection of AAV, which infects both RPE and photoreceptor cells, cell stress responses in the RPE arise in response to expression of specific transgenes and/or their expression levels. The stress response in the RPE kills them and leads to secondary photoreceptor death. Maladaptive type I interferon responses, possibly triggered by cGAS sensing of cytosolic double-stranded DNA, contribute to retinal toxicity. Infected RPE and photoreceptor cells respond to infection by expressing and secreting type I interferon molecules, e.g., IFNb. Type I IFN molecules can activate the expression of interferon stimulated genes (ISGs) in cells that express the Type I IFN receptor (IFNAR), including RPE, photoreceptors, and possibly myeloid cells. Myeloid cells include microglia and peripheral infiltrating macrophages. Myeloid cells are recruited to the ONL and RPE layers in response to toxic AAV injection and may participate in interferon signaling. Myeloid cells also undergo pyroptosis as visualized by ASC speck formation, which is indicative of inflammasome assembly. However, pyroptosis itself does not contribute to the observed toxicity. Temporally, both stress responses and induction of certain ISGs occur by week 1 according to RNA-seq data from the RPE, suggesting that these responses may occur simultaneously. Myeloid cell recruitment to the ONL/RPE likely follows these responses. Figure created with BioRender.

A key observation is that the overall severity of AAV-associated toxicity following neonatal subretinal injections is largely transgene-dependent, rather than preparation method, capsid, or genome dependent. We replaced GFP with other transgenes, such as RPE65, and found no measurable toxicity. In addition, AAVs with considerably less or no measurable toxicity were created when using promoters that limit or abolish transgene expression in the RPE. The Rho, RK, CAR, and RedO promoters have been used for this purpose.^12^ Lastly, a control vector, Best1::6xSTOPmutGFP, constructed to have 2 tandem STOP codons in each reading frame (6xSTOP), followed by a GFP sequence with 41 point mutations that abolished any ORF >73 amino acids, was found to be benign (Figure 5A).^43^ While valuable for understanding toxicity mechanisms, these approaches are not necessarily clinically generalizable for making AAVs less toxic, as RPE-specific expression of therapeutic transgenes may be required for certain diseases. However, these results do imply that transgene expression should be limited to the target cell type of interest, whenever possible, using capsid, promoter, and/or other regulatory mechanisms.

GFP toxicity in the eye is likely due to properties inherent to excessive GFP protein, rather than GFP RNA, as the Best1::6xSTOPmutGFP vector is benign despite being able to produce a mutated, non-translatable GFP RNA roughly ∼94% identical to the wild type GFP sequence. It is difficult to predict the toxicity of any particular transgene, as our small survey of 5 transgenes did not yield strong correlations. The lack of toxicity from an AAV expressing CasMini in the RPE suggests that transgene toxicity is not governed by self vs. non-self. The level of expression may play a role in determining transgene toxicity, as one correlation for GFP toxicity is the dose delivered to the eye.^11,82^ Similarly, the tragic cases of AAV-induced death also followed injections of high doses.^16–19^

Toxicity was evaluated not only in neonatal mice, but also in adult mice via subretinal injections. It was not clear what dosing to use to evaluate the relative toxicity at these two ages. Adult eyes are larger, with about twofold more cells, mostly in the photoreceptor layer. The greater retinal thickness and maturity of the adult eye may provide greater resilience to procedural trauma. The anatomy also differs in that the outer segments and RPE processes are more intimate and thus an injection subretinally tends to create a detachment. In neonates, due to lack of outer segments and RPE apical processes, injections seem to be much less damaging, and the younger eye also tends to heal more quickly. The inoculum remains as a bleb in adults,whereas it flows throughout the subretinal space in neonates. Best1::GFP injection still produced toxicity in adults, although the sensitivity to viral toxicity was lower in adults (Figure S16-S17). A 2e10 gc (in ∼1 uL injection volume) Best1::GFP dose in adults gave less severe RPE toxicity than the 4e9 gc (in ∼0.2 uL injection volume) Best1::GFP dose in neonates (see the toxicity data in Figure S16). The 2e10 gc dose of Best1::GFP in adults did, however, give comparable toxicity to the neonatal dose of 4e8 gc. Similarly, a 2e9 gc CMV::GFP dose in adults gave less severe RPE toxicity than the 4e8 gc CMV::GFP dose in neonates (Figure S17). Lower cone toxicity accompanied lower RPE toxicity in the context of Best1::GFP, but not CMV::GFP. In addition to the structural differences between the two stages, toxicity may be blunted in adult mice due to age-dependent biological susceptibility. Adult mice likely have more complex antiviral responses against AAV vectors injected subretinally than neonates, which could reduce ocular toxicity.

Overall, future work on AAV toxicity should involve testing a variety of clinically relevant transgenes using approaches similar to those in this work to determine whether the mechanisms of toxicity implicated here (Figure 6) apply in other cases. Finding ways to inhibit, genetically or pharmacologically, pathways implicated in toxicity can then be leveraged to mitigate AAV-induced toxicity and create safer AAV gene therapies.

## Materials and Methods

### Mice

Mice were purchased from the Jackson Laboratories (JAX) and maintained at Harvard Medical School on a 12-hour alternating light/dark cycle. All experiments were approved by the Institutional Animal Care and Use Committee (IACUC) of Harvard University and performed in accordance with institutional guidelines. The following mouse strains were used in this study: C57BL/6J (JAX #000664), 129S1/SvImJ (JAX #002448), MAVS KO (JAX #008634), TNFRp55/Tnfrsf1a/TNFR1 KO (JAX #002818, Sv129 background, or JAX #003242, C57BL/6 background^41^), IFNAR1 KO (JAX #028288), IFNGR1 KO (JAX #003288), C3 KO (JAX #029661), MyD88 KO (JAX #009088), Ticam1 KO (#JAX 005037), Rag1 KO (JAX #002216), IL2Rγ KO (JAX #003174), CHOP KO (JAX #005530), cGAS KO (JAX #026554), Cx3cr1-GFP (JAX #005582), Cx3cr1-GFP, Ccr2-RFP (JAX #032127), Ai75D (JAX #025106), and GSDMD KO (JAX #032410). The MyD88/Ticam1 double KO mouse was generated by crossing the respective single KO strains together in-house. Sometimes, Ai75D heterozygous animals were used in place of C57BL/6J for experiments due to animal availability and timing of litters. The equivalence of the C57BL/6J strain and the Ai75D heterozygote animals is shown in Figure S18 (injected with 4e8 gc Best1::GFP, RPE scores and cone counts) and Figure S9A (uninjected cone counts). Uninjected RPE flatmounts from both strains were comparably healthy. Genotyping was performed by Transnetyx.

### AAV Production/Titering

The Best1::GFP and Rho::GFP vector (version 1 of 2, see note below) were cloned from the previously published CMV::EGFP vector^12^, which was obtained from the Harvard DF/HCC DNA Resource Core (clone ID: EvNO00061595) and consisted of a CMV promoter, b-globin intron, cDNA sequence for EGFP, and b-globin polyadenylation sequence. Best1 vectors with new transgenes were designed from NCBI RefSeq or available Addgene sequences, purchased as IDT gBlocks, and replaced EGFP by Gibson Assembly. Promoters and introns were similarly cloned into the CMV::EGFP backbone by Gibson Assembly.

Two versions of the Rho::GFP vector were used in this study. A map of version 1 is shown in Figure 1A and is used for Figures 1-2, 4A, S3-S5, and the Rho::GFP injected control samples in Figure S9. Version 2 of Rho::GFP was used previously by our lab and consists of the following genome sequence:

### 5’ ITR — Rho promoter — EGFP — WPRE element — BGH poly A sequence — 3’ ITR.^12^

This vector was used for the HCR experiments in Figure 4B-E.

Recombinant AAV2/8 vectors were produced by transfecting 80% confluent HEK293T cells with the AAV vector, AAV8 rep/cap plasmid, and adenoviral helper plasmid mixed with polyethylenimine.^42^ A few hours before transfection, media was replaced with DMEM + 10% Nuserum + 1% penicillin G/streptomycin, which was then replaced with DMEM 16-24 hours post-transfection. 72 hours post-transfection, supernatant was collected and AAV was precipitated by mixing with 8% wt/vol PEG-8000 + 0.5 M NaCl and leaving on ice at 4°C overnight, then centrifuging for 30 minutes at 7,000 x g. The PEGylated pellet was resuspended in lysis buffer (150 mM NaCl + 20 mM Tris, pH 8), filtered through a 0.45 micron filter, and ultracentrifuged through a previously described iodixanol gradient^42^ in a Beckman VTi 50 rotor at 46,500 rpm (∼200,000 x g) for 2 hours at 16°C. The 40% iodixanol fraction was collected into an Amicon 100k 15 mL filter tube (MilliporeSigma #UFC910008), and buffer exchanged with PBS three times for a final volume of 75-150 uL. AAV was titered by SYPRO Ruby Protein Gel Stain (Thermo Fisher Scientific #S-12000) relative to a reference vector titered by qPCR. AAV was stored in 10 uL aliquots at -80°C. For subretinal injection, AAV was thawed at RT, then placed on ice.

### Subretinal AAV injections

Glass needles for subretinal injection were pulled and sharpened as previously described.^43^ AAV was delivered by subretinal injection into P0-P2 mouse pups or adult mice >12 weeks old (see legends for more specific ages of injected cohorts) as previously described.^43^ AAV solutions contained 0.01% wt/vol Fast Green FCF dye to visualize the injection bleb. Doses of each AAV used for injections are reported in genome copy (gc) units.

Example dilution calculation for pups: AAVs administered at a 4e8 gc injection dose were diluted to 1.6e9 gc/uL in PBS/Fast Green dye, and ∼0.25 uL was injected per eye.

Example dilution calculation for adults: AAVs administered at a 2e9 gc injection dose were diluted to 2e9 gc/uL in PBS/Fast Green dye, and ∼1-2 uL was injected per eye.

### Histology

Mice were euthanized by CO_2_ overdose followed by cervical dislocation. Eyes were removed and dissections/immunofluorescence staining steps were carried out as previously described for mouse eyes.^43^ Staining buffer (used to dilute primary antibodies) consisted of 1% Triton X-100 and 4% normal donkey serum in PBS and was prepared fresh before each use. Secondary antibodies (if applicable) were diluted in PBS.

Primary antibodies/lectins used in this study include: Alexa Fluor 647 Phalloidin (1:100, Invitrogen #A22287), Alexa Fluor 568 Phalloidin (1:100, Invitrogen #A12380), cone arrestin Rb polyclonal (1:3000, Millipore Sigma #AB15282), IbaI Rb polyclonal (1:200, GeneTex, #GTX100042), and ASC/TMS1 Rb monoclonal (1:1000, Cell Signaling Technology, #67824). Secondary antibodies (all used at a 1:1000 dilution) in this study include: Alexa Fluor® 647 Donkey Anti-Rabbit IgG (Jackson ImmunoResearch, #711-605-152) and Dylight^TM^ 405 AffiniPure® Donkey Anti-Rabbit IgG (Jackson ImmunoResearch, #711-475-152).

For whole eyecup section analysis, sections containing the optic nerve were used for analysis, as the optic nerve served as a landmark for identifying regions in the mid-periphery to count the number of ONL layers for quantification (∼1 mm from optic nerve head). In cases where the optic nerve fell off or was missing, centrally located sections were taken for analysis.

After secondary antibody staining (if applicable) and 3x washes in PBS, RPE flatmounts were mounted RPE-side down onto coverslips (VWR, 24×50mm No. 1.5, #48393-241) in Fluoromount G mounting media (Southern Biotech) on Superfrost Plus slides (Fisher Scientific, #22-037-246) and subsequently imaged. Retinal flatmounts were similarly mounted (ONL-side down) prior to imaging. Sections were stained with DAPI and coverslipped in Fluoromount G mounting media prior to imaging.

### RNA-sequencing

For Figures 2A-C, 4A, Figures S3-S5, and Tables S1-S7, B6J mice were subretinally injected with 4e8 gc Best1::EGFP in one eye and 4e8 gc Rho::EGFP (version 1 vector map in Figure 1A) in the contralateral eye. Eyes were harvested at 1 and 2 weeks post-transduction (n=5). The neural retina was separated from the RPE-choroid-sclera. Upon separation, the RPE-choroid-sclera was incubated for 10-15 minutes in 200 uL of RNAprotect Cell Reagent (QIAGEN #76526) to promote the separation of RPE cells from the choroid-sclera. After incubation, the samples were centrifuged at 200 x g for 5 minutes. The scleral layer was then removed and discarded. Samples were stored at -20°C until ready for processing. For processing, samples were sent to the Biopolymers Facility at Harvard Medical School for RNA purification by RNeasy Micro Kit, analysis on an Agilent TapeStation, library prep, and 75 bp paired-end sequencing on an Illumina NovaSeq 6000 S1 for 200 cycles.

The resulting data was aligned to Ensembl’s *Mus musculus* GRCm39 release 112 by STAR^44^ 2-pass mapping. Results were analyzed by FastQC^45^, further processed by Picard Tools, and re-analyzed by Qualimap.^46^ Reads were counted using featureCounts^47^ for paired end datasets, counting at the gene (meta-feature) level, counting only reads mapping to exons, allowing reads to overlap multiple genes, and excluding chimeric fragments or multimapping reads. Overall metrics were aggregated by MultiQC for review.^48^ The featureCounts results for differential gene expression were analyzed in R. All 20 samples were tidied together, reviewed by PCA, and then analyzed for differential gene expression by DESeq2.^49^ Read counts were normalized using DESeq2’s pipeline, which uses the median of ratios method.^50^ Since by PCA, the timepoint had the greatest contribution to variability within the dataset, differential expression (DE) analysis was ultimately performed by timepoint, such that DE was evaluated between the Week 1 Rho vs. Best1 samples, and then performed again for the Week 2 Rho vs. Best1 samples. Since multiple groups were present in this analysis, shrinkage of the log2 fold-change estimates was performed by ashr instead of apeglm. GO enrichment and GSEA for KEGG pathways were performed using clusterProfiler.^51^

For the RNA-seq data in Figure S6A, mice injected with the indicated AAVs at birth were harvested at 2 weeks. RPEs and retinas were dissected and immediately placed in 55 uL of DNA/RNA Shield (Zymo #R1100-50) and incubated overnight at room temperature. The next day, RPE samples were flicked to enable the RPE cells to fall off the eyecup, the eyecup was removed, and the RNA was submitted to Plasmidsaurus for bulk RPE RNA-sequencing. Analysis was also performed by Plasmidsaurus.

### HCR RNA-FISH

HCR RNA-FISH was performed to detect immune-related transcripts on RPE and retinal flatmounts. Probes were manufactured by Molecular Instruments (M.I.) to target the mouse cDNA of the indicated gene. RPE and retinal flatmounts were fixed in 4% paraformaldehyde in PBS (Thermo Scientific, #28908) overnight at 4°C, then washed 3x in PBS (5 minutes/wash), then permeabilized in 70% ethanol/PBS for 2-4 hours at room temperature. Samples were then washed 1x in PBS, then incubated for 5 minutes in probe wash buffer at room temperature. Samples were then incubated in probe hybridization buffer for 30 minutes at 37°C, then hybridized with the desired probes diluted in probe hybridization buffer (6 uL M.I. probe per 1 mL probe hybridization buffer) overnight at 37°C.

The next day, probe hybridization solutions were removed and samples were processed at room temperature for the rest of the protocol. Samples were washed in probe wash buffer twice (30 minutes/wash), then washed in 5X SSCT in PBS with 0.1% Tween-20 twice (20 minutes/wash), then washed once in probe amplification buffer for 10 minutes (this buffer was pre-warmed to room temperature).

In parallel with these wash steps, HCR amplifiers (8 uL of hairpin 1 and 8 uL of hairpin 2 amplifiers for every 1 mL of amplification buffer) were prepared by heating h1 and h2 hairpins separately at 95°C for 90 seconds, followed by cooling for 30+ minutes in a dark drawer at room temperature. After cooling, HCR amplifiers were diluted in amplification buffer (prewarmed to room temp).

When samples finished incubating in (plain) amplification buffer for 10 minutes, the buffer was removed and replaced with the amplification buffer containing the fluorescent h1 and h2 amplifier hairpins. Samples were left overnight at room temperature for amplification, covered in foil.

The next day, samples were washed twice in 5X SSCT in PBS with 0.1% Tween-20 twice (20-30 minutes/wash), then washed once in PBS. Retinas were then ready for mounting/imaging, unless additional antibody staining was desired (e.g., CAR staining to identify the ONL region in flatmount images). In this case, primary antibodies diluted in staining buffer were immediately added, and the rest of the antibody staining and flatmounting protocol follows the methods outlined above (see section on Histology).

### Image Acquisition and Analysis

Tissues were imaged using one or more of the following microscopes from the Harvard MicRoN core: a point scanning confocal microscope (for some sections), a spinning disk confocal microscope (for sections, RPE flatmounts, and retinal flatmounts), or a widefield slide scanner microscope (for sections, RPE flatmounts, and retinal flatmounts). Details on each system are below:

- A Zeiss LSM 710 and LSM 780 single point scanning confocal microscope was used. The 20x air, 25x oil, and 63x oil objectives and the 405, 488, 561, and 633 nm laser lines were used.
- A Nikon Ti or Nikon Ti2 inverted microscope (W1 Yokogawa spinning disk confocal microscopes) were used. The 10x air, 20x air, or 40x oil objectives and the 405, 488, 561, and 640 nm laser lines were used.
- An Olympus VS200 slide scanner with a UPlan X Apo 10x/0.4 Air objective was used. The DAPI, FITC, mCherry, and Cy5 channels were used.

Image analysis was performed using ImageJ/Fiji or OlyVIA. OlyVIA was only used to analyze ONL layers from images taken using the slide scanner. Since ONL layer quantification can vary by location, counts were always obtained from regions in the mid-periphery, defined as ∼1 mm away from the optic nerve head on one or both sides of each section. If ONL layer measurements were taken on both sides of the optic nerve head, the median of the layer counts was reported. If tissue on one side of a section was damaged (e.g., due to a technical/sectioning issue), then only the undamaged side’s measurements were reported for that sample.

Flatmount quantification was performed as described previously, with a few modifications.^43^ For RPE scoring, up to four fields of view of identical size (e.g., 250 µm^2^) in each flatmount were chosen randomly around the flatmount for analysis. The semiquantitative RPE toxicity grading scale established in a previous publication from our lab^12^ was applied to each field of view for each RPE flatmount by an investigator blinded to the treatment conditions. The median toxicity score was plotted. If a region was damaged due to technical regions, out of focus, or not well stained, it was excluded from scoring.

For cone counting, four 250 µm^2^ boxes were placed at the midpoints of 4 radial lines drawn from the outer edge of each retinal flatmount leaflet to the center of the flatmount, as previously described.^43^ Care was taken to avoid areas of tissue damage or uneven staining. An automated ImageJ/Fiji algorithm counted the number of CAR+ cone puncta in each box, and the median cone count was reported. Retinal flatmount boxes were excluded from analysis if uneven staining issues, tissue damage, or technical focusing/imaging issues in a box impeded algorithmic detection of CAR+ puncta.

For microglia counting, four 500 µm^2^ images were taken around the RPE or retinal flatmount, and microglia in each region was manually counted. The median microglia count for each sample was plotted.

Unless otherwise noted, for neonatal AAV subretinal injections, only Best1::GFP RPE flatmounts and corresponding retinal flatmounts with >90% coverage of the eyecup, judged by GFP+ signal on RPE flatmounts, were analyzed to reduce variability in injection technique. Samples that were not well stained or were damaged due to technical issues (e.g., dissection-related damage) were excluded from analysis.

For subretinal AAV injections into adult mice, the transduction coverage varies but is usually <50% of the total eyecup surface area, so images were not excluded on the basis of lack of full coverage as for neonates. Instead, quantification was done in transduced areas and/or untransduced areas separately by drawing boxes in areas with evidence of GFP+ signal, and reported as such (see figure legends for more information).

### Statistical Analysis

Outside of RNA-seq, statistical analyses were performed using GraphPad Prism software (Dotmatics). Data are represented as mean ± standard deviation. For all figures, n refers to individual eyes. Comparisons of two groups were performed using an unpaired Student’s t-test. In cases where multiple t-tests are shown on the same plot, a multiple comparisons correction was usually implemented (and indicated in the legend). Unless otherwise noted, comparisons of three or more groups were performed using a one-way ANOVA with a multiple comparisons test. For all tests, a p-value less than 0.05 was considered significant; p-values above 0.05 were reported as not significant or n.s.

## Supporting information

Tables Gardner et al. 2026

Gardner et al. Supp figures and Table and FIgure legends

## Data Availability Statement

Raw data supporting the findings of this study can be made available upon request. Plasmids have been deposited at Addgene. RNA-sequencing data has been deposited to GEO (accession numbers GSE338362 and GSE338363).

## Acknowledgments

The authors would like to thank the Harvard Medical School MicRoN core for their help with microscopy and the Biopolymers Facility at Harvard Medical School for discussions and technical support. The authors would like to thank Heer Joisher, Kanae Sasaki, and Anjali Vlassak for insightful discussions and help with the study. The authors would also like to thank the Isaac Chiu lab at Harvard Medical School for generously sharing the GSDMD KO mouse strain used in this study. This study was funded by the Howard Hughes Medical Institute and the National Eye Institute (R01 EY029348-01A1 awarded to CLC and F31 5F31EY035867-02 awarded to AG).

## Author Contributions

AG, CMH, and CLC designed the study. AG, CMH, and SRZ performed experiments. AG, CMH, AD, and CLC analyzed data. AG, CMH, and CLC wrote the manuscript.

