## Supplementary material for "Host Endoplasmic Reticulum Stress and Interferon Responses Contribute to AAV-Induced Ocular Toxicity": Gardner et al. Supp figures and Table and FIgure legends

**Supplemental Figure Legends**

**Figure S1. Representative wild type C57BL/6J mouse RPE flatmount and eyecup sections.**

A) Top: Representative RPE flatmount from a wild type C57BL/6J mouse at P30. RPE flatmount is stained with phalloidin (red). Scale bar: 100 µm.

Bottom: RPE semiquantitative scoring method.

B) Representative eyecup section from a wild type C57BL/6J mouse at P28. Section is stained with DAPI (blue). Scale bar: 50 µm. To count the number of ONL layers, 3 lines were drawn spanning the ONL and the number of nuclei layers was counted. In this example, the left orange line spans ~12 layers, the middle line spans ~13 layers, and the right line spans ~12 layers. The median of these values was reported for the sample, which in this case is ~12 layers.

**Figure S2. Dose response characterization of Best1::GFP.** All mice in this figure were harvested at 4 weeks of age. The uninjected C57BL/6J mouse control image is the same as the one shown in Figure S1B.

A) Top row: Representative eyecup section from an uninjected C57BL/6J mouse (left) or from C57BL/6J mice injected with 2e7 gc Best1::GFP (middle) or 2e8 gc Best1::GFP (right). Bottom row: Representative eyecup sections from C57BL/6J mice injected with 4e8 gc Best1::GFP (left) or 2e9 gc Best1::GFP (right). Sections are stained with DAPI (blue) and the GFP channel is overlaid. Scale bar: 50 µm.

B) Median number of ONL layers remaining from samples injected with the indicated dose of Best1::GFP. Uninjected eyes serve as age-matched controls (n=4-17 eyes per group, ****$p<0.0001$, mean ± SD, one-way ANOVA with Bonferroni multiple comparisons correction). Uninjected control samples are the same as shown in Figure 1.

**Figure S3. Principal components analysis on bulk RNA-sequencing data from AAV-infected RPE.** This dataset is the same as the one described in Figure 2. P0 pups were subretinally injected with 4e8 gc of either Rho::EGFP or Best1::EGFP. Enriched RPE cells were collected at 1 week and 2 weeks post-transduction for bulk RNA-seq (n=5).

A) PC1-2 (36% and 14% of variance) after PCA of normalized counts for all genes.

B) PC2-3 (14% and 9% of variance) after PCA of normalized counts for all genes.

C) The cumulative amount of variance captured by the addition of each principal component from PCA.

**Figure S4. Differential gene expression analysis for RPE samples injected with Rho::EGFP vs. Best1::EGFP.** Differential expression analysis was performed by DESeq2. This dataset is the same as the one described in Figure 2. P0 pups were subretinally injected with 4e8 gc of either Rho::EGFP or Best1::EGFP. Enriched RPE cells were collected at 1 week and 2 weeks post-transduction for bulk RNA-seq (n=5).

A) Heatmap of differentially expressed genes. For this heatmap, the thresholds for differential expression were an adjusted p-value $< 0.05$ and a log2 fold-change $> 1$.

**Figure S5. GO terms for downregulated differentially expressed genes from bulk RPE RNA-sequencing dataset.** This dataset is the same as the one described in Figure 2. P0 pups were subretinally injected with 4e8 gc of either Rho::EGFP or Best1::EGFP. Enriched RPE cells were collected at 1 week and 2 weeks post-transduction for bulk RNA-seq (n=5).

A) Tree plots of the top 25 Gene Ontology Biological Process terms associated with downregulated DE genes at week 1 (top) and week 2 (bottom). Pathways are downregulated in the Best1::GFP relative to the Rho::GFP infection context.

**Figure S6: Survey of immune related gene abundances in unstimulated microglia.** Data from panel A was generated in this study. Data from panel B was previously published and represents unstimulated microglial immune-related counts per million reads (CPMs).[1]

(A) CPM for selected immune-related genes at 2 weeks of age. Mice were injected with 4e8 gc Best1::GFP or uninjected. Genes were selected on the basis of being an AAV transgene (EGFP), a housekeeping or visual cycle gene, an immune related receptor/sensor, a cell stress-related gene, or an interferon stimulated gene.

B) Plot of immune-related transcript CPMs. Cx3cr1 and Tmem119 are included as reference microglial marker genes.

**Figure S7: Representative RPE flatmounts from the KO mouse screen.** All mice were harvested 4-5 weeks post infection. Each image represents an individual eye. Scale bar: 20 μm.

A) RPE flatmounts from uninjected C57BL/6J mice (n=3 shown). 16 regions of 4 RPE flatmounts from 2 animals were scored, with the average score = 0.

B) RPE flatmounts from C57BL/6J mice injected with 4e8 gc Best1::GFP (n=9 shown). 38 regions of 16 RPE flatmounts from 10 animals were scored, with the average score = 2.2.

C) RPE flatmounts from IFNAR1 KO mice injected with 4e8 gc Best1::GFP (n=9 shown). 22 regions of 9 RPE flatmounts from 8 animals were scored, with the average score = 1.9.

D) RPE flatmounts from IFNGR1 KO mice injected with 4e8 gc Best1::GFP (n=4 shown). 15 regions of 4 RPE flatmounts from 3 animals were scored, with the average score = 2.8.

E) RPE flatmounts from CGAS KO mice injected with 4e8 gc Best1::GFP (n=9 shown). 30 regions of 9 RPE flatmounts from 7 animals were scored, with the average score = 1.7.

F) RPE flatmounts from MAVS KO mice injected with 4e8 gc Best1::GFP (n=4 shown). 9 regions of 4 RPE flatmounts from 2 animals were scored, with the average score = 2.3.

G) RPE flatmounts from RAG1 KO mice injected with 4e8 gc Best1::GFP (n=8 shown). 25 regions of 8 RPE flatmounts from 5 animals were scored, with the average score = 2.4.

H) RPE flatmounts from IL2RG KO mice injected with 4e8 gc Best1::GFP (n=12 shown). 38

regions of 12 RPE flatmounts from 8 animals were scored, with the average score = 2.2.

I) RPE flatmounts from TNFR1 KO mice injected with 4e8 gc Best1::GFP (n=10 shown). 30 regions of 10 RPE flatmounts from 6 animals were scored, with the average score = 2.

J) RPE flatmounts from C3 KO mice injected with 4e8 gc Best1::GFP (n=3 shown). 12 regions of 3 RPE flatmounts from 3 animals were scored, with the average score = 2.6.

K) RPE flatmounts from CHOP KO mice injected with 4e8 gc Best1::GFP (n=3 shown). 6 regions of 3 RPE flatmounts from 3 animals were scored, with the average score = 1.1.

L) RPE flatmounts from TLR double KO mice injected with 4e8 gc Best1::GFP (n=5 shown). 19 regions of 5 RPE flatmounts from 5 animals were scored, with the average score = 2.4.

**Figure S8: Assessment of IFNAR KO strain on CMV::GFP toxicity phenotype.** All mice in this figure were harvested at 4-5 weeks of age.

A) RPE and retinal flatmounts were harvested from B6J or IFNAR KO mice injected at birth with 4e8 gc CMV::GFP. The RPE toxicity scores (n=4-6 eyes per group, mean ± SD, unpaired t-test) and median cone counts (n=4-8 eyes per group, *p=0.02, mean ± SD, unpaired t-test) are plotted. Uninjected control samples are the same as shown in Figure 3.

**Figure S9: Baseline RPE, cone, and ONL layer quantifications for the mouse strains used in this paper.** All mice were 1-2 months of age at harvest.

A) Quantification of cone cell counts (n≥3 eyes per group, mean ± SD, *p=0.03, one-way ANOVA with Dunnett's multiple comparisons correction). All samples quantified in this plot are uninjected. All mice were 1-2 months of age at harvest. B6J uninjected control samples are the same as shown in Figure 3.

B) Median number of ONL layers (n≥3 eyes per group, *p=0.01, mean ± SD, one-way ANOVA with Dunnett's multiple comparisons correction). Unless labeled as (uninj) for uninjected samples on the plot, all samples were injected with 4e8 gc Rho::GFP. B6J uninjected control samples are the same as shown in Figure 3.

**Figure S10: Pharmacologic depletion of microglia to assess the role of microglia in AAV-associated ocular toxicity.** For panels B-D, the Best1::GFP AAV dose was 4e8 gc. For panels E-G, the Best1::GFP AAV dose was 4e9 gc.

A) Experimental workflow for depleting microglia in pups. Neonatal mice were subretinally injected with Best1::GFP at a dose of 4e8 or 4e9 gc. At 1 week of age, the lactating mother was given chow with 1200 ppm PLX5622 (or fed normal chow as a control) incorporated to initiate microglial depletion. The dam and her pups remained on the chow until P28, when the pups were harvested.

B) RPE toxicity scoring (n≥4 eyes per group, mean ± SD, **p=0.007, unpaired t-test). Best1-GFP (4e8 gc) and uninjected control samples are the same as shown in Figure 3.

C) Quantification of cone counts (n≥4 eyes per group, mean ± SD, unpaired t-test). Best1-GFP (4e8 gc) and uninjected control samples are the same as shown in Figure 3.

D) Median number of ONL layers from sectioning retinal flatmounts (n≥3 eyes per group, mean ± SD, unpaired t-test). Uninjected control samples are the same as shown in Figure 3.

E) RPE toxicity scoring (n≥4 eyes per group, mean ± SD, **p=0.007, unpaired t-test). Uninjected control samples are the same as shown in Figure 3.

F) Quantification of cone counts (n≥4 eyes per group, *p=0.02, mean ± SD, unpaired t-test). Uninjected control samples are the same as shown in Figure 3.

G) Median number of ONL layers from sectioning retinal flatmounts (n≥4 eyes per group, mean ± SD, unpaired t-test). Uninjected control samples are the same as shown in Figure 3.

**Figure S11: Quantification of pharmacologic depletion of microglia (AAV Best1::GFP dose: 4e8 gc).** Injections were performed in mice at birth.

A) P28 representative RPE flatmounts stained with IbaI (magenta) to label microglia. These mice were fed regular chow throughout the experiment. Scale bar: 100 μm.

B) P28 representative RPE flatmounts stained with IbaI (magenta) to label microglia. These mice were fed PLX5622-containing chow from P7-P28. Scale bar: 100 μm.

C) Analysis of IbaI+ depletion efficiency. The number of IbaI+ cells were counted in 3-4 boxes (500 $\mu m^2$) around each RPE flatmount and the median count was plotted (n≥14 eyes per group, mean ± SD, ***p=0.0002, unpaired t-test). Efficiency was ~91% for this experiment.

**Figure S12: Quantification of pharmacologic depletion of microglia (AAV Best1::GFP dose: 4e9 gc).** Injections were performed in mice at birth.

A) P28 representative RPE flatmounts stained with IbaI (magenta) to label microglia. These mice were fed regular chow throughout the experiment. Scale bar: 100 μm.

B) P28 representative RPE flatmounts stained with IbaI (magenta) to label microglia. These mice were fed PLX5622-containing chow from P7-P28. Scale bar: 100 μm.

C) Analysis of IbaI+ depletion efficiency. The number of IbaI+ cells were counted in 3-4 boxes (500 $\mu m^2$) around each RPE flatmount and the median count was plotted (n≥10 eyes per group, mean ± SD, ****p<0.0001, unpaired t-test). Efficiency was ~98% for this experiment.

**Figure S13: Quantification of pharmacologic depletion of microglia in uninjected animals.** For all images in this figure, microglia are labeled in green.

A) Representative retinal flatmounts from P19 Cx3cr1-GFP heterozygous reporter mice. The lactating dam and pups were given access to PLX5622-containing chow from P8-P19.

B) Representative retinal flatmounts from P19 Cx3cr1-GFP heterozygous reporter mice. The lactating dam and pups were given access to regular chow from P8-P19.

C) Representative retinal flatmounts from P38 Cx3cr1-GFP heterozygous reporter mice. The mice were given access to PLX5622-containing chow from P31-P38.

D) Representative retinal flatmounts from P38 Cx3cr1-GFP heterozygous reporter mice. The mice were fed regular chow throughout their life.

E) Quantification of the images presented in panels A-D (n=4 eyes per group, *p<0.0001, mean ± SD, multiple unpaired t-tests with Bonferroni-Dunn multiple comparisons correction).

**Figure S14: Assessment of myeloid cell pyroptosis on AAV-associated ocular toxicity.** For all bar plots in this figure, the B6J controls (either injected with 4e8 gc Best1::GFP or uninjected) were previously plotted in Figure 3.

A) C57BL/6J mice were injected with 4e8 gc Best1::6xSTOP-mutGFP (left), 4e8 gc Best1::GFP (middle), or 4e9 gc Best1::GFP (right) and harvested at 4-5 weeks of age. Retinal flatmounts were stained for ASC (magenta). Scale bar: 100 µm.

B) Cx3cr1-GFP heterozygous reporter mice were injected with 4e8 gc CMV::null and harvested at P19. Retinal flatmounts were stained for ASC. Left image is the ASC channel (magenta), middle image is the Cx3cr1-GFP channel (green), and the right image shows both channels merged (colocalization of signals is depicted in white). Scale bar: 100 µm.

C) GSDMD KO mice or C57BL/6J controls were injected at birth with 4e8 gc Best1::GFP and harvested 4 weeks post injection. RPE flatmounts (n=16 B6J eyes, n=8 GSDMD KO eyes, mean ± SD, ****p<0.0001), retinal flatmounts (n=22 B6J eyes, n=8 GSDMD KO eyes, mean ± SD, unpaired t-test), and the number of ONL layers (n=17 B6J eyes, n=7 GSDMD KO eyes, mean ± SD, unpaired t-test) were quantified.

D) Uninjected GSDMD KO mice or C57BL/6J controls were harvested 4 weeks post injection. RPE flatmounts (n=4 B6J eyes, n=5 GSDMD KO eyes, mean ± SD, unpaired t-test), retinal flatmounts (n=4 B6J eyes, n=7 GSDMD KO eyes, mean ± SD, unpaired t-test), and the number of ONL layers (n=4 B6J eyes, n=3 GSDMD KO eyes, mean ± SD, unpaired t-test) were quantified.

**Figure S15: Assessment of chemokine receptors on AAV-associated ocular toxicity.**

A) CX3CR1/CCR2 double heterozygous, KO, or WT mice were injected at birth with 4e8 gc Best1::GFP and harvested 4 weeks post injection. RPE flatmounts (n=4-6 eyes per group, mean ± SD, one-way ANOVA with Dunnett's multiple comparisons test), retinal flatmounts (n=3-5 eyes per group, mean ± SD, one-way ANOVA with Dunnett's multiple comparisons correction), and the number of ONL layers (n=5-6 eyes per group, mean ± SD, unpaired t-test) were quantified.

**Figure S16: Effect of Best1::GFP expression in mice subretinally injected as adults.**
C57BL/6J mice (~12-14 weeks old) were injected with 2e10 gc Best1::GFP and harvested 4 weeks later. The neonatal B6J (4e8 gc and 4e9 gc Best1::GFP) RPE and cone counts are replotted here from Figures 3 and 5, respectively. The uninjected B6J (P30) data are replotted here from Figure 1. The uninjected B6J (11 weeks) data was previously published by our lab.[2] Only transduced areas of flatmounts were quantified for adult injected RPE samples. Because transduced areas of retinal flatmounts (as assessed by GFP+ signal) could not be determined for retinal flatmounts from mice injected with Best1::GFP as adults, the adult B6J counts plotted represent quantifications made from boxes drawn roughly equidistantly around each flatmount.
A) Representative images of transduced (left) and untransduced (right) areas of RPE flatmounts from mice subretinally injected with Best1::GFP as adults.
B) RPE toxicity scores from mice subretinally injected with Best1::GFP as adults. Neonatal B6J injections are replotted here for comparison (n=4-16 eyes per group, mean ± SD, ****p<0.0001, **p=0.007, one-way ANOVA with Bonferroni's multiple comparisons correction). Only transduced areas of RPE flatmounts (as assessed by GFP+ signal) were quantified in adult RPE flatmounts.
C) Cone counts for mice subretinally injected with Best1::GFP as adults (n=4-22 eyes per group, mean ± SD, ****p<0.0001, one-way ANOVA with Bonferroni's multiple comparisons correction).

**Figure S17: Effect of CMV::GFP expression in mice subretinally injected as adults.** C57BL/6J mice (16-20 weeks old) were injected with 2e9 gc CMV::GFP and harvested 4 weeks later. The neonatal B6J and IFNAR KO (4e8 gc CMV::GFP) RPE and cone counts are replotted here from Figure S8. The uninjected B6J (P30) data are replotted here from Figure 1. The uninjected B6J (11 weeks) data was previously published by our lab.[2] Only transduced areas of flatmounts were quantified for adult injected samples.

A) RPE scores from RPE flatmounts of mice subretinally injected with CMV::GFP as adults (n=4-6 eyes per group, mean ± SD, *p=0.0158 for adult B6J vs. neonatal B6J, **p=0.005 for adult B6J vs neonatal IFNAR KO, **p=0.001 for adult B6J vs uninjected B6J, *p=0.02, one-way ANOVA with Bonferroni's multiple comparisons correction).

B) Cone counts for mice subretinally injected with CMV::GFP as adults (n=4-8 eyes per group, mean ± SD, ****p<0.0001, one-way ANOVA with Bonferroni's multiple comparisons correction).

**Figure S18: Comparison of injected Ai75d heterozygous animals and C57BL/6J animals used interchangeably for experiments.** The B6J RPE and cone count controls (injected with 4e8 gc Best1::GFP) are replotted here from Figure 3.

A) RPE toxicity scores from RPE flatmounts of C57BL/6J or Ai75d heterozygous pups injected at birth with 4e8 gc Best1::GFP and harvested at 4-5 weeks of age (n=6-16 eyes per group, mean ± SD, unpaired t-test).

B) Cone counts for mice C57BL/6J or Ai75d heterozygous pups injected at birth with 4e8 gc Best1::GFP and harvested at 4-5 weeks of age (n=6-22 eyes per group, mean ± SD, unpaired t-test).

#### Supplemental Table Legends

**Table S1: GO pathways associated with genes upregulated in the Best1::GFP vs**

**Rho::GFP samples at Week 1.** The number and identity of genes significantly upregulated from each GO pathway are also provided.

**Table S2: GO pathways associated with genes upregulated in the Best1::GFP vs Rho::GFP samples at Week 2.** The number and identity of genes significantly upregulated from each GO pathway are also provided.

**Table S3: List of genes significantly upregulated (p-adj < 0.05) in the Best1::GFP vs Rho::GFP samples at Week 1.** List is sorted from lowest to highest p-adj value.

**Table S4: List of genes significantly upregulated (p-adj < 0.05) in the Best1::GFP vs Rho::GFP samples at Week 2.** List is sorted from lowest to highest p-adj value.

**Table S5: List of ISGs significantly upregulated (p-adj < 0.05) in the Best1::GFP vs Rho::GFP samples at Week 1.** Genes were identified as ISGs by submitting the list of genes in Table S3 to the Interferome database.[3]

**Table S6: List of ISGs significantly upregulated (p-adj < 0.05) in the Best1::GFP vs Rho::GFP samples at Week 2.** Genes were identified as ISGs by submitting the list of genes in Table S4 to the Interferome database.[3]

**Table S7: RPE RNA-sequencing gene counts per sample.** Read counts were normalized using DESeq2's median of ratios method.[4] Conditions include Rho::GFP or Best1::GFP RPE samples injected at a 4e8 gc dose. RPE were harvested at week 1 or week 2 post infection.

### Figure S1

A

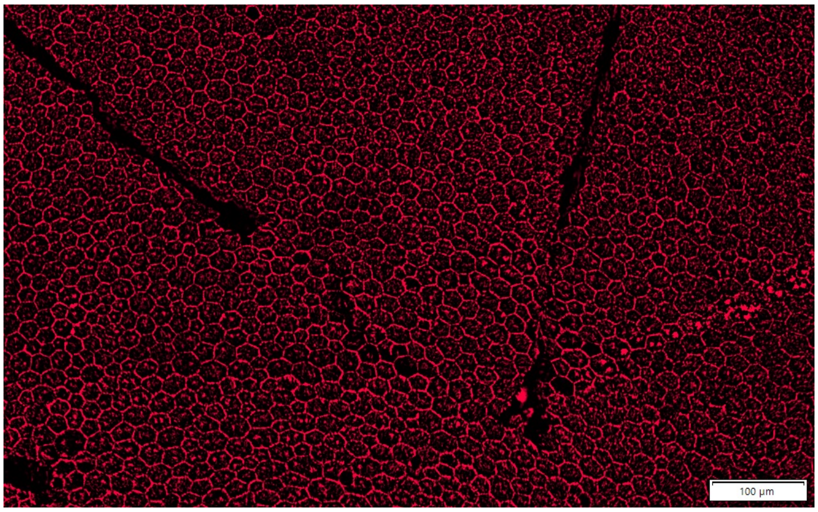


| RPE Toxicity Score | Meaning |
|---|---|
| 0 | Regular hexagonal array of cells. Phalloidin staining is mostly restricted to cell outlines |
| 1 | Imperfect hexagonal array, some cells have disrupted morphology, increased and intracellular phalloidin staining |
| 2 | More disrupted RPE cells, some small RPE cells or rosette-like arrangements, phalloidin staining increased throughout cells |
| 3 | Cell enlargement, in addition to (2) |
| 4 | Even more cell enlargement |
| 5 | Cell dropout |

B

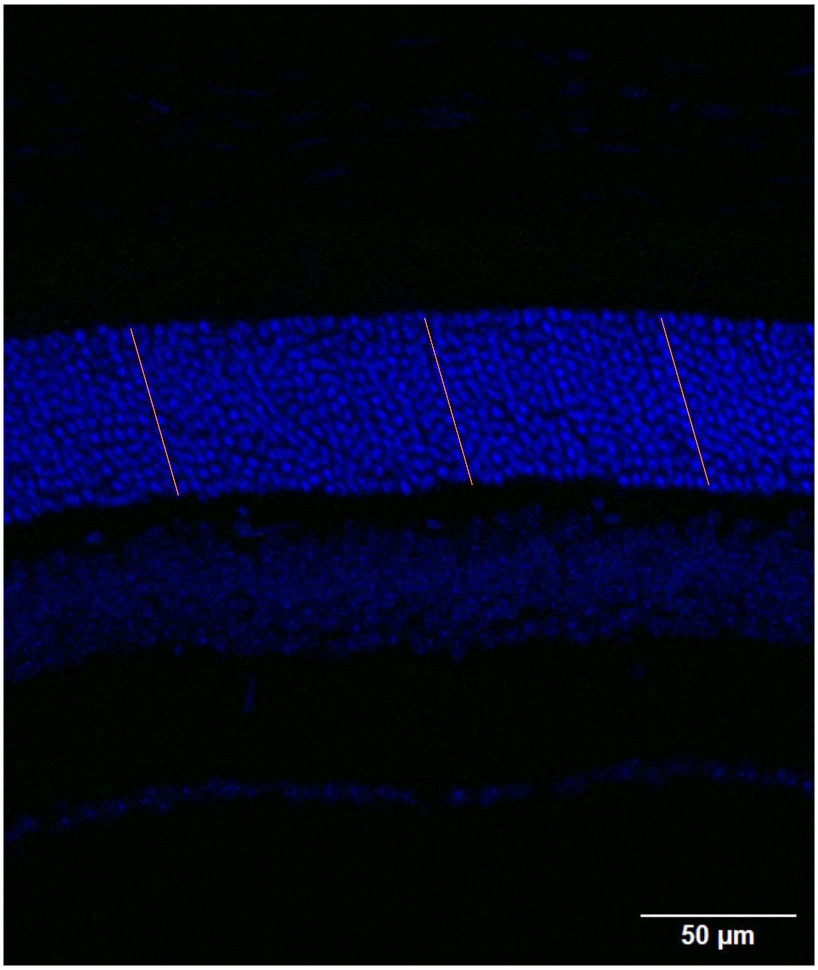

### Figure S2

A

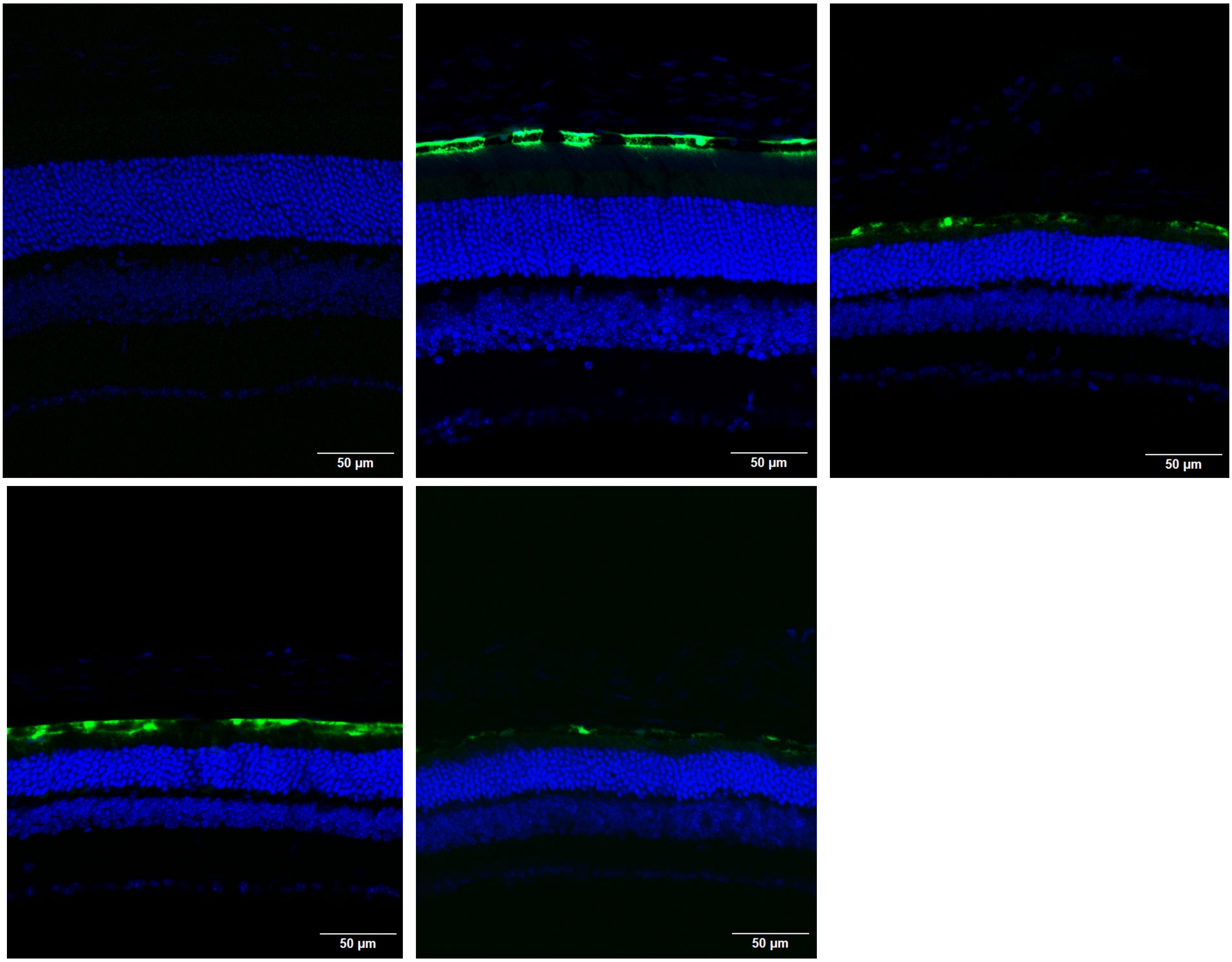
50 µm
50 µm
50 µm
50 µm
50 µm

B

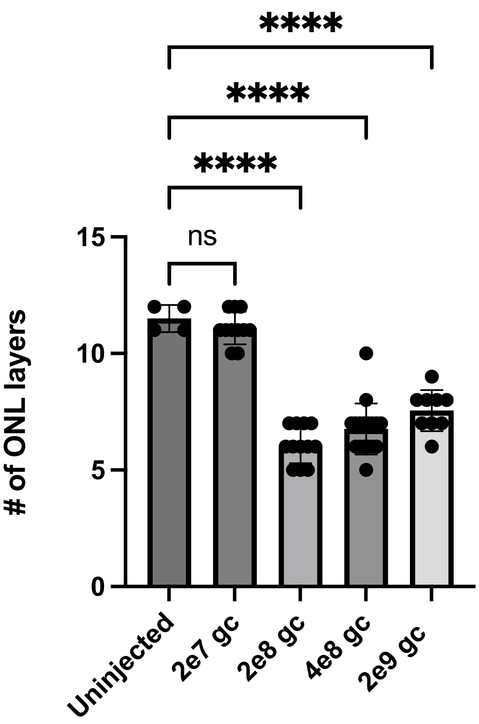
****
****
****
ns
15
10
5
0
### of ONL layers
Uninjected
2e7 gc
2e8 gc
4e8 gc
2e9 gc

### Figure S3

**A**

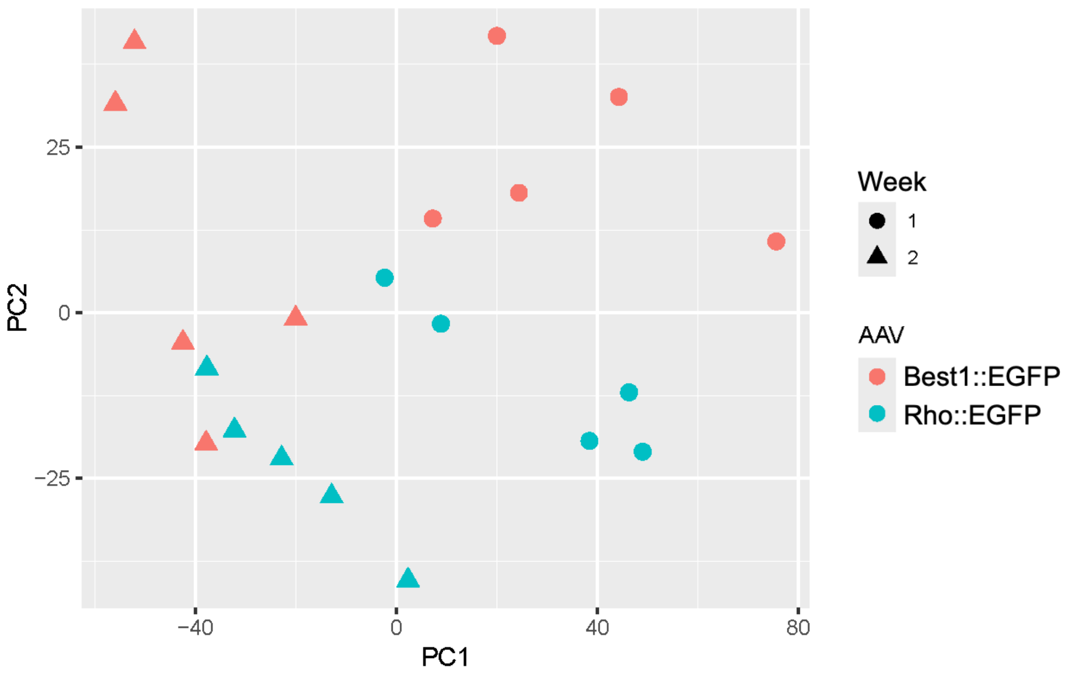


**B**

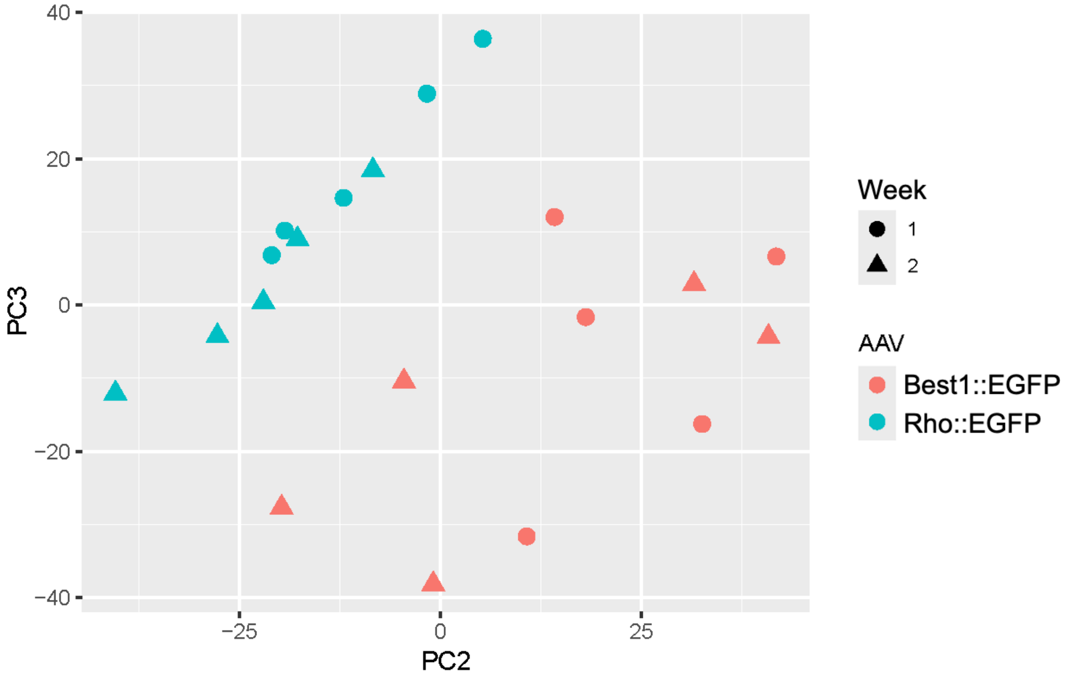


**C**

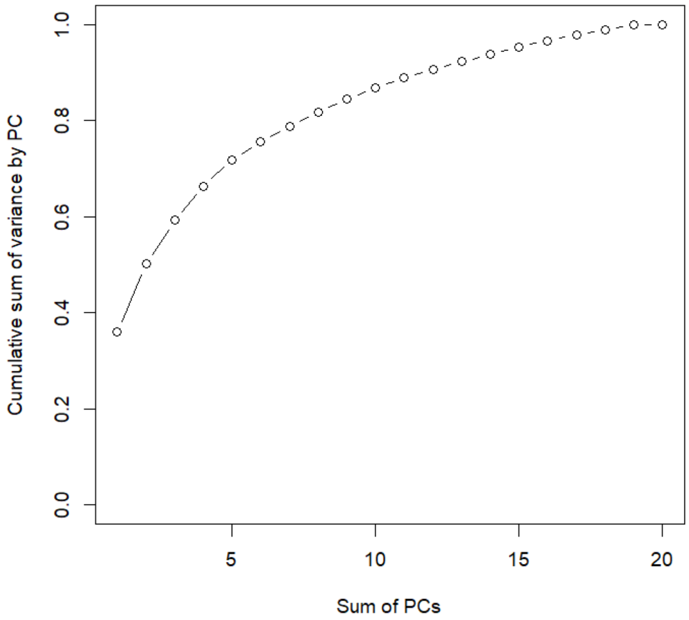

### Figure S4

**A**

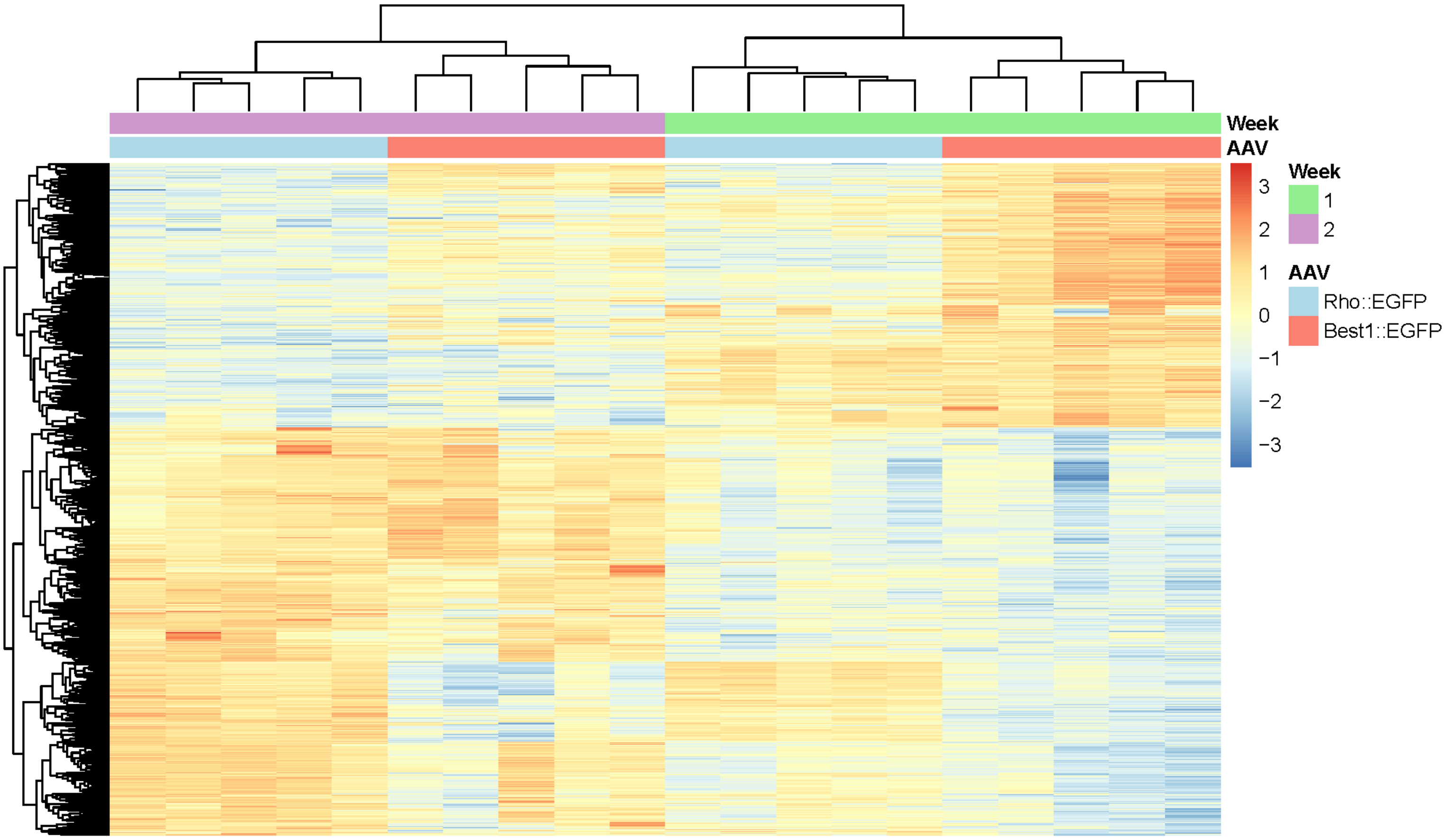

### Figure S5

**A**

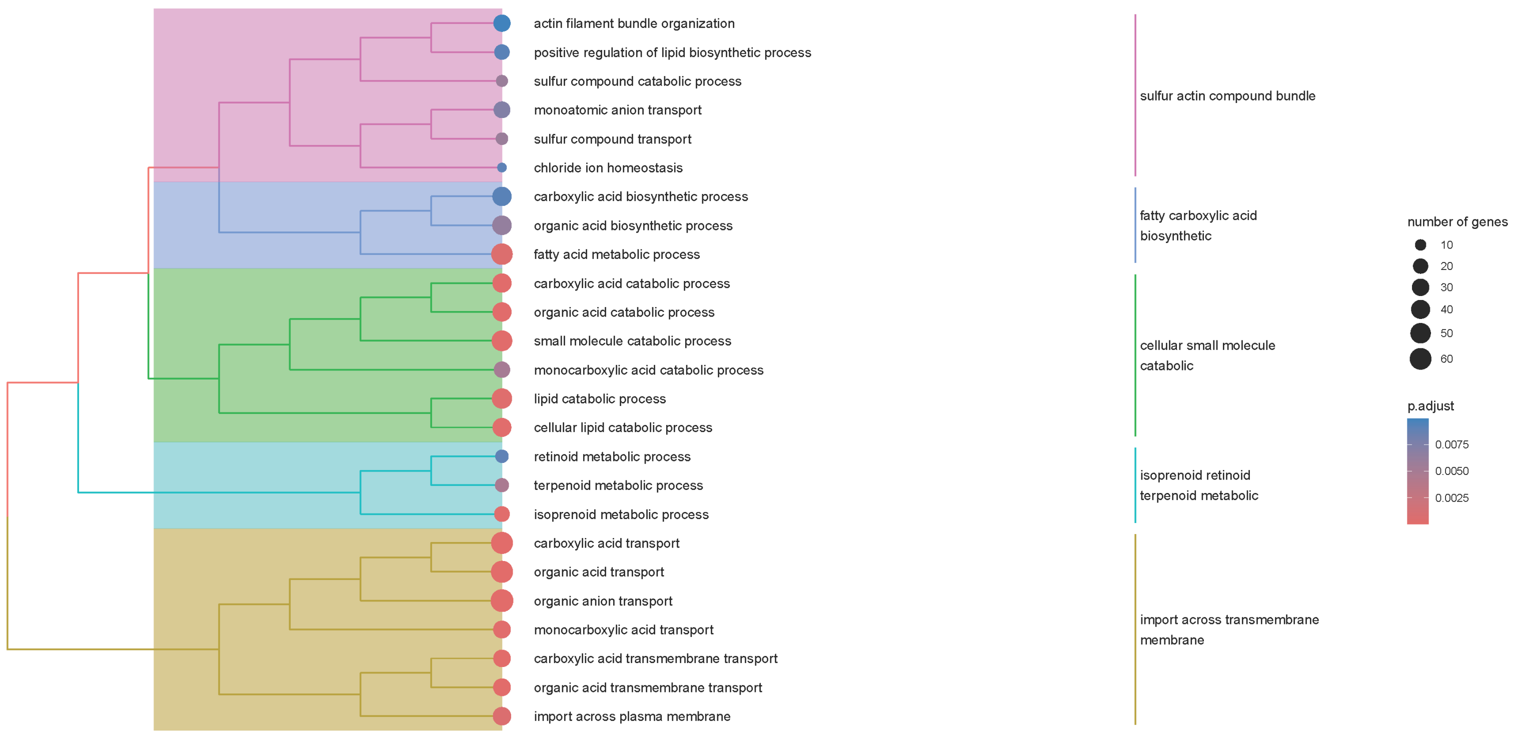

actin filament bundle organization
positive regulation of lipid biosynthetic process
sulfur compound catabolic process
monoatomic anion transport
sulfur compound transport
chloride ion homeostasis
carboxylic acid biosynthetic process
organic acid biosynthetic process
fatty acid metabolic process
carboxylic acid catabolic process
organic acid catabolic process
small molecule catabolic process
monocarboxylic acid catabolic process
lipid catabolic process
cellular lipid catabolic process
retinoid metabolic process
terpenoid metabolic process
isoprenoid metabolic process
carboxylic acid transport
organic acid transport
organic anion transport
monocarboxylic acid transport
carboxylic acid transmembrane transport
organic acid transmembrane transport
import across plasma membrane
sulfur actin compound bundle
fatty carboxylic acid biosynthetic
cellular small molecule catabolic
isoprenoid retinoid terpenoid metabolic
import across transmembrane membrane
number of genes
10
20
30
40
50
60
p.adjust
0.0075
0.0050
0.0025


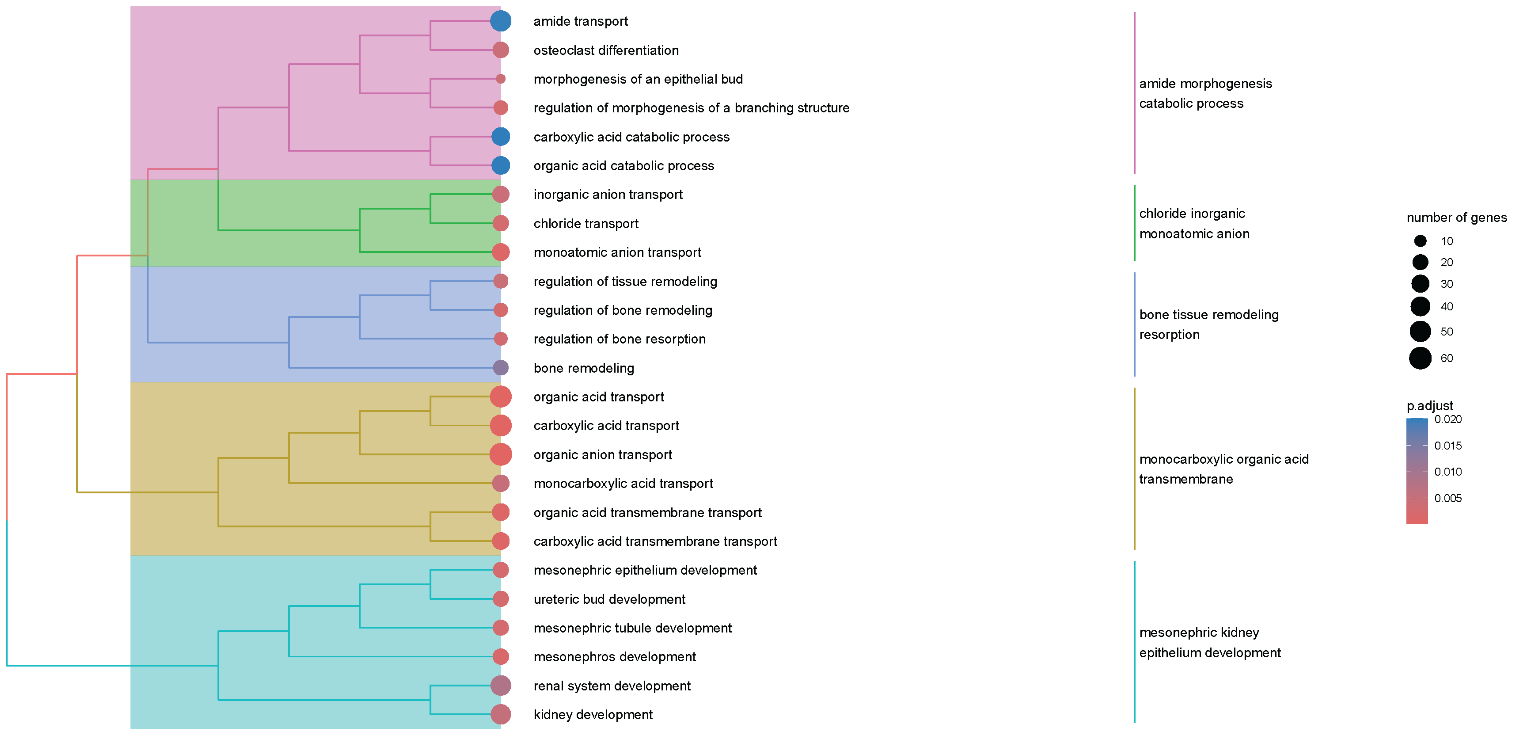

amide transport
osteoclast differentiation
morphogenesis of an epithelial bud
regulation of morphogenesis of a branching structure
carboxylic acid catabolic process
organic acid catabolic process
inorganic anion transport
chloride transport
monoatomic anion transport
regulation of tissue remodeling
regulation of bone remodeling
regulation of bone resorption
bone remodeling
organic acid transport
carboxylic acid transport
organic anion transport
monocarboxylic acid transport
organic acid transmembrane transport
carboxylic acid transmembrane transport
mesonephric epithelium development
ureteric bud development
mesonephric tubule development
mesonephros development
renal system development
kidney development
amide morphogenesis catabolic process
chloride inorganic monoatomic anion
bone tissue remodeling resorption
monocarboxylic organic acid transmembrane
mesonephric kidney epithelium development
number of genes
10
20
30
40
50
60
p.adjust
0.020
0.015
0.010
0.005

### Figure S6

A

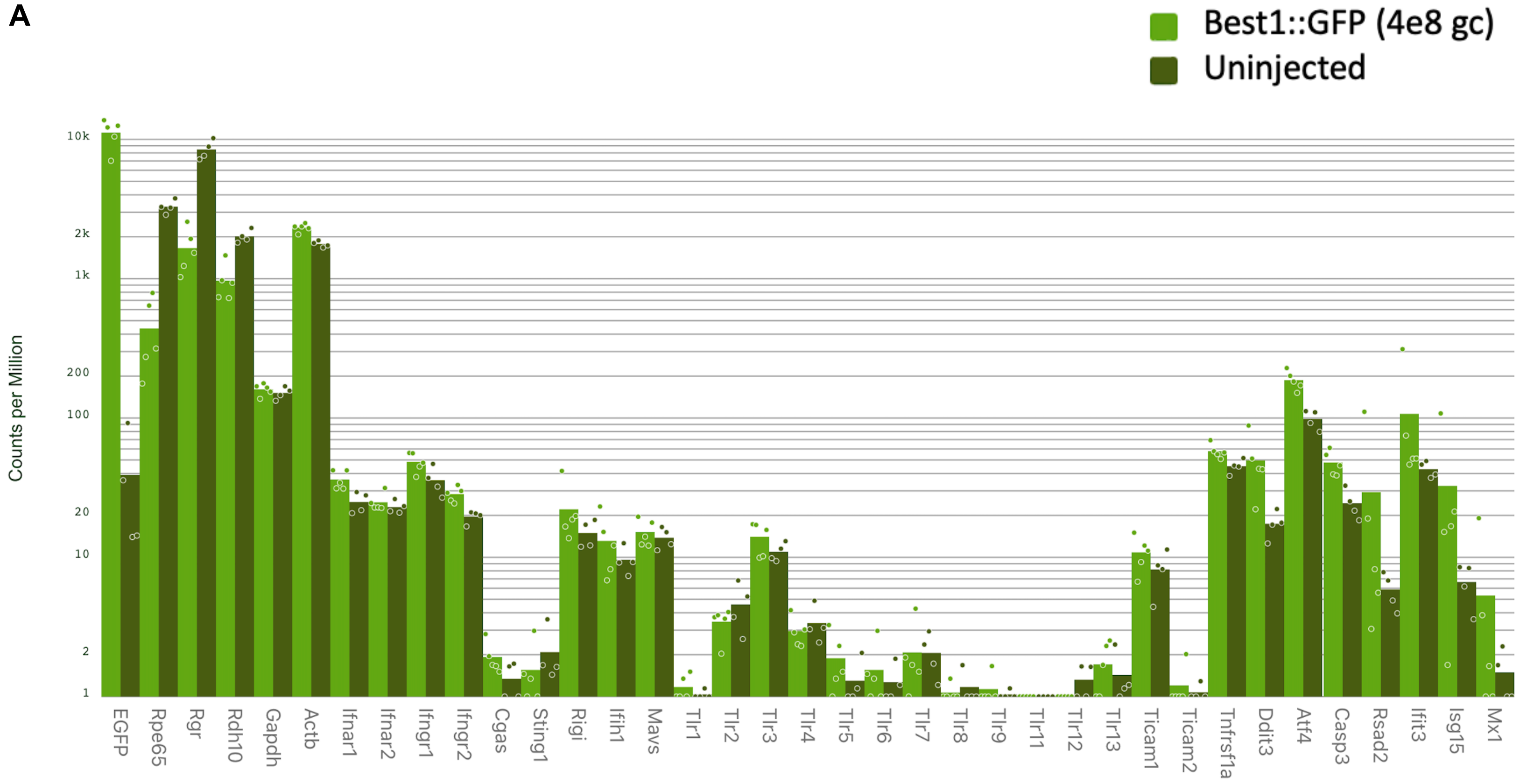
Best1::GFP (4e8 gc)
Uninjected
Counts per Million
10k
2k
1k
200
100
20
10
2
1
EGFP
Rpe65
Rgr
Rdh10
Gapdh
Actb
Ifnar1
Ifnar2
Ifngr1
Ifngr2
Cgas
Sting1
Rigi
Ifih1
Mavs
Tlr1
Tlr2
Tlr3
Tlr4
Tlr5
Tlr6
Tlr7
Tlr8
Tlr9
Tlr11
Tlr12
Tlr13
Ticam1
Ticam2
Tnfrsf1a
Ddit3
Atf4
Casp3
Rsad2
Ifit3
Isg15
Mx1

B

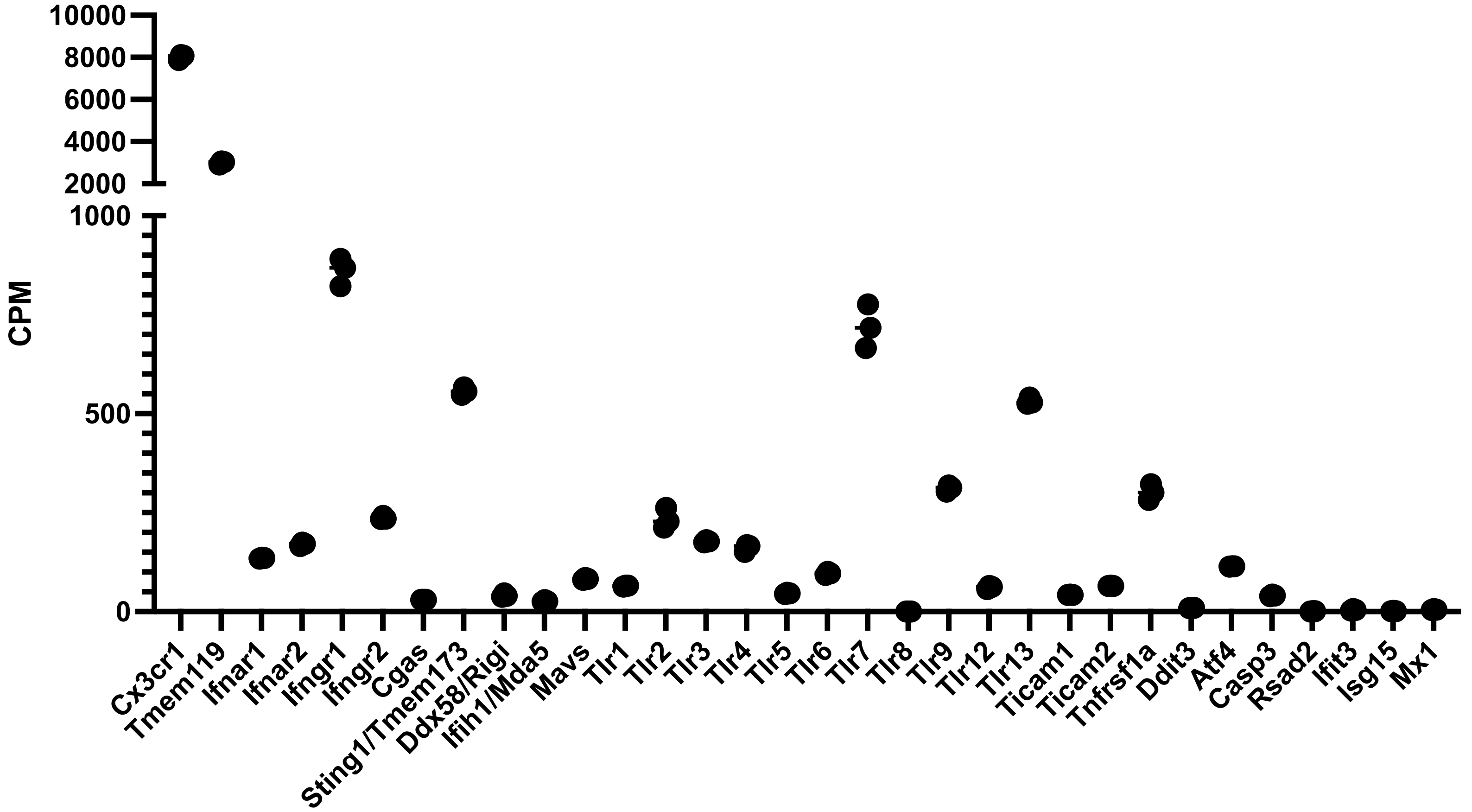
CPM
10000
8000
6000
4000
2000
1000
500
0
Cx3cr1
Tmem119
Ifnar1
Ifnar2
Ifngr1
Ifngr2
Cgas
Sting1/Tmem173
Ddx58/Rigi
Ifih1/Mda5
Mavs
Tlr1
Tlr2
Tlr3
Tlr4
Tlr5
Tlr6
Tlr7
Tlr8
Tlr9
Tlr12
Tlr13
Ticam1
Ticam2
Tnfrsf1a
Ddit3
Atf4
Casp3
Rsad2
Ifit3
Isg15
Mx1

### Figure S7

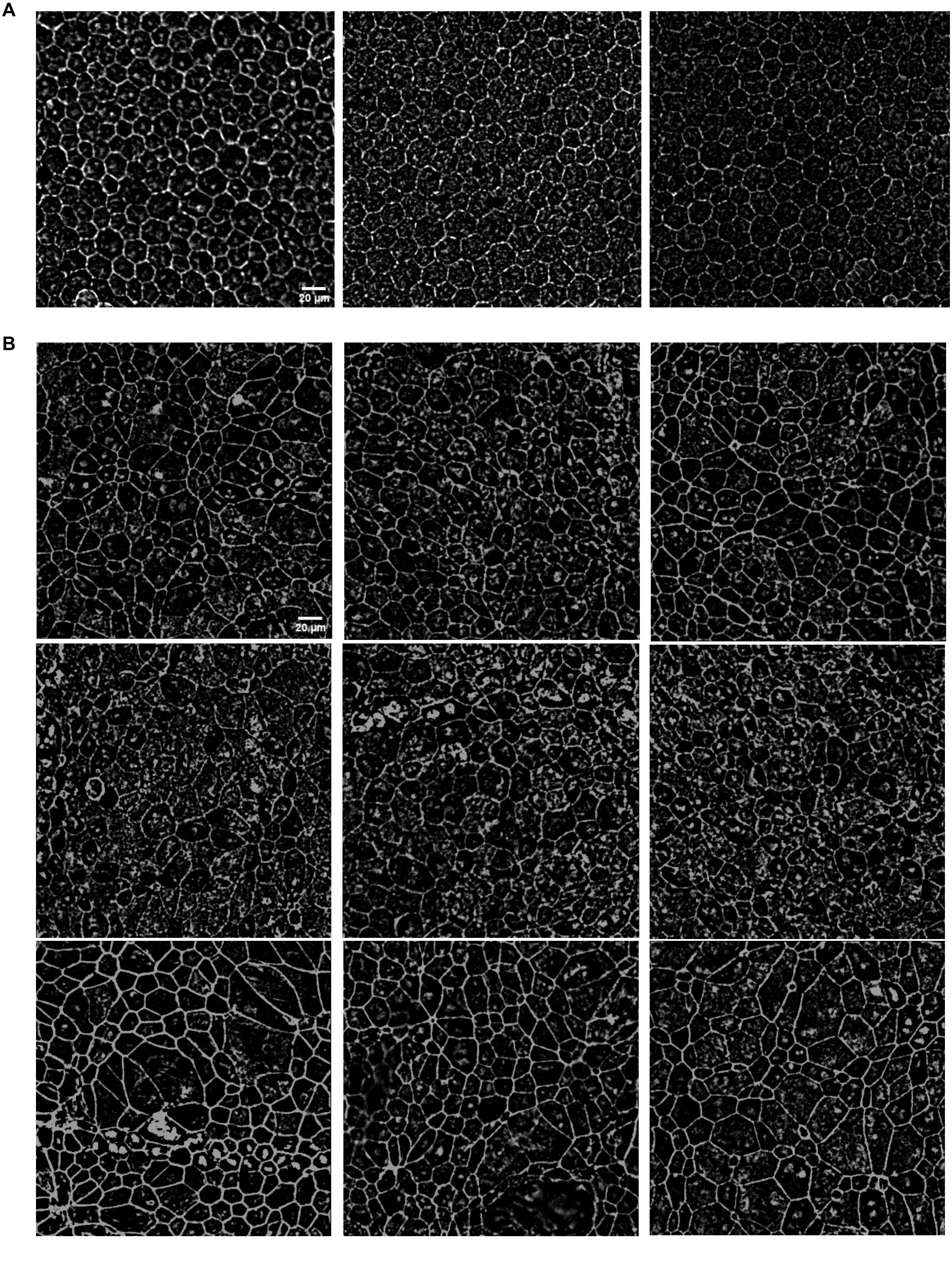

### Figure S7

c

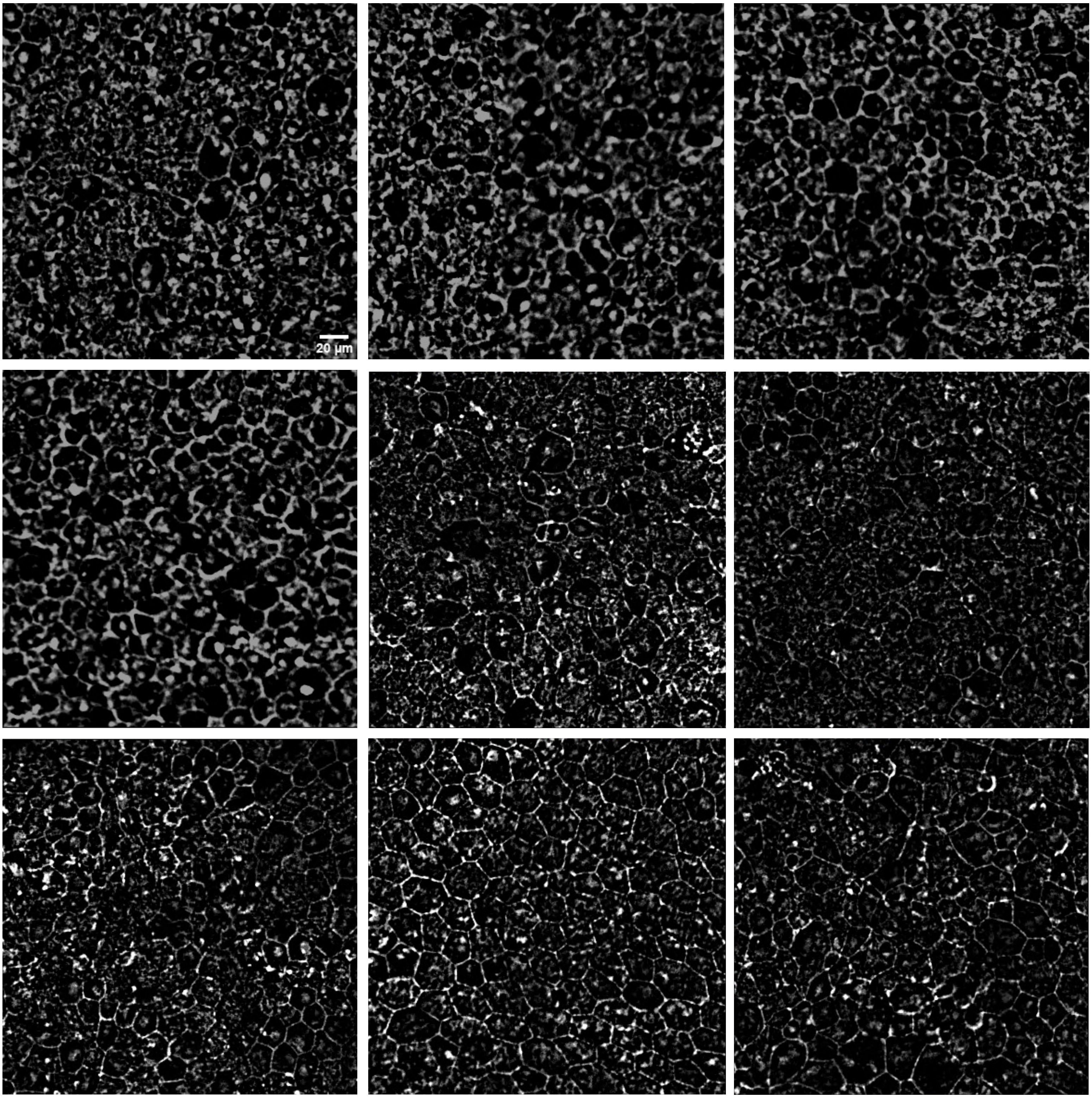

### Figure S7

D

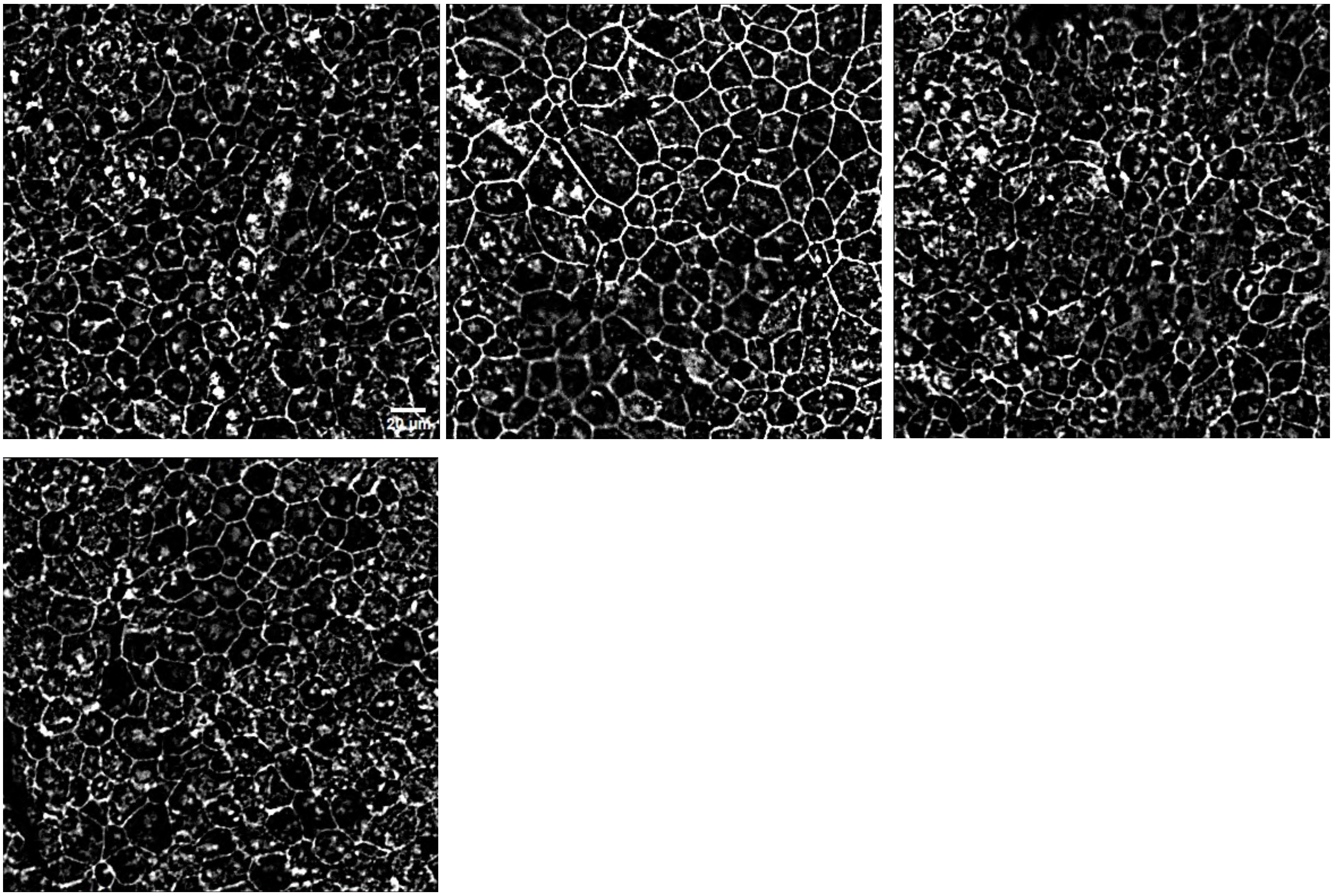

### Figure S7

E

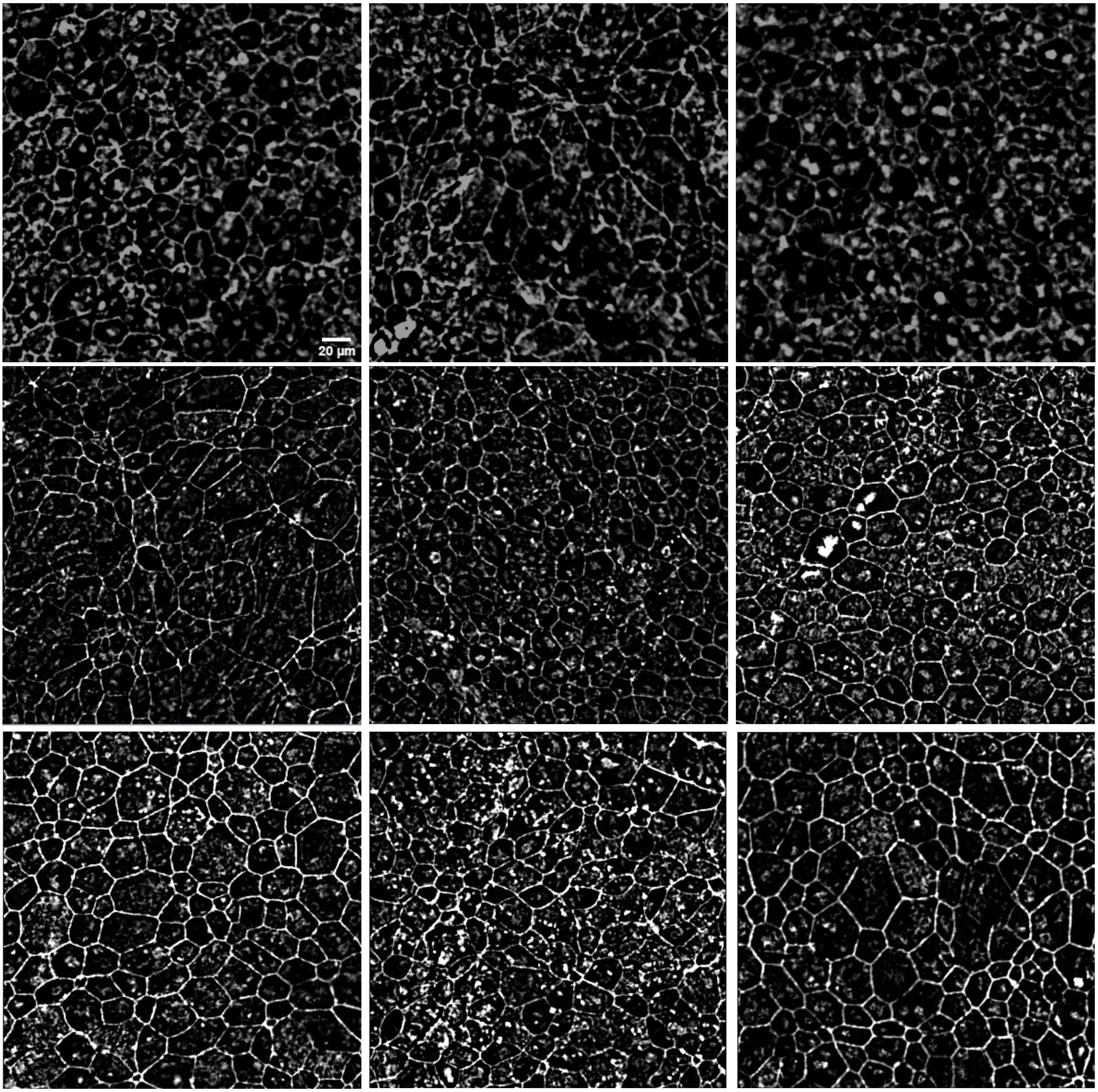

### Figure S7

**F**

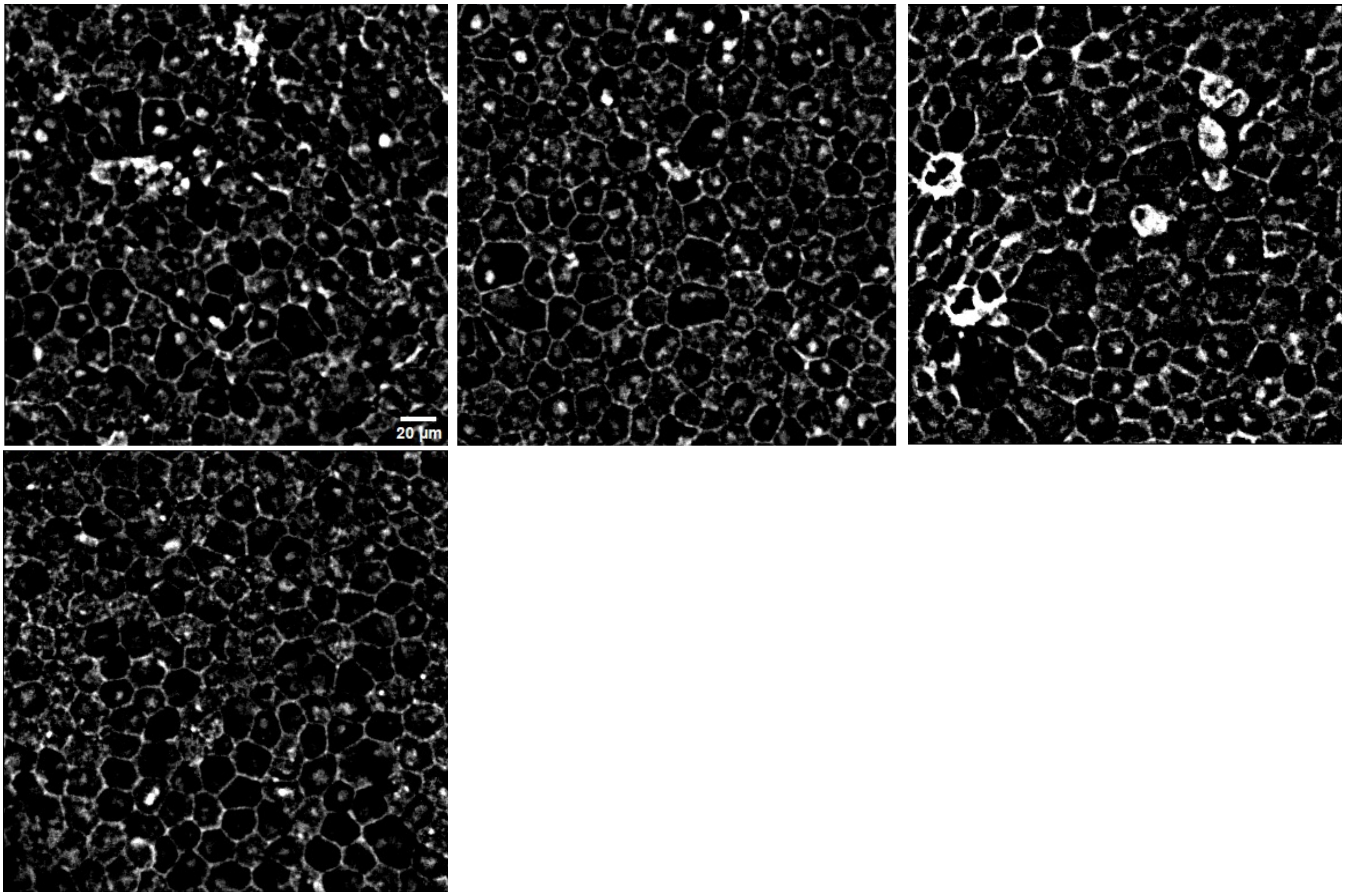

### Figure S7

G

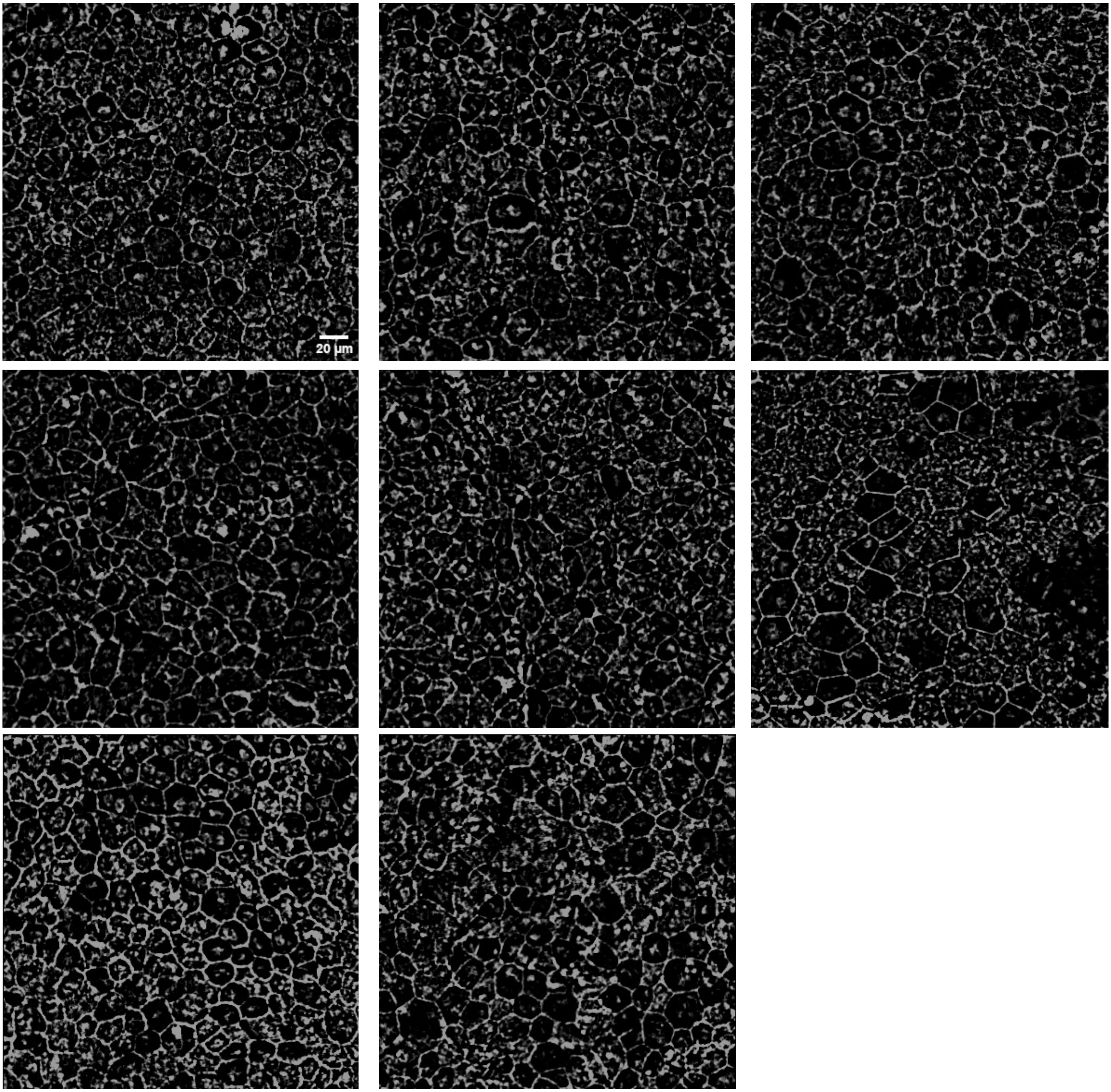

### Figure S7

H

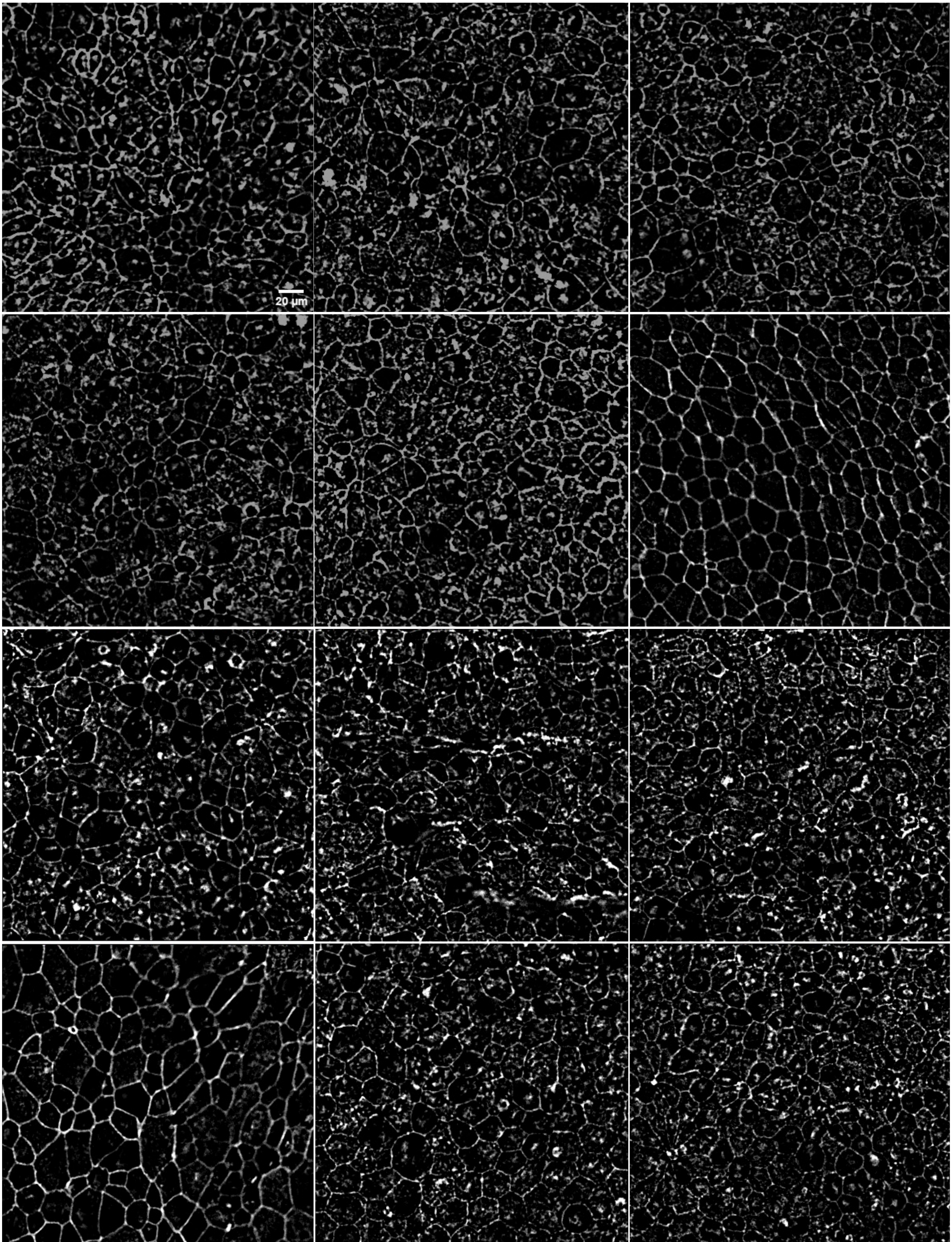

### Figure S7

I

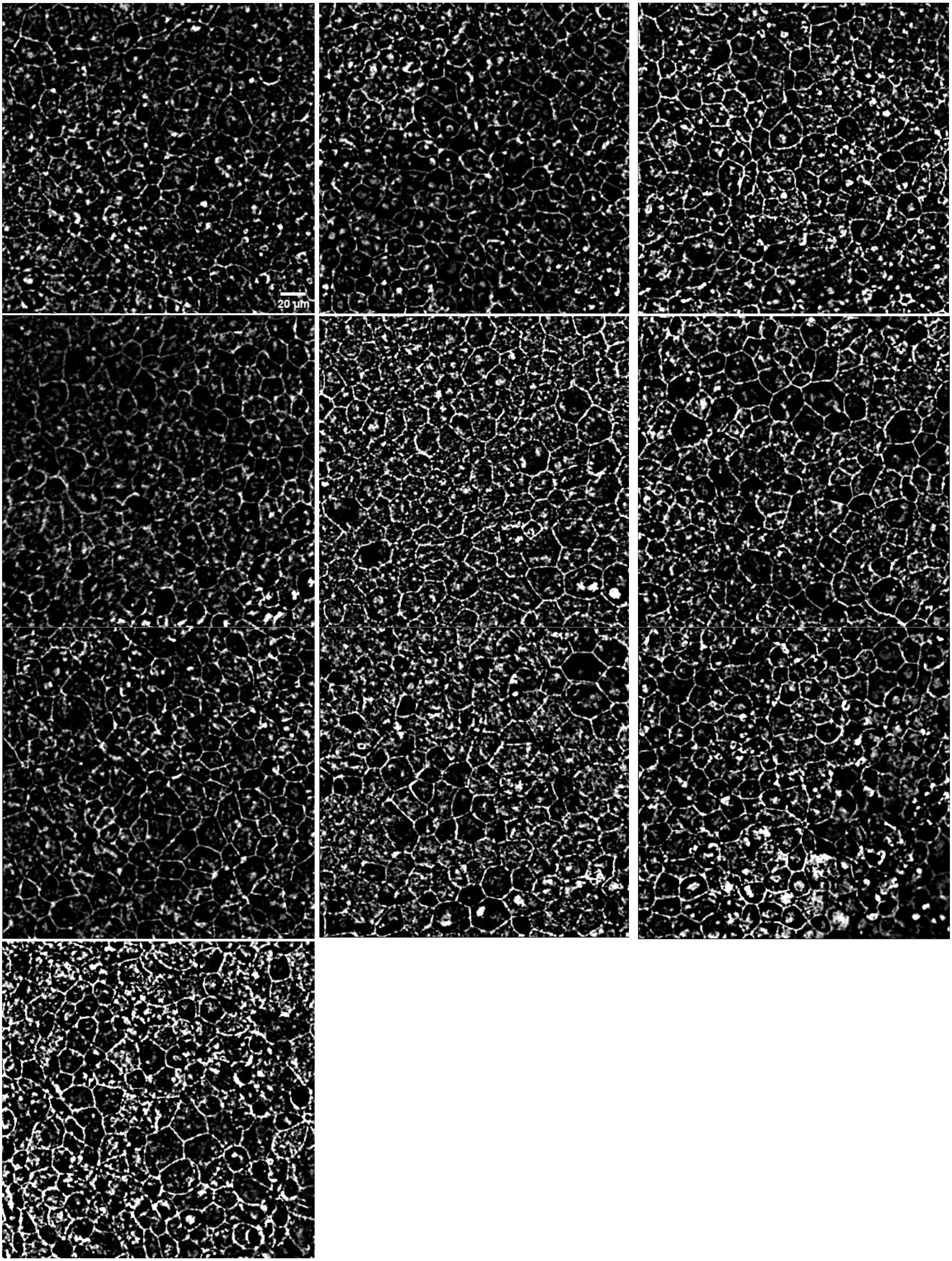

### Figure S7

J

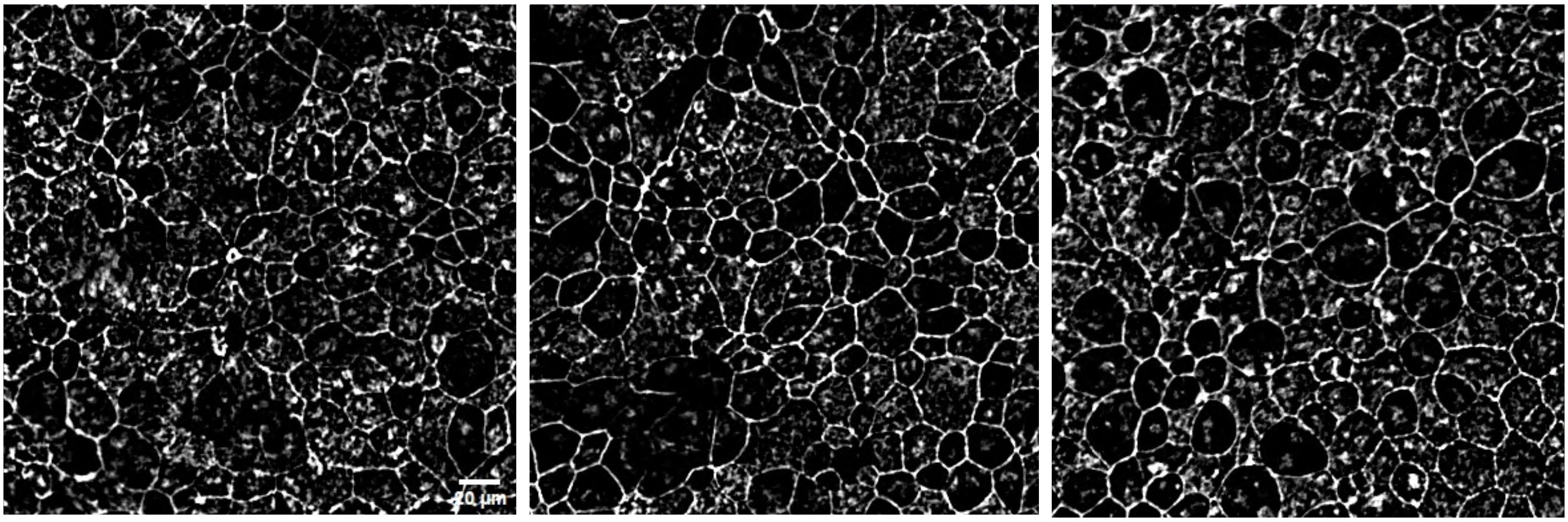


K

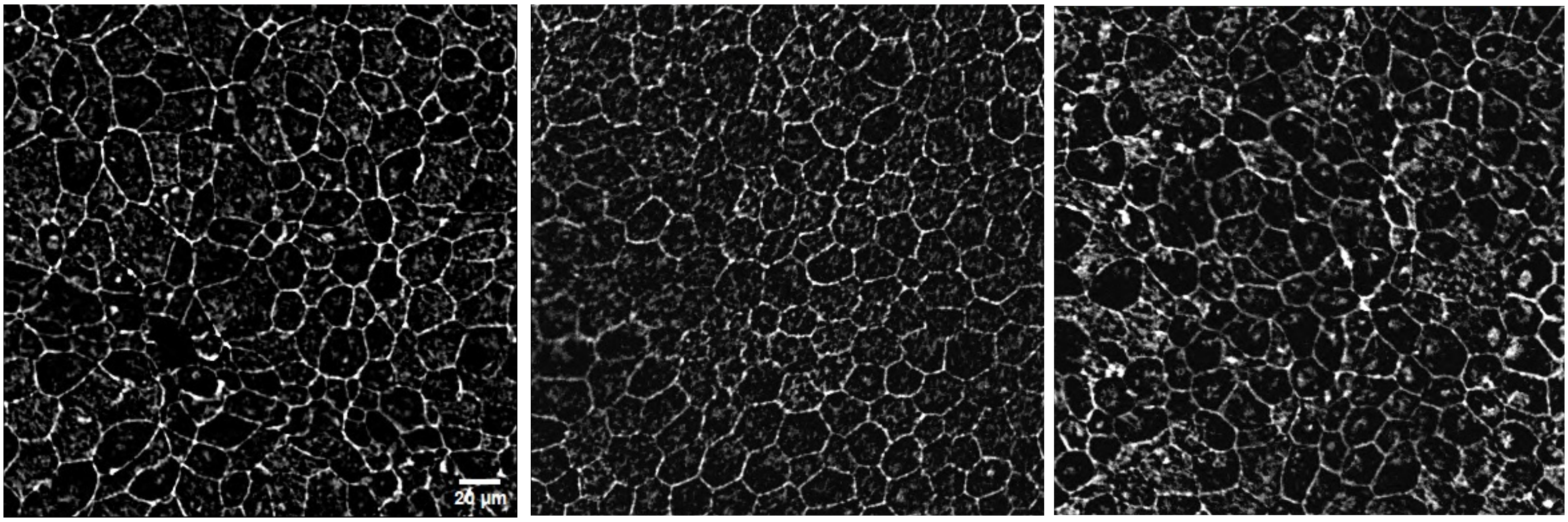

### Figure S7

L

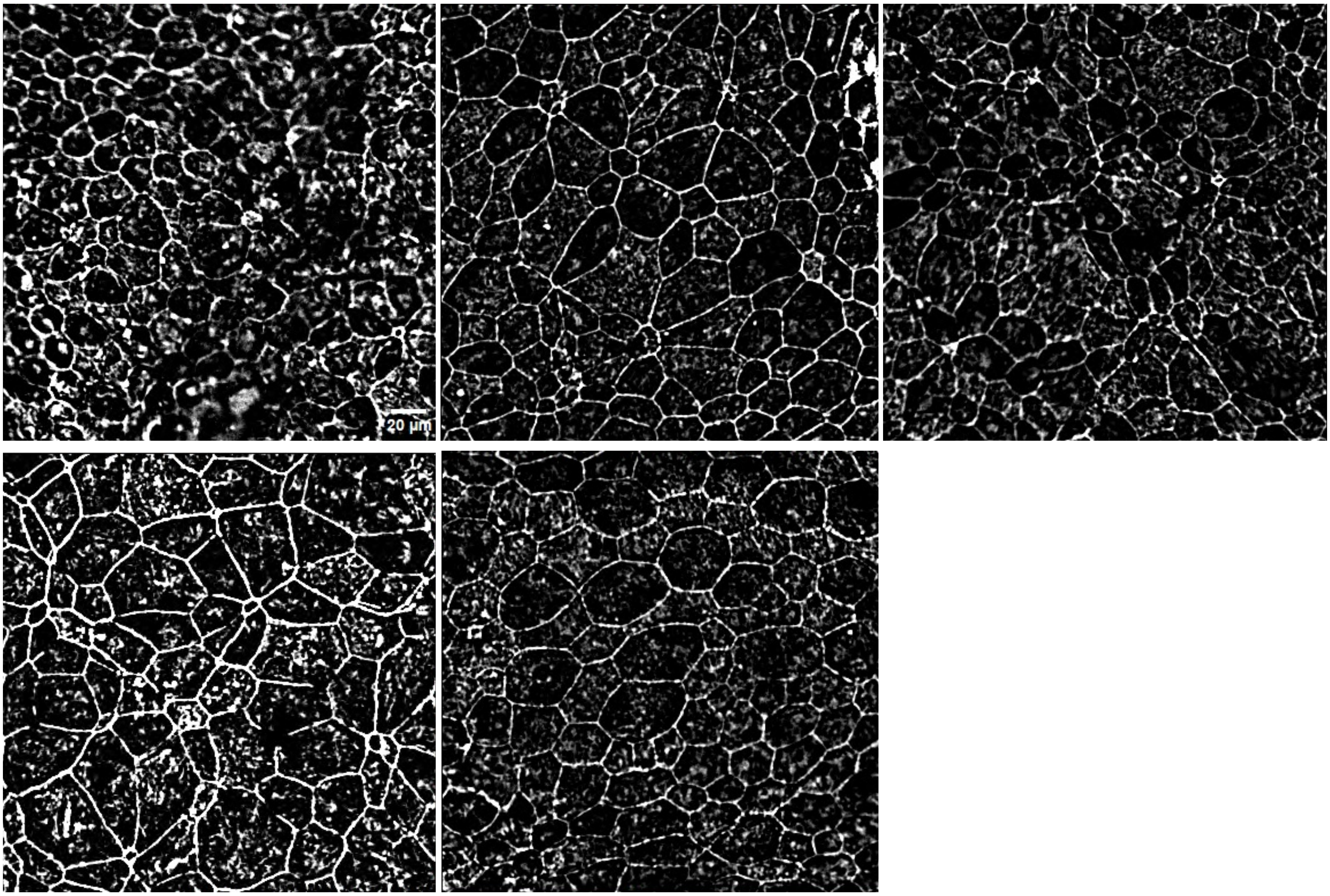

### Figure S8

A

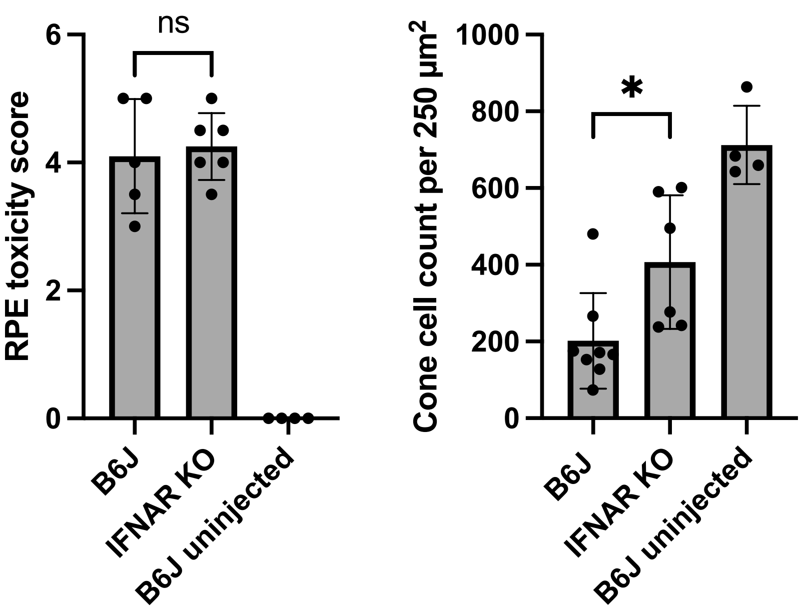

RPE toxicity score
6
4
2
0
ns
B6J
IFNAR KO
B6J uninjected
Cone cell count per 250 µm²
1000
800
600
400
200
0
*
B6J
IFNAR KO
B6J uninjected

### Figure S9

**A**

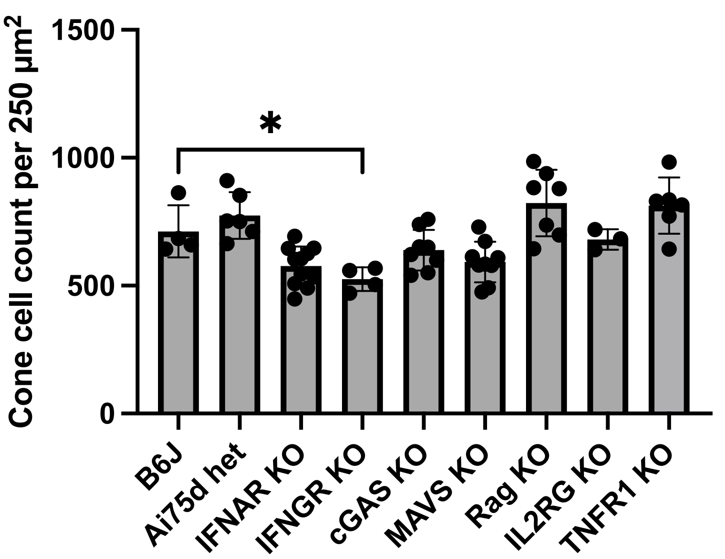

Cone cell count per 250 µm²
1500
1000
500
0
*
B6J
Ai75d het
IFNAR KO
IFNGR KO
cGAS KO
MAVS KO
Rag KO
IL2RG KO
TNFR1 KO


**B**

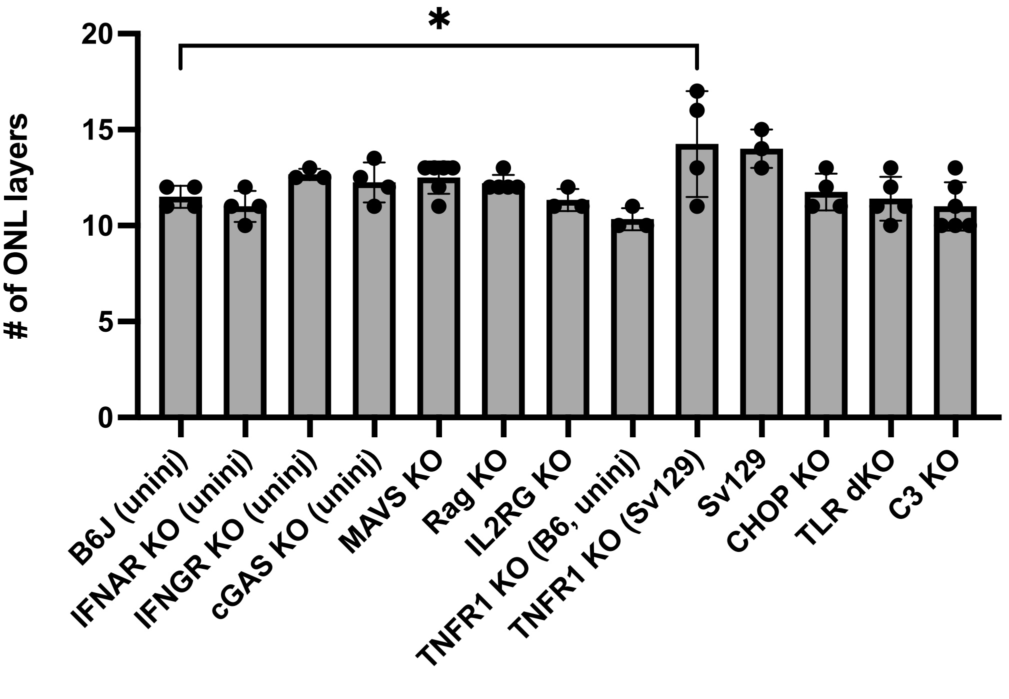

### of ONL layers
20
15
10
5
0
*
B6J (uninj)
IFNAR KO (uninj)
IFNGR KO (uninj)
cGAS KO (uninj)
MAVS KO
Rag KO
IL2RG KO
TNFR1 KO (B6, uninj)
TNFR1 KO (Sv129)
Sv129
CHOP KO
TLR dKO
C3 KO

### Figure S10

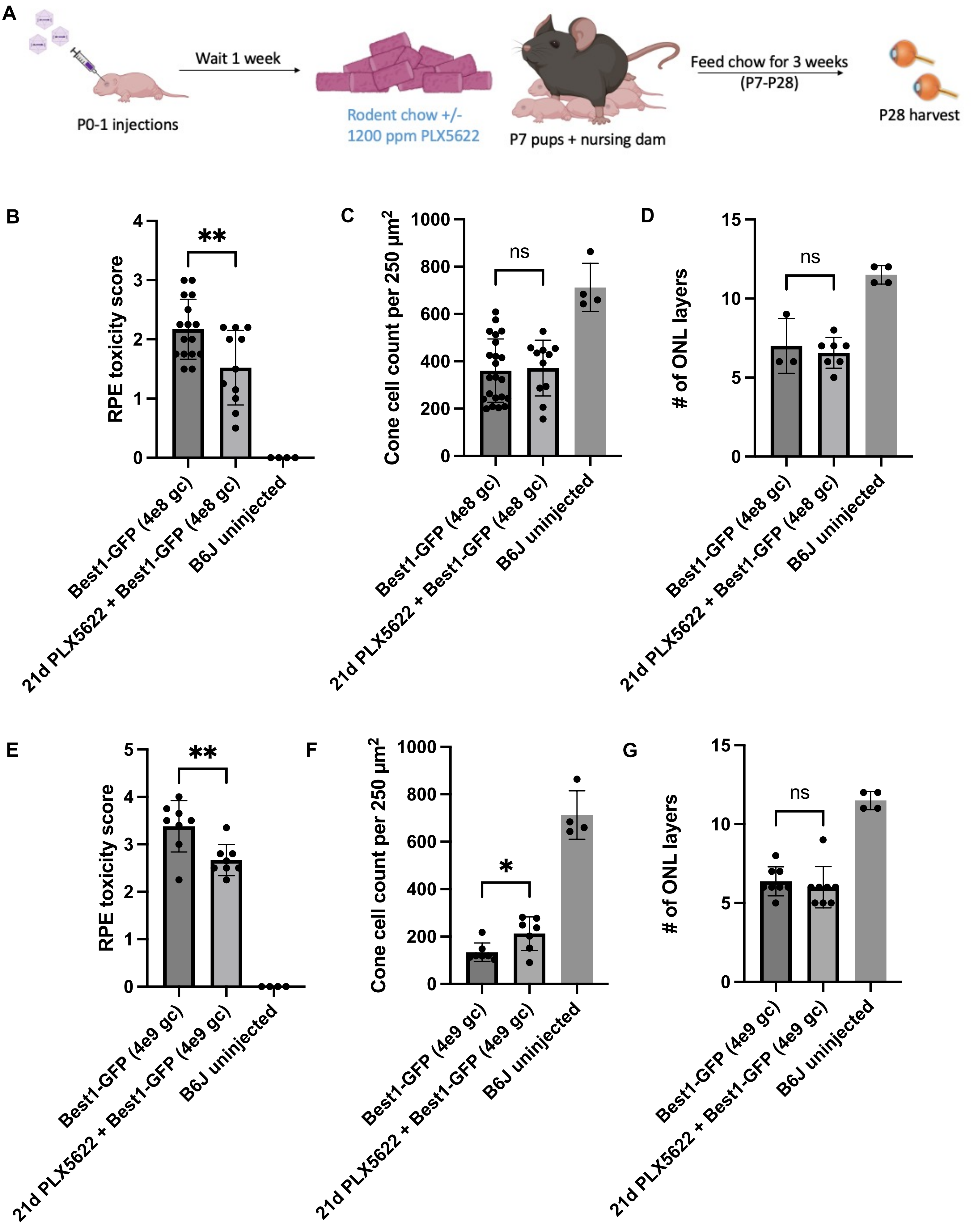

A
Wait 1 week
P0-1 injections
Rodent chow +/-
1200 ppm PLX5622
P7 pups + nursing dam
Feed chow for 3 weeks
(P7-P28)
P28 harvest
B
RPE toxicity score
**
Best1-GFP (4e8 gc)
21d PLX5622 + Best1-GFP (4e8 gc)
B6J uninjected
C
Cone cell count per 250 μm²
ns
D
### of ONL layers
ns
E
RPE toxicity score
**
Best1-GFP (4e9 gc)
21d PLX5622 + Best1-GFP (4e9 gc)
B6J uninjected
F
Cone cell count per 250 μm²
*
G
### of ONL layers
ns

### Figure S11

**A** Regular chow, 4e8 gc Best1::GFP injections at P0

**B** 21d PLX5622 chow, 4e8 gc Best1::GFP injections at P0

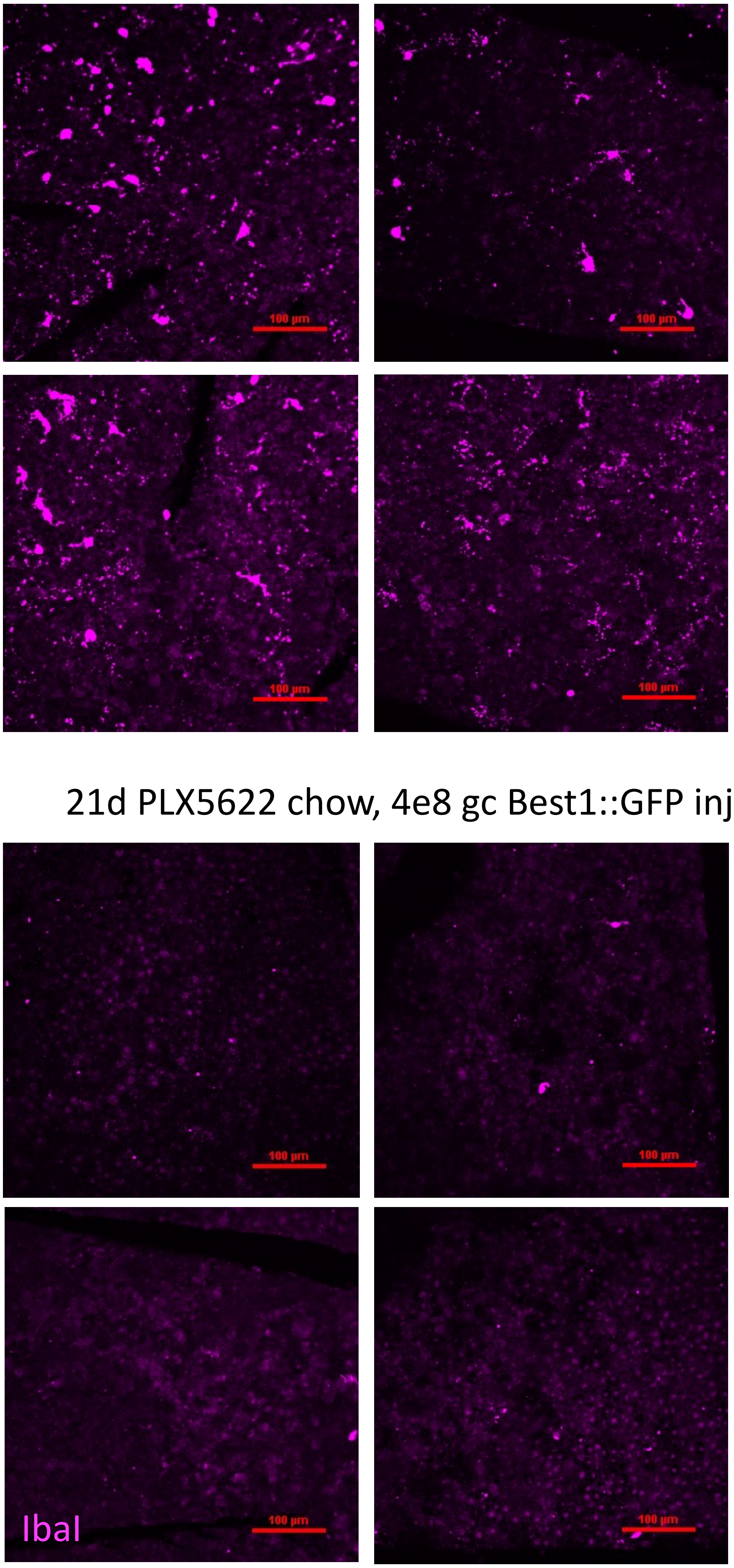


**C**

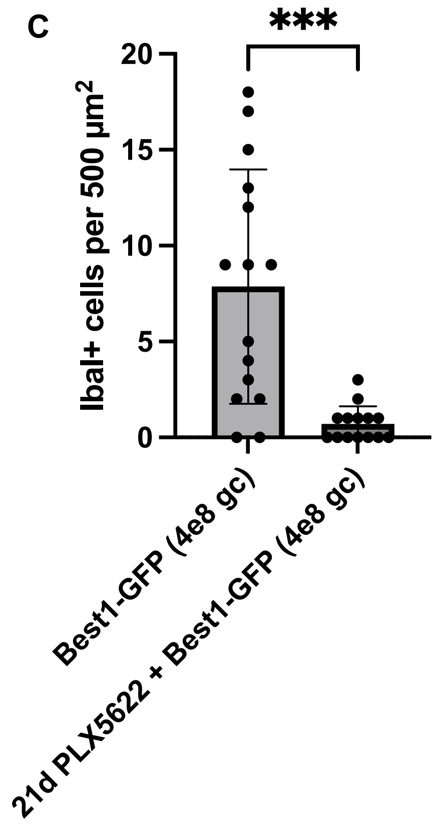

### Figure S12

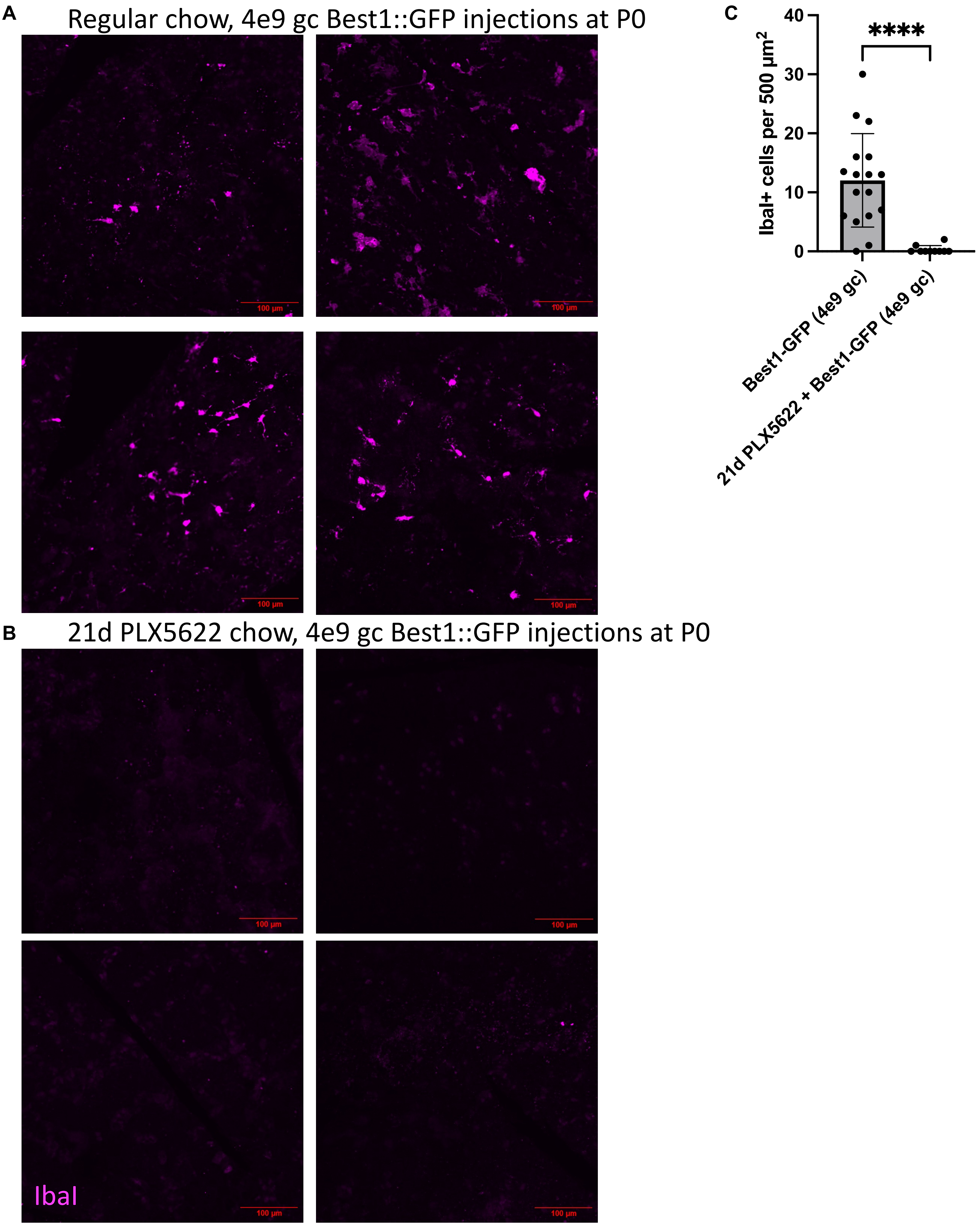
A
Regular chow, 4e9 gc Best1::GFP injections at P0
100 µm
B
21d PLX5622 chow, 4e9 gc Best1::GFP injections at P0
IbaI
C
IbaI+ cells per 500 µm²
40
30
20
10
0
****
Best1-GFP (4e9 gc)
21d PLX5622 + Best1-GFP (4e9 gc)

### Figure S13

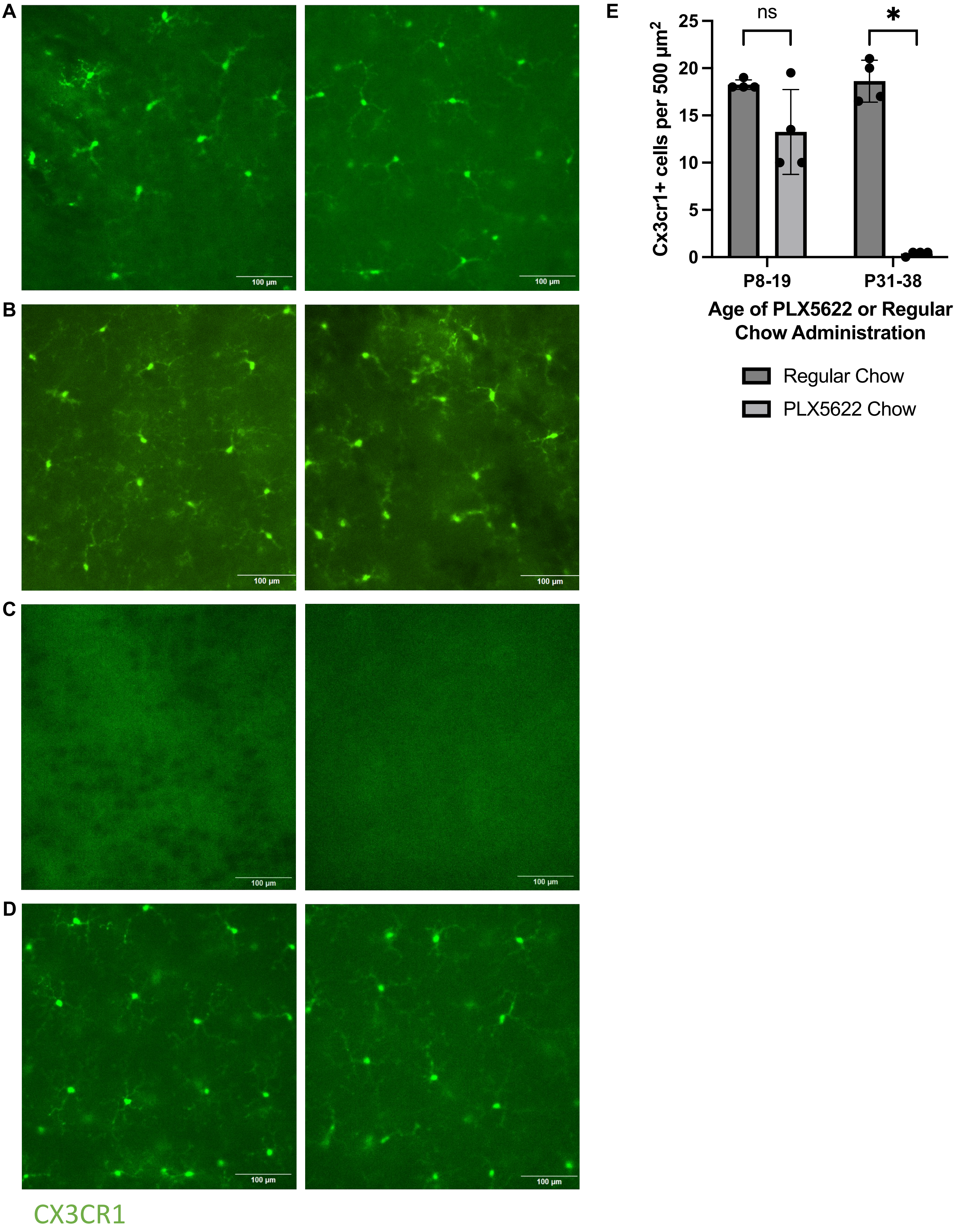
A
B
C
D
E
100 µm
ns
*
Cx3cr1+ cells per 500 µm2
25
20
15
10
5
0
P8-19
P31-38
Age of PLX5622 or Regular Chow Administration
Regular Chow
PLX5622 Chow
CX3CR1

### Figure S14

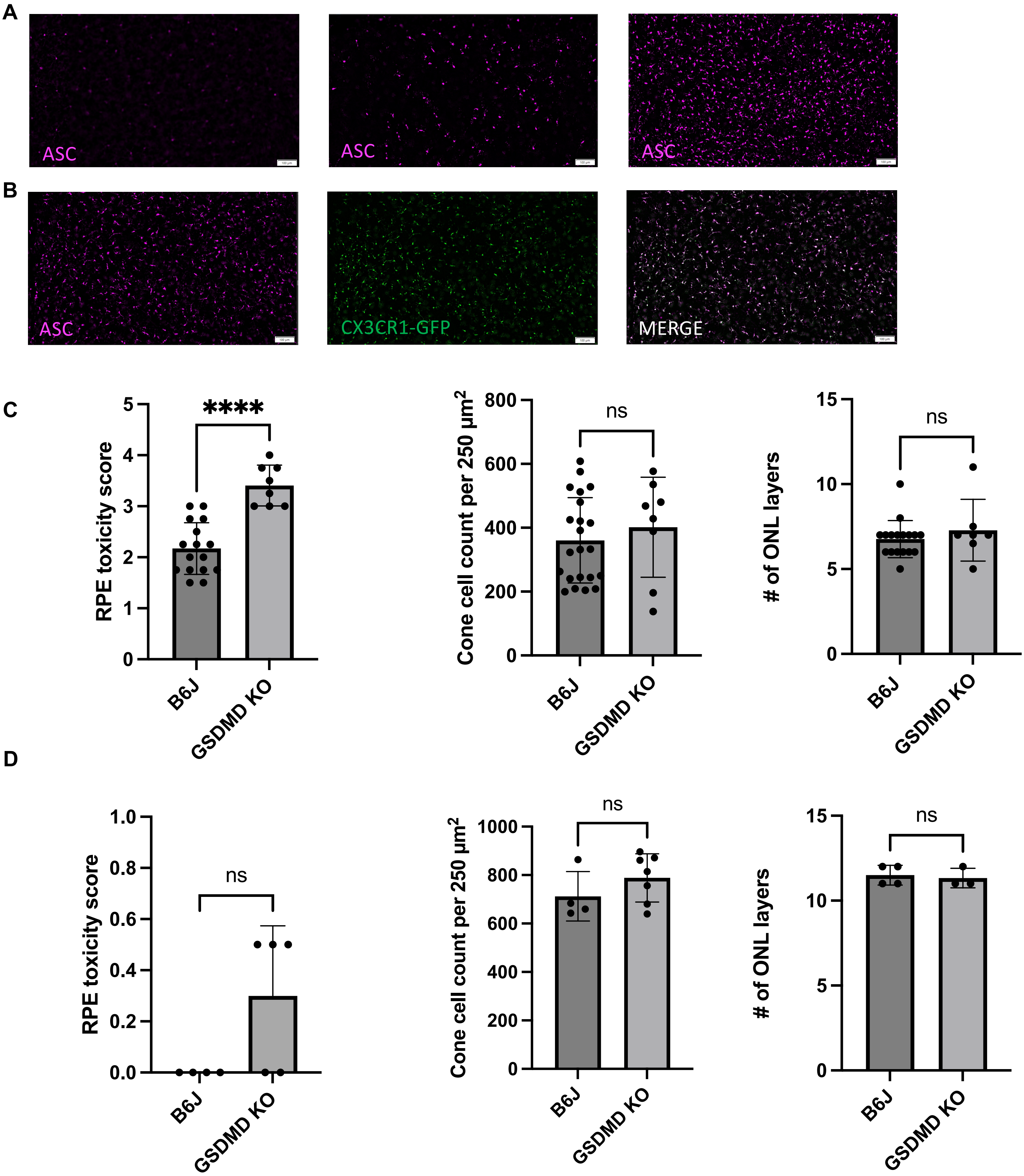
A
ASC
ASC
ASC
B
ASC
CX3CR1-GFP
MERGE
C
RPE toxicity score
****
B6J
GSDMD KO
Cone cell count per 250 µm²
ns
### of ONL layers
ns
D
RPE toxicity score
ns
Cone cell count per 250 µm²
ns
### of ONL layers
ns

### Figure S15

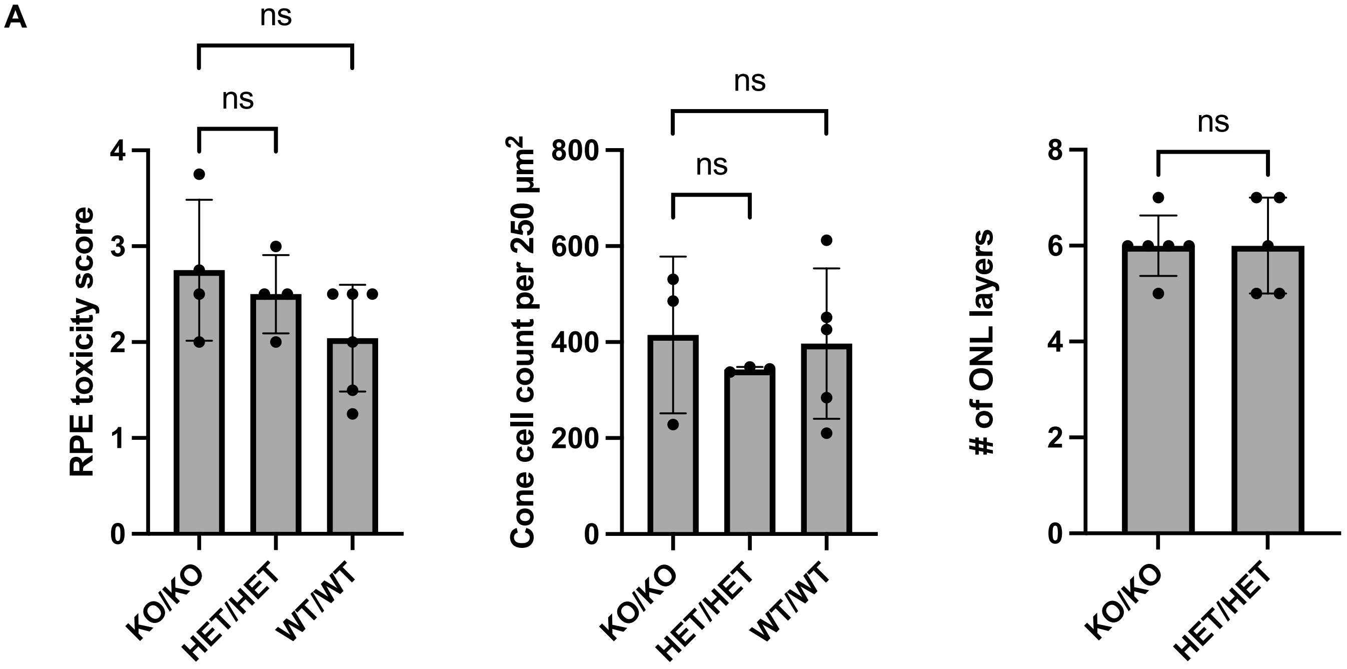
A
ns
ns
RPE toxicity score
4
3
2
1
0
KO/KO
HET/HET
WT/WT
ns
ns
Cone cell count per 250 µm2
800
600
400
200
0
KO/KO
HET/HET
WT/WT
ns
### of ONL layers
8
6
4
2
0
KO/KO
HET/HET

### Figure S16

A

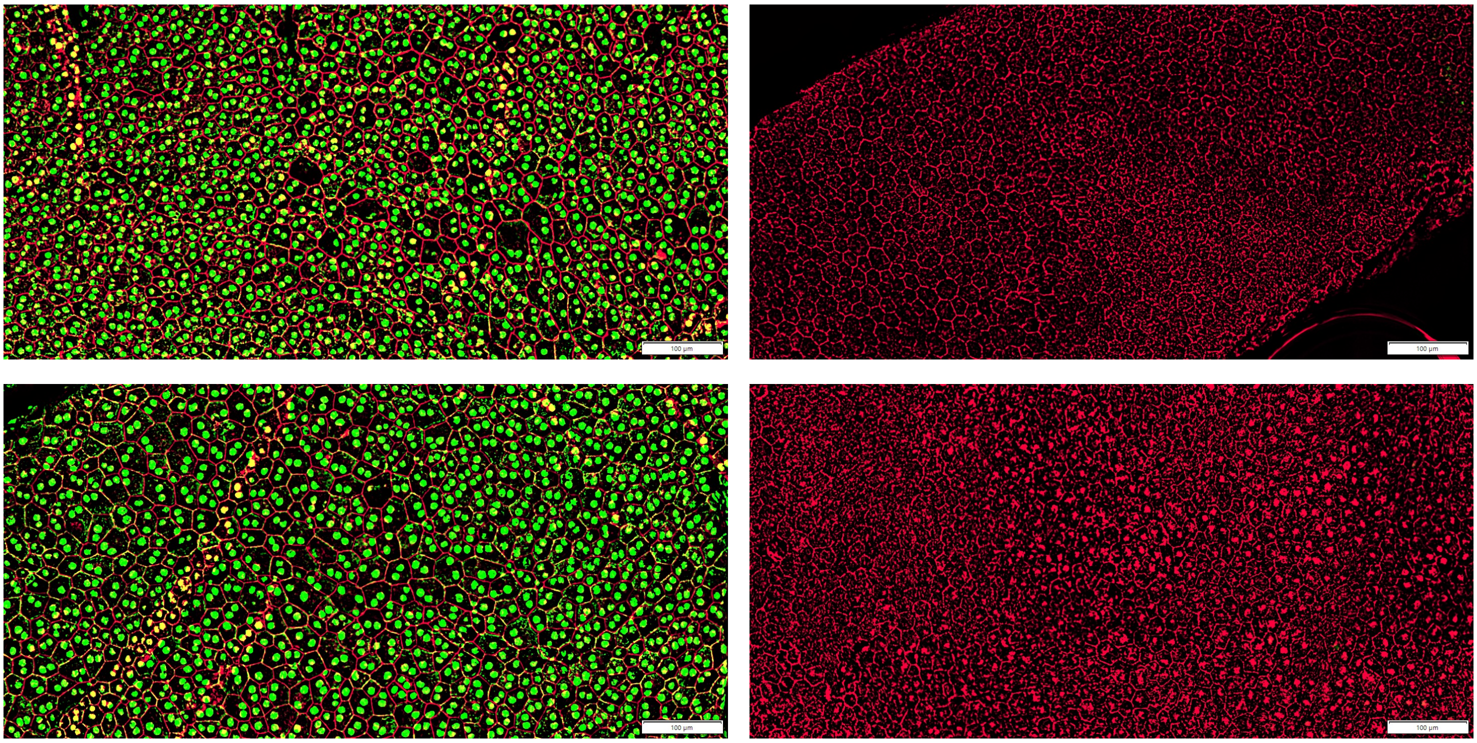


B

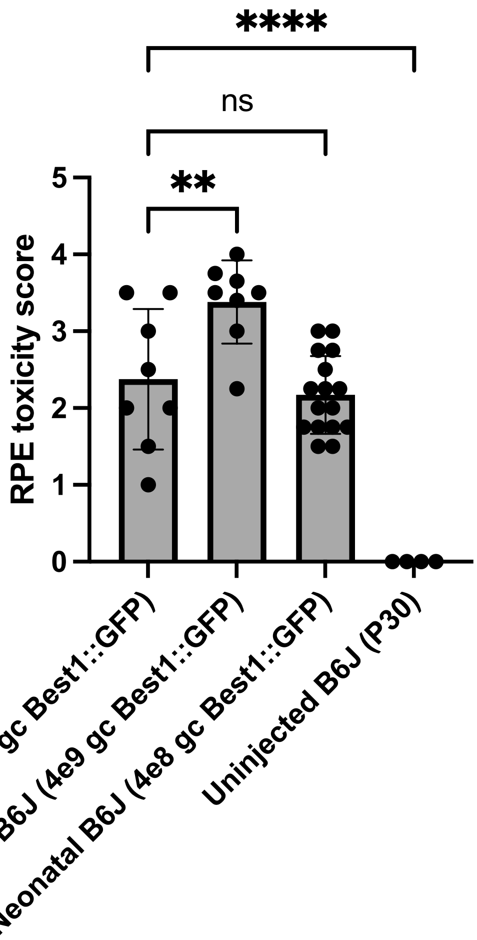


C

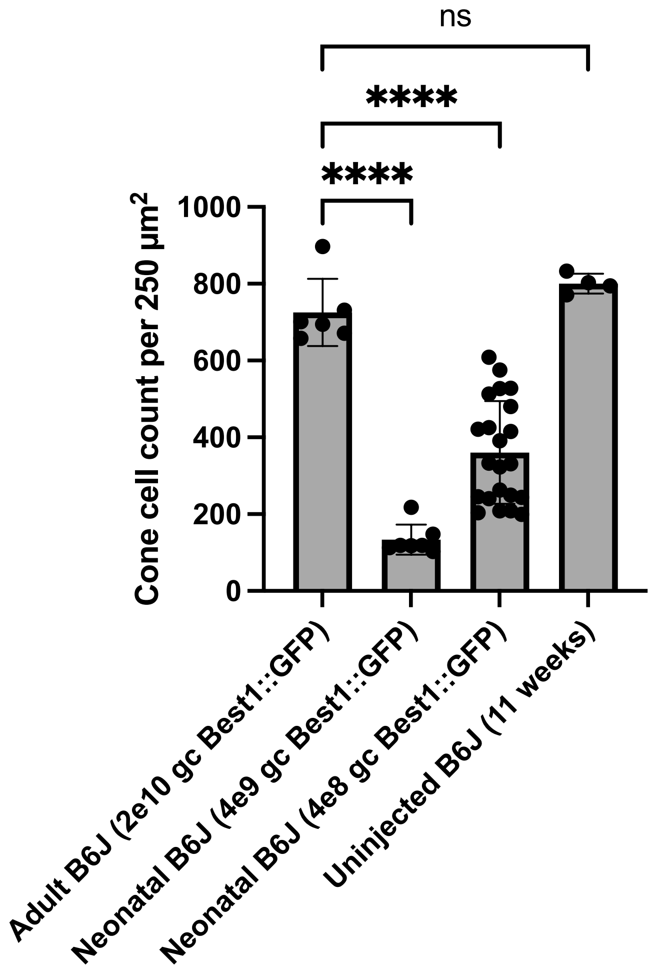

### Figure S17

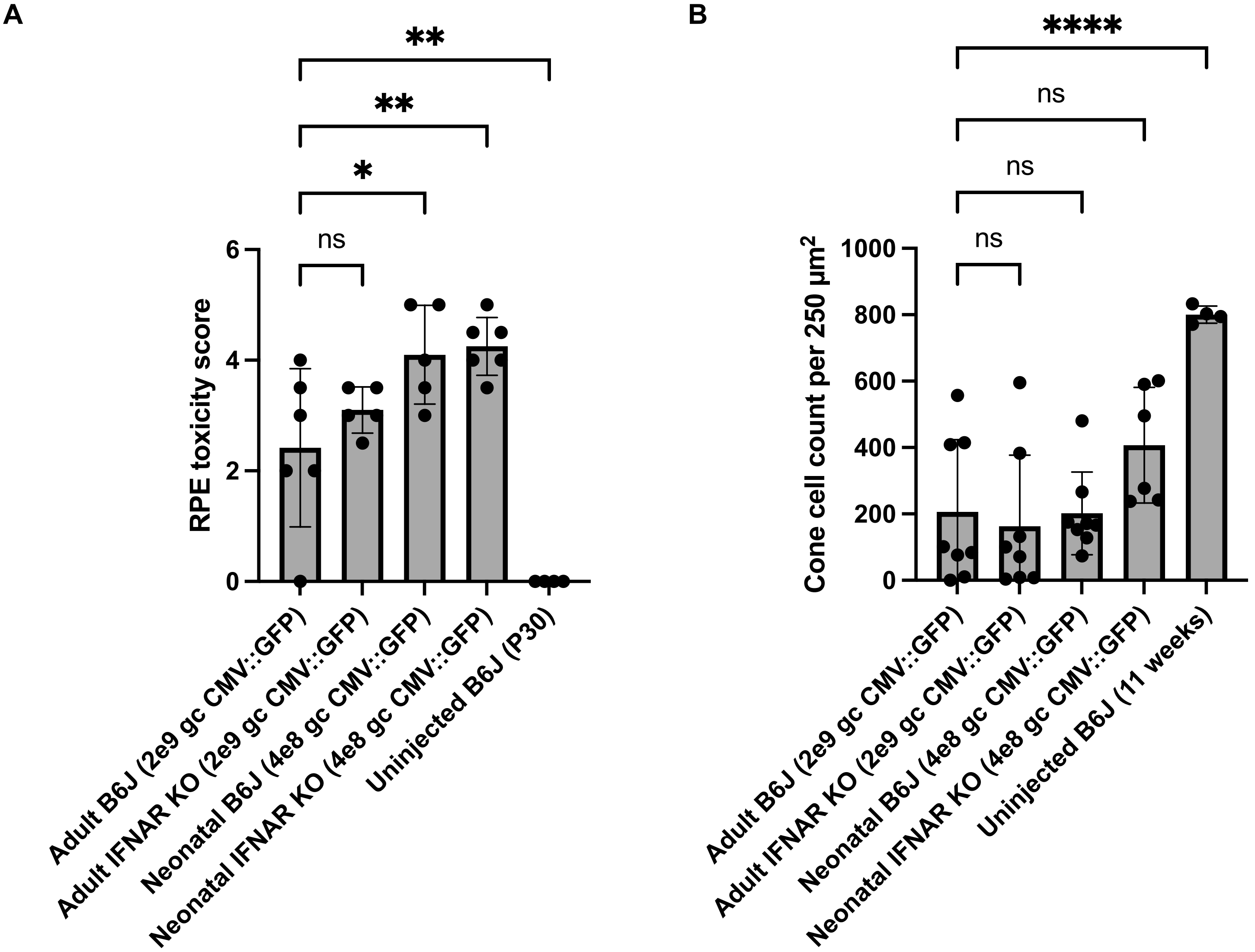
A
**
**
*
ns
6
4
2
0
RPE toxicity score
Adult B6J (2e9 gc CMV::GFP)
Adult IFNAR KO (2e9 gc CMV::GFP)
Neonatal B6J (4e8 gc CMV::GFP)
Neonatal IFNAR KO (4e8 gc CMV::GFP)
Uninjected B6J (P30)
B
****
ns
ns
ns
1000
800
600
400
200
0
Cone cell count per 250 µm²
Adult B6J (2e9 gc CMV::GFP)
Adult IFNAR KO (2e9 gc CMV::GFP)
Neonatal B6J (4e8 gc CMV::GFP)
Neonatal IFNAR KO (4e8 gc CMV::GFP)
Uninjected B6J (11 weeks)

### Figure S18

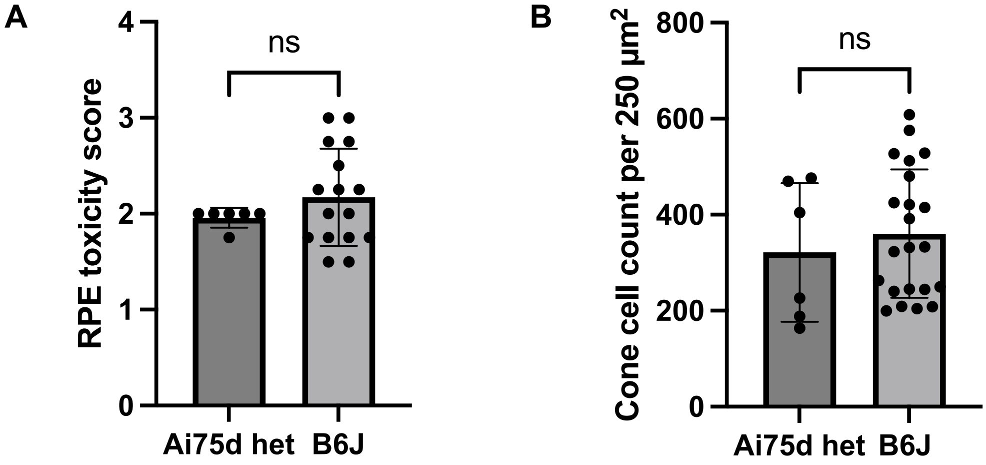
A
RPE toxicity score
ns
0
1
2
3
4
Ai75d het
B6J
B
Cone cell count per 250 µm²
ns
0
200
400
600
800
Ai75d het
B6J
